# Quantifying cell transitions in *C. elegans* with data-fitted landscape models

**DOI:** 10.1101/2021.01.22.426019

**Authors:** Elena Camacho-Aguilar, Aryeh Warmflash, David A. Rand

## Abstract

Increasing interest has emerged in new mathematical approaches that simplify the study of complex differentiation processes by formalizing Waddington’s landscape metaphor. However, a rational method to build these landscape models remains an open problem. Building on pioneering work by ***Corson and Siggia*** (***2012***, 2017) we study vulval development in *C. elegans* by developing a framework based on Catastrophe Theory (CT) and approximate Bayesian computation (ABC) to build data-fitted landscape models. We first identify the candidate qualitative landscapes, and then use CT to build the simplest model consistent with the data, which we quantitatively fit using ABC. The resulting model suggests that the underlying mechanism is a quantifiable two-step decision controlled by EGF and Notch-Delta signals, where a non-vulval/vulval decision is followed by a bistable transition to the two vulval states. This new model fits a broad set of data and makes several novel predictions.

## Introduction

A key stage in the development of an organism is cell differentiation, in which unspecialized cells, called stem cells, become specialized ones depending on the signals that they receive. This is controlled by a very large network of interacting genes (***Davidson, 2010***), the state of which defines the characteristics of the cell. However, this process is still not completely understood.

With the recent development of experimental techniques that allow us to obtain detailed quantitative information about the state of cells over time, new data analysis methods and mathematical models are required for the understanding of cell differentiation. A common approach to the modeling of stem cell differentiation is by means of gene regulatory network (GRN) models, which aim to define the possible differentiation states of a cell by its genetic expression profile. However, these models require a large number of parameters and variables which rapidly increases with the size of the network, complicating its analysis. Moreover, the complexity of such models often means they offer little mechanistic insight.

With this in mind, there has recently been an increasing interest in new kinds of mathematical models that formalize Waddington’s epigenetic landscape metaphor without reference to the underlying molecular network (***Corson and Siggia, 2012***; ***Marco et al., 2014***; ***Corson and Siggia, 2017***; ***Corson et al., 2017***). In this picture, a differentiating cell is represented as a marble rolling down a landscape of hills and valleys, encountering decision points between different lineages, and eventually settling in a valley that defines its cell fate. In particular, the models developed in ***Corson and Siggia*** (***2012***, 2017) reason directly on the phenotype, expressed in geometrical terms, focusing on the dynamics of the general process rather than on the deep molecular scale. These models contain the essence of the process that is necessary for its understanding, which is the structure of cell fates in the Waddington’s landscape and the effect that inducing signals have upon this landscape. In their formulation, the relevant dynamical systems are gradient-like, and the system’s trajectories, which represent the developmental path of a cell, move downhill in this landscape until they reach a minimum representing the cell’s state. Variations in the signals received by the cell change the underlying control parameters of the lanscape and cause bifurcations to occur, allowing the cell’s state to change by moving to another minimum.

In these previous models a function representing the landscape was formulated in an ad-hoc manner, however, and developing a general method to mathematically construct these landscape models remains an open problem. The key features of the models are the ways in which the attractors appear and disappear, i.e. the allowed bifurcations, and how the biological signals are mapped into the parameters representing the landscape. Catastrophe Theory (CT) and Dynamical Systems Theory provide very powerful tools for classifying the types of bifurcations or singularities that can be present in a gradient-like landscape that, in mathematical terms, can be expressed as a family of potential functions. While CT seemingly only pertains to local bifurcations, used together with other ideas from Dynamical Systems Theory, CT and its key results can be used to gain understanding of the global structure of landscape bifurcations.

Vulval development is an excellent but challenging problem to test this approach. It involves a broadly studied stage in *C. elegans* development, and although there is an extensive amount of data available, it is far from completely understood. The vulva is an adult structure that develops during the larval stages of the hermaphrodite worm. The mature vulva contains 22 cells of different types, and more than 40 genes are involved in its development. It is derived from six ectodermal cells (P3-8.p), called **vulval precursor cells** (VPCs); they are partially differentiated and situated in a row along the antero-posterior axis on the ventral side of the larva, and they all have equivalent developmental potential at the start of the process (***Figure 1***A).

**Figure 1.**
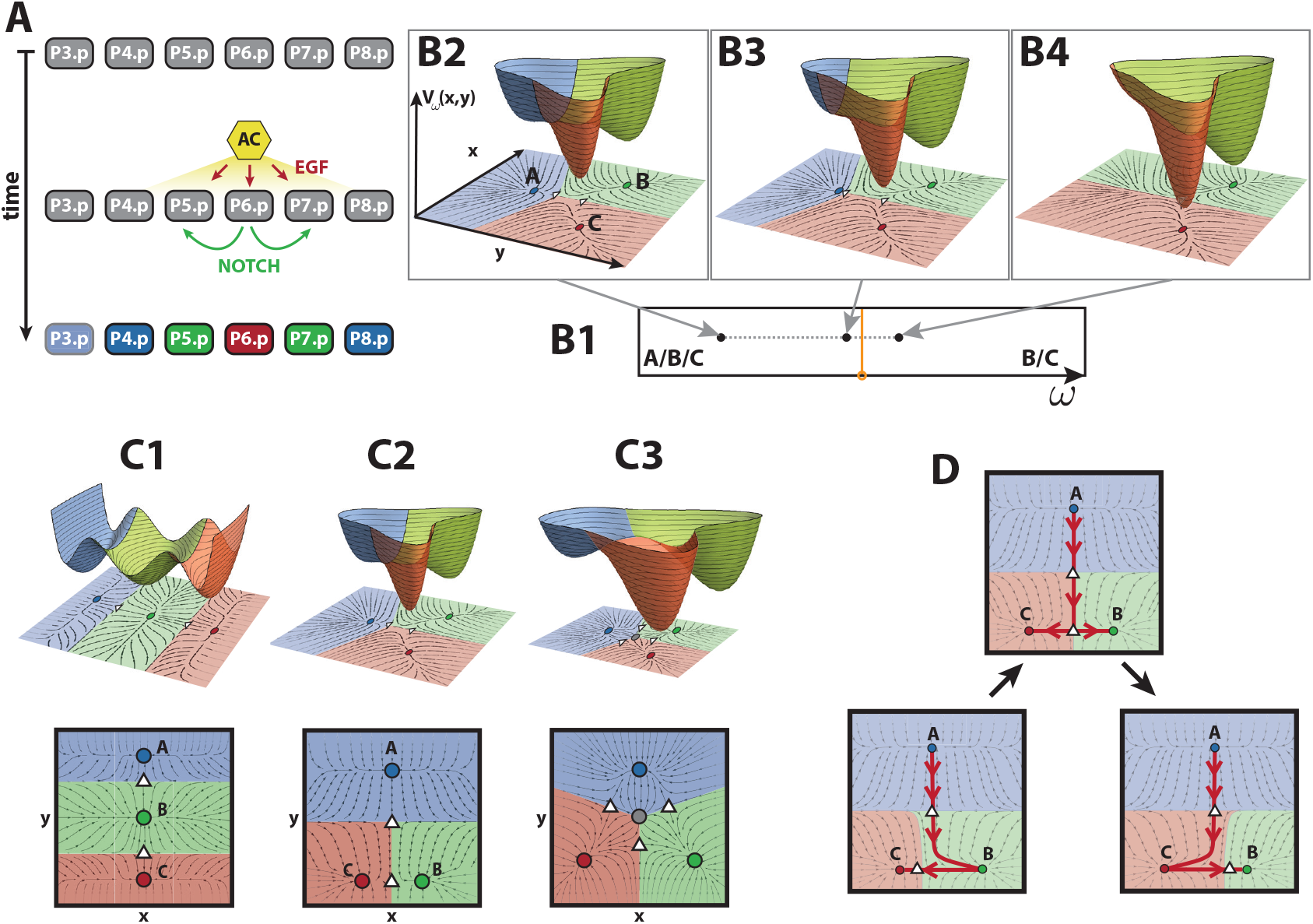
Vulval development in *C. elegans* and landscape models. (A) Schematic representation of vulval development in *C. elegans*. Anchor cell (AC) induces vulval precursor cells (VPCs, P3-8.p) to differentiate into three different cell types: primary fate (red), secondary fate (green) tertiary fate (blue). The pattern is controlled by two signals, EGF from the AC (red arrows) and paracrine Notch (green arrows). VPCs are colored according to the WT pattern. P3.p is colored in a shaded blue because it is not included in our study. (B) Example of a bifurcation. (Top) Example of a landscape with three attractors over the corresponding 2-dimensional flow, attractors and saddles (top). Attractors and their corresponding basins of attraction are circles colored accordingly. (Bottom) Fate map defined by the parameter *ω*, where the bifurcation set is colored in orange. (B2) For this value of *ω*, the landscape contains three attractors *A, B* and *C*. (B3) A change in the parameter of the potential that is now closer to the bifurcation set produces a change in the stability of the landscape, making the valley corresponding to the blue attractor shallower as the attractor approaches the saddle. (B4) The parameter value has crossed the bifurcation set and is now positioned in another region of the fate map. The bifurcation has happened: the blue attractor has bifurcated away and is no longer present in the landscape. (C) The three possible landscape topologies for a process involving three fates, ordered by simplicity. Basins of attraction and attractors are colored according to the fate they could represent. Saddles and repellers are represented as white triangles and gray circles, respectively. (C1) Thom’s butterfly catastrophe where the three attractors representing the three different cell types are positioned on a curve. This topology corresponds to the morphogen model and is inconsistent with AC ablation data in ***Table 1***. (C2) 2-dimensional landscape with two decision points: vulval vs non-vulval fate and primary vs secondary fate. This is the binary flip with cusp landscape and is our choice of landscape. (C3) Symmetric 2-dimensional landscape where all transitions are allowed. This is the landscape studied in ***Corson and Siggia*** (***2012***, 2017) (D) Schematics of the transitions allowed in the binary flip with cusp model. The unstable manifold of the top saddle can flip from attractor B (bottom left) to attractor C (bottom right) by going through an intermediate state where the unstable manifold of the top saddle directs the flow to the bottom saddle (middle top). This intermediate state is called a heteroclinic connection.

During vulval development, the VPCs can develop into three different fates: primary (1°), secondary (2°) and tertiary (3°). The first two fates correspond to vulval cells, while the tertiary fate is non-vulval, and cells adopting the tertiary fate fuse with the large syncytial epidermis hyp7. In the wild type (WT) (i.e. the typical form of *C. elegans* as it occurs in nature), P6.p assumes a primary fate, P5.p and P7.p assume a secondary fate and the rest of VPCs adopt a tertiary fate (***Figure 1***A). This means that only three precursor cells (P5.p, P6.p and P7.p) form the vulva, while the remainder fuse with the syncytial epidermis (***Sternberg, 2005***; ***Lints and Hall, 2009***; ***Wolpert et al., 2015***). P3.p often does not divide, fuses with the hypodermis, and assumes a tertiary fate (***Sternberg and Horvitz, 1986***; ***Sternberg, 2005***; ***Corson and Siggia, 2012***).

This process is orchestrated by the anchor cell (AC), positioned in the gonad of the larva, and VPCs determine their fates through the activation of two signalling pathways (***Schmid and Hajnal, 2015***): the EGF and Notch pathways. The EGF ligand (*lin-3*) is a signalling molecule secreted by the AC; whereas the Notch ligands, which bind to Notch receptors (*lin-12*), are produced by the VPCs themselves (***Figure 1***A). The period of competence for the VPCs to respond to these external signals begins in the L2 larval stage and stops shortly after the first round of division in L3.

The sequence of events that promote this precise pattern of cells fates (in which cells P.4-8 adopt the fates 3°2°1°2°3°), which happens with an accuracy higher than 97% (***Braendle and Félix, 2008***), is still unknown. Two models have been proposed to describe the patterning mechanism: the **morphogen model** and the **sequential model**. In the morphogen model, EGF acts as a morphogen and its levels determine the fate of each cell. Consequently, under this model, the fate of a cell depends on the distance to the AC (***Sternberg and Horvitz, 1986***). On the other hand, the sequential model hypothesizes that the anchor cell induces the primary fate in the closest VPC which, in turn, induces the neighboring cells towards secondary fate through activation of the Notch pathway. Cells is which neither of these pathways are activated adopt a tertiary fate (***Simske and Kirn, 1995***). However, there is experimental evidence supporting characteristics of both of these two mechanisms and a more accurate description likely includes aspects of both of these models.

Phenotypes have been determined for a number of mutant worms with alterations in either Notch or EGF signaling. These experiments shed light on the mechanisms that regulate the development of the vulva. For example, it has been observed that a significant reduction of the EGF signal (either by AC ablation at a very early time or by EGF loss of function) produces the pattern 3°3°3°3°3° (***Komatsu et al., 2008***), suggesting that the activation of the EGF pathway is necessary for the induction of the VPCs.

In their pioneering work, ***Corson and Siggia*** (***2012***, 2017) use a relatively complex model to quantitatively fit a substantial and complex set of experimental results, and use this to provide a number of quantitative predictions. Building on their work, here we focus on vulval development in *C. elegans* to illustrate a new mathematical framework to develop and quantitatively fit landscape models to biological data, based on CT and approximate Bayesian computation (ABC) fitting. Given a few reasonable assumptions, we first argue that CT allows us to rationally identify the candidate landscapes. The elementary catastrophes enumerated by CT are polynomial equations in at most two variables and a few parameters, which simplifies the analysis. The central idea is that we can use these relatively simple dynamical systems as modules to build up a global system with the desired characteristics, which dynamics might not be reproducible by a single polynomial equation. Once this model is developed, we show a novel application of ABC methods and demonstrate how coupling CT with ABC fitting to effciently explore the parameter space provides a useful methodology to fit landscape models to a large amount of data. While the symmetric model developed in ***Corson and Siggia*** (***2012***, 2017) assumed that all cell states are equivalent and cells can transition to any fate at all times, with this approach we show that a simpler model where the the non-vulval fate is distinct from vulval ones and to which cells cannot go back to once they are partially differentiated, is also consistent with the data. Taken together, here we show that we can build a fundamentally different and considerably simpler model to the ones in ***Corson and Siggia*** (***2012***, 2017) that explains the large amount of data available, and makes a number of interesting predictions that suggest novel ways of understanding biological effects and new experiments to validate them.

## Methods

### A three-way decision landscape

Dynamical Systems Theory allows us to frame the Waddington’s landscape idea in mathematical terms. The state of a cell at a particular time *t* will be represented by the position on the landscape, ***x***(*t*) = (*x*(*t*)*, y*(*t*)). The evolution of ***x***(*t*) in time, representing the differentiation of a cell, will be related to the gradient or steepness of the landscape represented by a potential function *V* (***x***), and will move downhill in this landscape until it reaches a local minimum (an attractor). Each attractor of *V* (***x***) corresponds to a stable fate that a cell can adopt. Given an attractor ***x***\*, its basin of attraction is comprised of all the points with a trajectory that ends in that given attractor. Finally, the lowest point of the barriers that separate these basins are saddle points of the potential function (***Figure 1***B).

Now, in order for the trajectory to move from one attractor to another one, the barrier between them needs to disappear. The induction of a new fate is biologically controlled by cues, known as morphogens or signals, that the cell receives and produce a change in its gene expression, pushing the cell towards a new differentiated state. Translating this idea to mathematical terms, the shape of the landscape must be controlled by these signals and its shape will change as these signals change in time. Therefore *V* (***x***) also depends on some parameters *ω*, which will relate to the biological signals, becoming the family of functions *V* (***x***, *ω*) = *V_ω_*(***x***).

The most important changes in the landscape are the ones in which an attractor is created or destroyed as the parameters change, and these are called bifurcations. These normally happen when an attractor collides with a nearby saddle. An example of this is shown in ***Figure 1***B. From left to right, by changing the parameter *ω* of the landscape, the basin of attraction corresponding to the blue attractor, A, gradually becomes flatter as the attractor approaches the saddle point. These two points eventually merge together and disappear. This bifurcation would shift the trajectory of a cell starting in the blue attractor towards either the red or green one. The values of the parameters at which bifurcations occur form the **bifurcation set** of the potential function *V*. The bifurcation set of a potential defines regions in the parameter set that correspond to different stability configurations in the landscape, or fate map (***Figure 1***B).

#### Choice of landscape

The problem now is to find the simplest possible parameterized family of landscapes that can quantitatively reproduce the substantial set of experimental results summarized in ***Table 1***. A good starting point is to note that some of the landscapes in this family must contain three attractors as, during the process of *C. elegans* vulval development, a VPC must be able to adopt one of the three fates introduced above: primary, secondary and tertiary. If we demand simplicity by not permitting any repelling points in the landscape, there are essentially only two landscapes with three attractors (***Figure 1***C1-2). An example of a more complex landscape with a repeller is shown in ***Figure 1***C3, other, even more complex examples, also exist.

**Table 1.** Table of experimental data obtained from the literature (***Corson and Siggia, 2017***).

| Experiment |  | VPC fates (% 1°, 2°, 3°) |  |  | TD | VD | References |
| --- | --- | --- | --- | --- | --- | --- | --- |
|  |  | P4.p | P5.p | P6.p |  |  |  |
| Fully penetrant phenotypes |  |  |  |  |  |  |  |
| (1) | Wild type | 3° | 2° | 1° | × |  |  |
| (2) | *let-23* mosaic<br>(no EGF receptors in P5/7.p) |  | Wild type |  | × |  | (Koga and Ohshima (1995);<br>Simske and Kirn (1995)) |
| (3) | Half dose of *lin-3*<br>(Half EGF ligand) |  | Wild type |  | × |  | (Ferguson and Horvitz, 1985) |
| (4) | Half dose of *lin-12*<br>(Half Notch receptor) |  | Wild type |  | × |  | (Greenwald et al., 1983) |
| (5) | Notch null, and 2xWT EGF (2 ACs) | 3° | 1° | 1° | × |  | (Greenwald et al., 1983) |
| (6) | No Notch signalling, WT EGF | 3° | 3° | 1° | × |  | (Sundaram and Greenwald (1993);<br>Shaye and Greenwald (2002);<br>Komatsu et al. (2008)) |
| Excess EGF |  |  |  |  |  |  |  |
| (7) | JU1100 (overexpression of EGF ligand, level to fit) | 18,46,36 | 45.5,54.5,0 | 96,4,0 | × |  | (Hoyos et al., 2011) |
| (8) | JU1107 (2.75xWT EGF) | 2,15,83 | 19,81,0 | 100,0,0 |  | × | (Barkoulas et al., 2013) |
| Reduced Notch |  |  |  |  |  |  |  |
| (9) | JU2039 (WT EGF, reduced Notch) | 0,0,100 | 1,89,10 | 100,0,0 |  | × | (Barkoulas et al., 2013) |
| (10) | JU2113 (1.25xWT EGF, reduced Notch) | 4,6,90 | 18,70,12 | 100,0,0 |  | × |  |
| Excess EGF & ectopic Notch |  |  |  |  |  |  |  |
| (11) | JU2091 (1.25xWT EGF, ectopic Notch) | 0,0,100 | 0,100,0 | 100,0,0 |  | × | (Barkoulas et al., 2013) |
| (12) | JU2089 (1.79xWT EGF, ectopic Notch) | 0,6,94 | 1,99,0 | 100,0,0 |  | × |  |
| (13) | JU2092 (2.75xWT EGF, ectopic Notch) | 5,24,71 | 0,100,0 | 100,0,0 |  | × |  |
| Reduced EGF |  |  |  |  |  |  |  |
| (14) | CB1417 (*lin-3*(e1417) EGF hypomorph) | 0,0,100 | 1,99,0 | 54,0,46 |  | × | (Barkoulas et al., 2013) |
| (15) | JU2095 (mild ectopic Notch × EGF hypomorph) | 0,1,99 | 0,15,85 | 72,0,28 |  | × |  |
| Phenotypes following anchor cell ablation |  |  |  |  |  |  |  |
| (16) | L2 lethargus | - | 0,0,100 | 0,0,100 | × |  | (Milloz et al., 2008) |
| (17) | Early L3 | - | 1.5,21,77.5 | 18,18,64 |  | × |  |
| (18) | DU divided | - | 0,54.63,45.37 | 31,38,31 | × |  |  |
| (19) | VU divided | - | 4,90,6 | 52,48,0 |  | × |  |
| (20) | 3° divided | - | 1,99,0 | 65,35,0 |  | × |  |
| (21) | 2-cell stage | - | 1,99,0 | 93,7,0 |  | × |  |

Fates for P4.p and P8.p (or P5.p and P7.p) have been averaged, since we assume that the pattern is symmetrical around the AC, and P3.p is not considered in this study as it often does not divide and fuses with the hypodermis, assuming tertiary fate. Developmental stages: (L2 lethargus) lethargic L2; (Early L3) early L3; (DU divided) Dorsal Uterine precursor cells dividing or divided once; (VU divided) Ventral Uterine precursor cells dividing or divided once; (3° divided) 3° cells have divided; (2-cell stage) all Pn.p cells have divided once.

As discussed above, cell state transitions occur because changes in the signals a cell receives destabilize the attractor where the cell state lies allowing it to transition to a new one. In considering these landscapes it is important to keep in mind that since the attractors correspond to biological cell states, there must be a clear correspondence between each attractor and a particular state, and this relationship must be tracked as the attractors move due to signal changes. We label them here *A*, *B* and *C*. The first landscape (***Figure 1***C1), which is related to Thom’s butterfly catastrophe (***Poston and Stewart, 2014***), differs from the second (***Figure 1***C2) in that, with regard to decision-making, it is more restrained. It is characterized by the fact that, in the parameter region where there are three attractors, the identity of the central state *B* cannot change. The only transitions allowed are *A* → *B* (by bifurcating *A* with the top saddle), *B* → *A* (by bifurcating *B* with the top saddle), *B* → *C* (by bifurcating *B* with the bottom saddle) and *C* → *B* (by bifurcating *B* with the bottom saddle). This means that cells need to pass through state *B* in order to go from *A* to *C*. This landscape, in fact, corresponds to the morphogen model, in which a VPC would stay in tertiary fate (*A*) under a low EGF signal, would transition from tertiary fate to secondary fate (*A* → *B*) in response to a middle EGF signal, and to primary fate (*A* → *B* → *C*) in response to a high EGF signal. This landscape, however, is not consistent with the AC ablation experiments in ***Milloz et al.*** (***2008***), where early AC ablation experiments show an equal escape of cells to secondary and primary fates (***Table 1***) at early AC ablation times, since this means that cells take the same time to go from *A* to *B* than from *A* to *C*.

On the other hand, for the alternative landscape (***Figure 1***C2), the fact that the unstable manifold of the top saddle *A* can flip from *B* to *C* via a heteroclinic connection allows extra transitions and for either of *B* and *C* to be the central attractor (***Figure 1***D). Therefore, in this case, the transition *A* → *C* is also allowed and cells can go to the state *C* without passing through *B*, which is consistent with the AC ablation experiments mentioned earlier. We call the landscape in ***Figure 1***C2 the **binary lip with cusp landscape**.

The more complex landscape containing a repeller shown in ***Figure 1***C3 is essentially that used in ***Corson and Siggia*** (***2012***, 2017), where it was shown that it can reproduce the data we are concerned with. In this model, all attractors are equivalent to each other and all bifurcations are allowed. Biologically, this means that all cell states are equivalent and cells can transition to any fate at all times.

Our aim is to show that the same data set can also be explained by the considerably simpler binary flip with cusp landscape. This results in a different biological interpretation of the decision-making process. In this picture, not all the fates are equivalent, but the tertiary fate is a special fate. It is the default state for a cell if it does not receive any signal; and cells do not return to it once partially induced, even if the signals are switched off (as AC ablation experiments show in ***Table 1*** (***Milloz et al., 2008***)). Transitions from fate 3° to fate 1°, and fate 3° to fate 2°, are allowed (***Sternberg and Horvitz, 1989***). Also, the transitions from fate 2° to fate 1° is achieved by increasing the EGF signal and turning Notch signal off (***Greenwald et al., 1983***). These reasons lead us to hypothesize that the topology of decisions is represented by that of the binary flip with cusp landscape. First, cells will decide whether to differentiate into a vulval fate or stay non-vulval (determined by a bifurcation between the blue attractor and the top saddle in ***Figure 1***C2) and, if they do so and surpass the top saddle, then they will decide whether to become 1° or 2° fated cells (determined by bifurcations between the bottom critical points in ***Figure 1***C2).

#### Building the landscape

In order to find a representative model for this landscape, we could attempt to use CT to search for a parameterised set of functions such that the parameterised landscape is given by the gradient flows of these functions. However, this approach is not practicable for very complex landscapes. Therefore we employ an alternative method that can be used in this more general context. We use catastrophe and bifurcation theory to analyse the local structure and then use ideas from differential topology to glue the local models together. In our case this involves composing two simple catastrophes: a fold (commonly also known as a saddle-node bifurcation) that will explain the non-vulval vs vulval bifurcation (bifurcation between attractor *A* and top saddle in ***Figure 1***C2), and a cusp catastrophe that will explain the primary vs secondary fate bifurcation (bifurcation between attractors *B* and *C* and bottom saddle in ***Figure 1***C2). For gradient and gradient-like systems these are the only local bifurcations that occur generically and are universal in one and two-parameter systems (***Guckenheimer and Holmes, 2013***).

The fold or saddle-node is the only bifurcation that appears generically in one-parameter gradient-like systems and it has the normal form 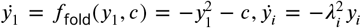 (***Figure 2***A). Any system displaying a saddle-node bifurcation can be transformed into this (***Guckenheimer and Holmes, 2013***). Here, when the parameter *c* < 0, the system has two equilibria, an attractor and a saddle, and when *c* > 0 it has none (see ***Appendix 1*** for more details). Thus, it can represent the disappearance of the tertiary fate upon receiving signals. The distance between these two equilibria grows like 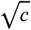. We will use it to define the attractors corresponding to 3° fate and the point of transition between this non-vulval state and the other two vulval states (bifurcation between 3° attractor and top saddle in ***Figure 2***C).

**Figure 2.**
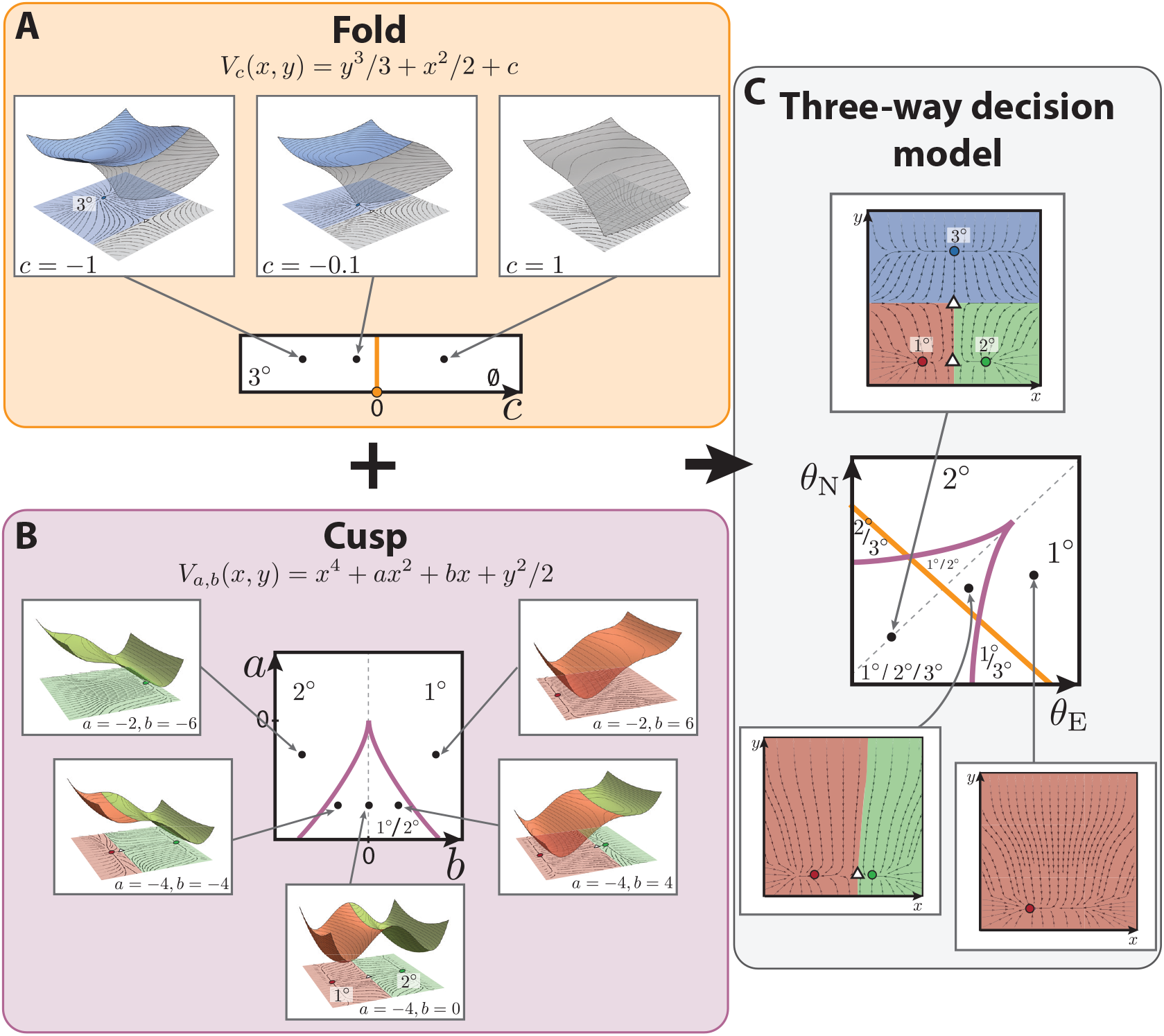
Binary lip with cusp model of vulval development in *C. elegans*. (A) The fold catastrophe, controlled by the parameter *c*, defines the stability of the attractor representing 3° fate. The fate map is represented at the bottom, showing that the fold has a bifurcation at *c* =0 (in orange), separating two stability regions: one containing 3° fate (for *c* < 0) and one where the fate disappears (*c* > 0). At the top, from left to right, different examples of landscapes for different values of *c*. For *c* < 0, the potential has two critical points in the state space: a minimum (blue circle) and a maximum (white triangle), such as the one represented on the top panels. Close to the bifurcation point, *c* = 0, the two critical points get near each other to merge together. For values of *c* > 0, the potential does not contain any critical point. (B) The cusp catastrophe, controlled by the parameters *a, b* defines the stability of the attractors representing 1° and 2° fates. The fate map, at the center, shows that the cusp has a bifurcation set determined by Δ = 8*a*^3^ + 27*b*^2^ =0 (purple cusp), separating three stability regions: one containing 1° and 2° fate (for Δ < 0) and two where only one of the two fates is present (Δ > 0). Examples of landscapes determined by different values of *a, b* are shown. On the cusp mid-line (gray dotted line) the two fates are equally probable. Getting closer to the cusp favors one fate over the other, by the saddle approaching one of the attractors. Outside the cusp, the landscape only has one attractor, either 1° or 2°. (C) The two catastrophes are combined to generate a 2-dimensional flow with two saddles (white triangles) and three attractors representing each fate (blue, green and red circles represent 3°, 2° and 1° fates, respectively). By mapping the signals *θ*_E_, *0*_E_ onto the control parameters *a, b, c*, a fate map is obtained, defining the available fates in the landscape for different signaling profiles. The presence of 3° fate is controlled by the fold line, as shown in (A), and the stability of the 1° and 2° fates is controlled by the position of the signaling profile with respect to the cusp line, as shown in (B).

For 2-parameter gradient-like systems the only other generic bifurcation is the cusp. The relevant normal form is 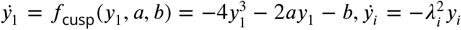. In this case, when the discriminant Δ = 8*a*^3^ + 27*b*^2^ < 0, the normal form system contains two minima and a maximum; when Δ = 0, it contains one minimum and a degenerate point; and if Δ > 0, it contains just one attractor (see ***Figure 2***B and ***Appendix 1*** for more details). We will use it to define the attractors corresponding to fates 1° and 2° and the saddle between them (***Figure 2***B-C).

We merge the fold and the cusp catastrophe models into a two-dimensional dynamical system, the mathematical details of which are in ***Box 1*** and ***Appendix 1*** (***Figure 2***C).

The steady states of this system will lie either on the *x*-axis, in which case their *x*-coordinates will be given by the steady states of the cusp (i.e. there will be three, two or one steady states on the *x*-axis); or they will have *y*-coordinates which are the zeroes of a fold, (there will be two, one or none depending on *c*) in which case their *x*-coordinates will be 0 (see ***Box 1***, ***Figure 2***C and ***Appendix 1*** for more details).

We choose the attractor with zero *x*-coordinate to represent the 3° fate, the attractor with zero *y*-coordinate and negative *x*-coordinate to represent the 1° fate and the attractor with zero *y*-coordinate and positive *x*-coordinate to represent 2°, as shown in ***Figure 2***A. Taken together, parameter *c* will control whether 3° fate is present in the landscape, and *a, b* will control whether the attractors representing 1° and 2° fates will be present in the landscape.

The advantage of building the landscape from these two catastrophes is that, firstly, mathematically one has the universality property mentioned above and further described in ***Appendix 1*** and, secondly, that the position and stability of the various restpoints and their dependence on the parameters is transparent and, as shown in ***Box 1***, one has complete understanding of the regions of the parameter space with common stability, being able to characterise the shape of the lanscape for any parameter value.

#### Dependence upon the morphogens

The vulval development process is controlled by induction of the VPCs by the anchor cell via the EGF signal and by lateral signalling through Notch. Therefore, to connect our model to the biological data we need to define how the control parameters *a*, *b* and *c* defined previously depend upon the levels *θ*_E_ and *θ*_N_ of the EGF and Notch signals, respectively.

As we describe in ***Box 2***, in more detail, we choose a flexible linear dependence of the control parameters upon the biological signals.

The transformation can be constrained by noting the following facts:

1. The origin of the (*θ*_E_, *θ*_N_) coordinate system is assumed to correspond to a point in which there is tristability in the landscape, as once induced all three fates are stable after the removal of signals (see ***Table 1***) (***Sternberg, 2005***).
2. The signals EGF and Notch drive the system out of the tristable region.
3. The Notch null, 2×WT EGF perturbation results in an induction of P5.p and P6.p cells to adopt primary fate (see 1), and therefore this signaling profile lies outside the tristable region.
4. Similarly for the Notch null with WT EGF perturbation, P6.p cell adopt primary fate and therefore its signaling profile lies outside the tristable region.

###### Box 1. Mathematical details of the model

We merge the fold and the cusp catastrophes into the following model, which will be used to model the evolution of the state of a single VPC in time, represented by ***x***(*t*)= (*x*(*t*)*, y*(*t*)):

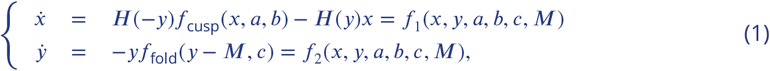

where *H* is a Heaviside function that is equal to 0, if *y* is less or equal than 0, and equal to 1 otherwise. Strictly speaking to get a smooth dynamical system we should smooth the Heaviside function but, as will become clear and is shown in ***Appendix 1***, this does not affect the analysis shown here.

The attractors of this system are obtained by solving *f*_1_ = *f*_2_ = 0. Since *f*_2_ does not depend on *x*, one can quickly solve it to obtain the *y*-coordinates of the critical points, which are given by 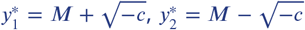 and 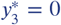. If *f*_1_ =0 is now solved with each of the possible values of *y* obtained, the *x*-coordinate of the critical points with 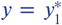 or 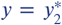 is equal to 0; while the *x*-coordinates of the critical points with 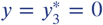 are given by the zeros of the cusp, i.e. *f*_cusp_(*x*\**, a, b*)= 0. The parameter *M* controls the relative position of top and bottom saddles in ***Figure 2***C.

The stability of these points can be obtained by looking at the Jacobian matrix of the system. It can be shown that the point 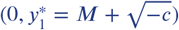 is stable (blue attractor in ***Figure 2***C, assigned to represent 3° fate), the point 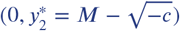 is unstable (white top triangle in ***Figure 2***C), while the points laying on the *x*-axis can be either two stable points (representing 1° and 2° fates) separated by a saddle (as in ***Figure 2***C), a stable point and a degenerate point or just a stable point. We refer the reader to ***Appendix 1*** for more details and Appendix 1 Figure 4 for more examples of landscape configurations.

Taken together, the parameter *c* controls the existence and positions on the *y*-axis of critical points with positive *y* coordinate (which can be two, one or none), while the parameters *a, b* control the the positions on the *x*-axis and existence of critical points with zero *y*-coordinate (which can be three, two or one). With this model, 3° attractor can be bifurcated away and be removed from the landscape, however, either 1° or 2° fates need to be present and both cannot be bifurcated away to give a landscape where only 3° fate is present, which is intrinsically different from the model in ***Corson and Siggia*** (***2012***, 2017).

Since the 3° attractor is controlled by the fold we know that if *c* < 0, the attractor corresponding to this fate will be present, no matter the values of *a* and *b*. On the other hand, the presence of the attractors corresponding to 1° and 2° fates depends only on the values of *a* and *b*, specifically on the value of the discriminant Δ = 8*a*^3^ + 27*b*^2^ as explained above, independently from the value of the parameter *c*, and therefore their presence will be controlled by the value of Δ.

One can then compute the bifurcation set of the landscape in Equation 1, and obtain:

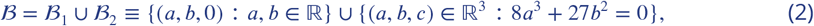

where 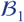 and 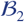 are represented in orange and purple colors, respectively, in ***Figure 2*** and Appendix 1 Figure 7. Bifurcations will happen at and only at the values of the parameters in 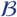. This bifurcation set divides the control space into regions with common stability, some of which are represented in Appendix 1 Figure 7. More details can be found in ***Appendix 1***.

###### Box 2. Mathematical details of the map between control parameters and signals

We will postulate a flexible functional form for the relationship between the control parameters *a*, *b* and *c* and the signaling levels *θ*_E_ and *θ*_N_, and then fit this to the data. We assume that the parameters *a*, *b* and *c* are affine functions of *θ*_E_ and *θ*_N_ so that

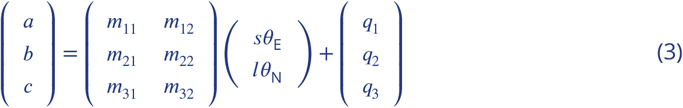

where *m_i,j_* < 1 and *s, l* are scaling parameters of the signals into the control space. Then (*a, b, c*) lies on the plane *π_T_* given by *Aa* + *Bb* + *Cc* = *D* where *A* = *m*_31_*m*_22_ − *m*_21_*m*_32_, *B* = *m*_11_*m*_32_ − *m*_31_*m*_12_, *C* = *m*_12_*m*_21_ − *m*_11_*m*_22_, and *D* = −*Aq*_1_ − *Bq*_2_ − *Cq*_3_

This plane intersects the bifurcation set 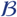 in different subsets depending on the values of the *m_ij_* and the scaling parameters *s, l*, while the origin of the (*θ*_E_, *θ*_N_) coordinate space in that plane will be determined by the parameters *q_i_*. Consequently, the bifurcations can be visualized in the (*θ*_E_, *θ*_N_)-signal space by intersecting the plane and the bifurcation set 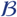 and this gives a fate map for the cell as a function of the signals EGF and Notch (See ***Appendix 1*** for more details).

#### More details can be found in *Appendix 1*

Next, to complete our model, we need to define the signaling profile of each cell in time, which we elaborate in ***Box 3***. As we lack dynamic signaling data, such as could be obtained from live-cell reporters, we make the following assumptions. First, to model the EGF signal, 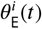, received by a cell P*i*.p, we consider that the EGF ligand is secreted by the AC, and therefore the signal received by a cell will depend on its proximity to the AC. We assume this signal is not instantaneous, so its increase is modeled as a monotonously increasing function in time. This is consistent with data published in ***Milloz et al.*** (***2008***), where the activity of the EGF pathway was measured using a transcriptional reporter, and showed a monotonal increase in time, as shown in ***Appendix 1***. Second, to model 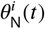, the Notch signal received by a cell P*i*.p, it is known that Notch signal production is a consequence of EGF pathway activation, so that the Notch signal is produced by cells receiving high EGF as they adopt the primary fate. Therefore, we assume cells produce Notch signals as they approach the basin of attraction corresponding to the 1° fate. Moreover, the Notch signal received by a cell is a sum of autocrine Notch via diffusible delta ligands plus paracrine Notch from the neighboring cells. Taking this together, we model 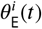 and 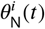 as described in ***Box 3*** and ***Appendix 1***, in more detail.

Finally, to account for variability in the outcomes, we add some random white noise in the dynamics of each cell, which is parameterised by a coeffcient of diffusion in the phase space.

If we now combine the model in ***Box 1*** together with the mapping from the signals to the control parameters in ***Box 2*** and the signalling dynamics to which a cell is exposed in time, described in ***Box 3***, we obtain the proposed model.

Each cell will start its trajectory in the basin of attraction representing the tertiary fate and will move in its own dynamic flow determined by Eq. 1, the shape of which depends, by the relationship in Eq. 3, on the signals it is receiving.

After the period of competence, the fate of a cell is defined by the basin of attraction at which it ended up. The mathematical details of how this is done are presented in ***Appendix 1***. This now allows us to fit the model to the experimental data.

###### Box 3. Mathematical details of the signaling dynamics

Finally, we model the signal profile of each cell in time in the following way. To reduce unnecessary dimensions, we only model the cells P4− 6.p as the pattern is generally symmetric around the AC and, as mentioned in the introduction, P3.p often assumes tertiary fate. Following the approach in ***Corson and Siggia*** (***2012***, 2017), we assume that the difference of EGF signal between consecutive cells is regulated by a scaling parameter *γ* ≤ 1, which derivation follows from the diffusion of EGF ligand. However, we also model the increase of EGF signal in time as a monotonous increasing function *σ*(*t*):

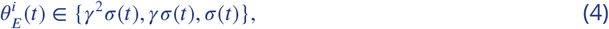

where we write *σ*(*t*) as

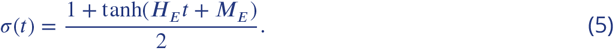

Regarding Notch signaling, following ***Corson and Siggia*** (***2012***, 2017), we define the production of Notch by a cell by a sigmoidal function *L*, that depends on the current state ***x***(*t*) of the cell:

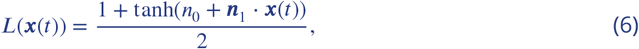

where *n*_0_ and ||***n***_1_|| are constants and the vector ***n***_1_ has negative *x*-coordinate so that *L* increases as the state of the cell approaches the basin of attraction of 1° fate. We now define 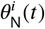 of a cell P*i*.p to be proportional to the sum of the autocrine and the paracrine Notch signal received by the cell. Also following ***Corson and Siggia*** (***2012***, 2017), the autocrine signal is scaled by a parameter *α* > 0 which parametrises the relative importance of autocrine vs paracrine signaling. We also take into account that 1° fated cells downregulate Notch receptor lin-12 (***Shaye and Greenwald, 2002***). This downregulation is related to the cell’s production of Notch signalling, scaled by a parameter *l_d_*, which controls the strength of such downregulation. Therefore the Notch signal received by a cell P*i*.p is modeled as.

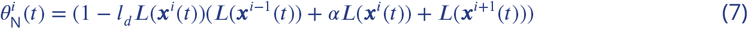

### Parameter estimation: ABC SMC implementation

Now that a parameterised model has been developed, the next step is to find whether there are parameters that allows the model to reproduce the experimental data in ***Table 1*** and to determine the extent of such parameters. ***Table 1*** contains the probabilities of each VPC (P4.p, P5.p, P6.p) acquiring each fate (1°, 2°, 3°) under twenty one experimental conditions (i.e. 171 data points or probabilities in total). From this data set, we will use a subset of nine experimental conditions as training data set for the model to fit, and the remaining ten as validation results (TD and VD in ***Table 1***, respectively).

Let us denote by *p_e,f,c_* the experimental probability of cell *c* becoming fate *f* under experimental condition *e*, the values of which are given in ***Table 1***. Similarly, 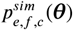 represents the simulated probability of cell *c* becoming fate *f* under experimental condition *e* given parameters ***θ***. Taking a Bayesian approach, we treat the parameters of the model as random variables with a probability distribution. Our goal is to estimate the posterior distribution of the parameters that accurately reproduce the training data set 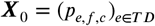, and check that they also reproduce the validation data set.

If we denote a parameter vector by ***θ***, the posterior distribution satisfies:

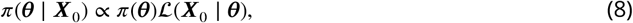

where *π*(***θ***) is the prior distribution of the parameters ***θ***, 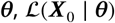 is the likelihood of the data ***X***_0_ given the parameters ***θ*** and *π*(***θ*** | ***X***_0_) is the posterior distribution of the parameters ***θ*** given the data ***X***_0_.

Since it is not possible to find an analytical expression for the likelihood of the model we propose, we approximate it using approximate Bayesian computation (ABC). ABC methods, also known as likelihood-free methods, have been developed for inferring the posterior distribution of the parameters when the likelihood function is either too complex or too expensive to compute but observations of the model can be simulated fairly easily. They have been successfully applied to a wide range of intractable likelihood problems over the past twenty years (***Marin et al., 2012***) including population genetics (***Beaumont et al., 2009***), pathogen transmission (***Tanaka et al., 2006***), reaction networks models (***Toni and Stumpf, 2010***; ***Liepe et al., 2014***) and epidemic modelling (***Mckinley et al., 2009***), among others. ABC methods use a comparison between the experimental and simulated data to measure the goodness of fit, instead of using the likelihood function, allowing for more flexibility.

In particular, here we take advantage of Sequential Monte Carlo ABC (ABC SMC) to explore the parameter space and determine the posterior distribution of the parameters. Various ABC SMC algorithms have been proposed in the literature (***Beaumont et al., 2009***; ***Moral et al., 2012***; ***Sisson et al., 2007***; ***Toni et al., 2009***), but here we focus on the fairly general ABC SMC algorithm in ***Toni et al.*** (***2009***). The ABC SMC sampler methodology approximates the posterior distribution by sequentially sampling from a sequence of intermediate distributions, or approximate posterior distributions, {*π_t_*}_0≤*t*≤*T*_, that increasingly resemble the posterior distribution given in Equation 8:

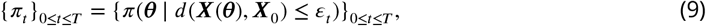

where *d* is a metric that compares the experimental and simulated data, and {*ε_t_*}_0≤*t*≤*T*_ is a decreasing sequence of distance values. This ABC method is particularly useful in cases where the number of parameters and the data are high dimensional, as it can be easily parallelised. It can also be tuned to effciently explore the parameter space maintaining areas of high likelihood via the choice of its perturbation kernel. A summary of how this is done is shown in ***Figure 3***.

**Figure 3.**
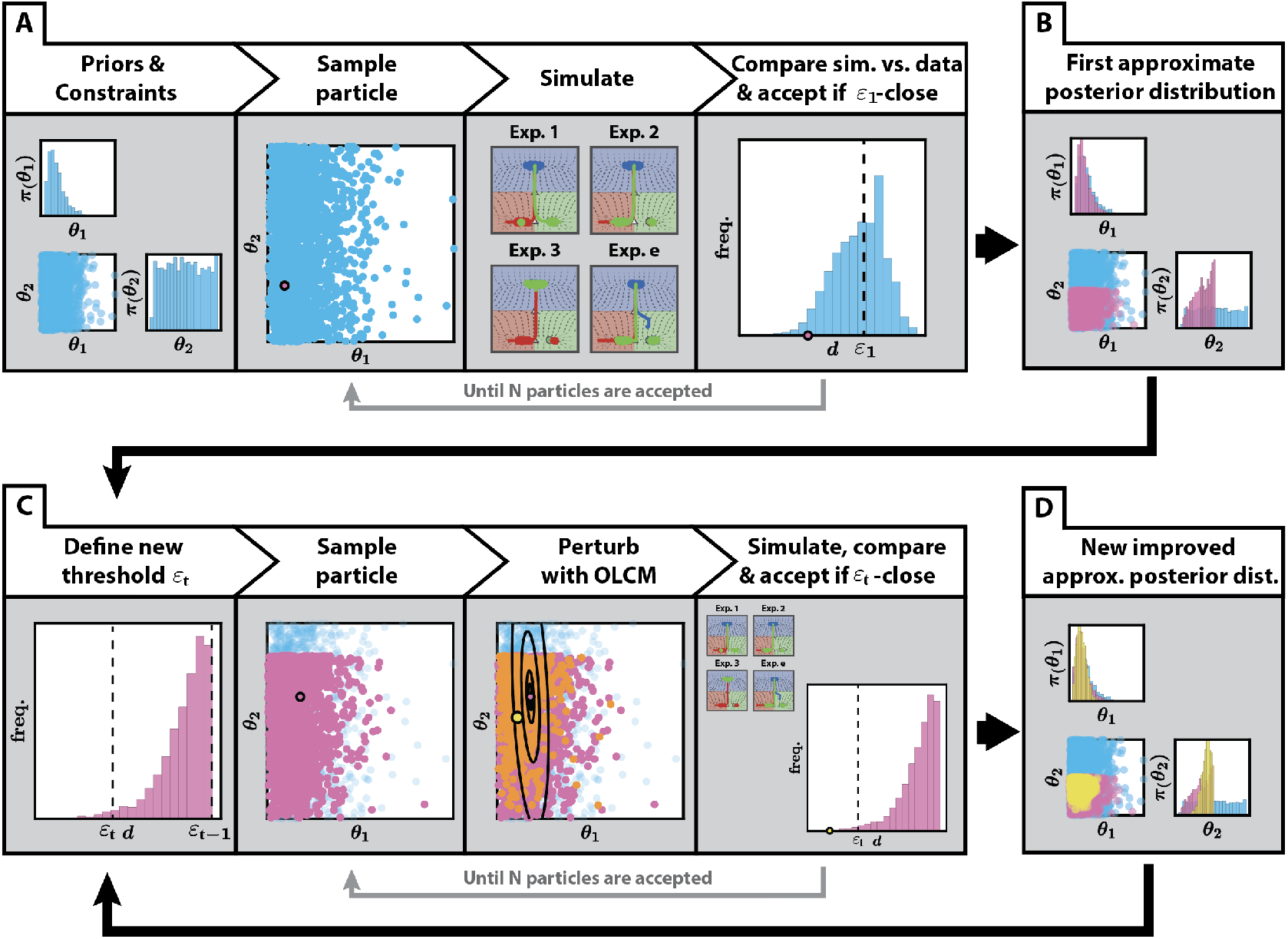
Application of ABC SMC to the fitting of the binary flip with cusp model. (A) First step of the algorithm. Having defined priors for the fitted parameters and constraints between them, as well as an initial threshold *ε*_1_, particles are sampled from the priors and accepted if the distance between the simulated data and the experimental data is less than the initial threshold. This is repeated until *N* particles are accepted. (B) An initial approximation of the posterior distribution is obtained, generated by the *N* particles accepted in (A). (C) The algorithm then proceeds in a sequential manner. In each step *t* of the algorithm, a new threshold *ε_t_* is defined, in our case, the 0.3 quantile of the previous distribution of distances *ε*_*t*−1_. Particles are sampled from the last approximate posterior distribution and perturbed using a Markov Kernel obtained from the Optimal Local Covariance Matrix (OLCM) (***Filippi et al., 2013***) (black ellipses in the figure represent confidence intervals), (as described in ***Appendix 2***). This is repeated until *N* new particles that satisfy the distance threshold are accepted. Also, each new particle is assigned a weight proportional to its prior probability and inversely proportional to the Markov Kernels evaluated at this particle, which control for effcient exploration. (D) After each step, a new improved approximate posterior distribution is obtained, which restricts the values of the parameters to a restricted region of the parameter set.

Our model depends on a total of 22 parameters: 1 time constant that controls the velocity of the trajectories in the landscape, 10 landscape parameters that map the signaling values into the shape of the landscape and 11 signaling parameters that define the signaling profile that each cell is exposed to in time (see ***Appendix 1***). Both the model’s definition and the experimental data constrain the range of the landscape parameters and inform the choice of their priors, for which we choose flat priors on their support. For the rest of parameters we choose fairly non-informative priors that still reflect our knowledge about their ranges. Moreover, taking into account the dimensionality of the parameter space, we choose to approximate the posteriors by sampling *N* = 2 × 10^4^ particles at each step of the algorithm. More details can be found in ***Appendix 2***.

In order to compare the experimental and simulated data we define a distance *d* that measures the level of similarity between two data sets. If 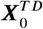 is the subset of data corresponding to experiments in the training data set (see ***Table 1***), we define the distance between 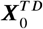 and the corresponding simulation ***X***^*TD*^ (***θ***) as

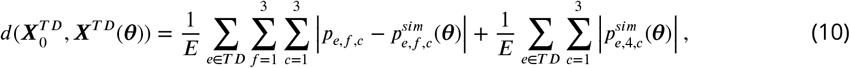

where *E* is the number of experiments in the training data set, *p_e,f,c_* and 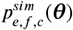 are the experimental and simulated probabilities of cell *c* becoming fate *f* in experimental condition *e*, respectively, and 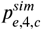 is the proportion of times that our model could not assign a fate to cell *c* when simulating experiment *e* (see ***Appendix 2*** for more information). The distance function penalizes parameter values for which our model cannot assign a fate, since we would like to avoid these parameter values. More details about our implementation of the fitting algorithm such as the choice of thresholds sequence and perturbation kernel function can be found in ***Appendix 2***.

Our choice of experiments for training was based on trying to find a maximally informative set while constraining the number of experiments. For example, we included all experimental conditions (a total of 6) with fully penetrant phenotypes resulting from different signalling combinations, which explored different regions of the signal space. With regards to partially penetrant phenotypes, we included one experimental condition involving EGF overexpression. We expected that including these experiments would inform the fitting of the values of the scaling parameters *s* and *l* and other parameter values related to the strength of the signals and the crosstalk between them. To constrain the temporal features of the model, we also added two AC ablation conditions which we considered the most informative ones, expecting that the model would be able to reproduce the rest. The particular details of the fitting can be found in ***Appendix 2***.

## Results

### Fitting the model to the training data: a two-step decision

The results after 14 generations of ABC SMC are shown ***Figure 4***, ***Figure 4***–***Figure Supplement 1*** and ***Figure 5*** (and supplemental videos), which show excellent agreement overall between the simulations and the data. We are able to reproduce both fully and partially penetrant phenotypes.

**Figure 4.**
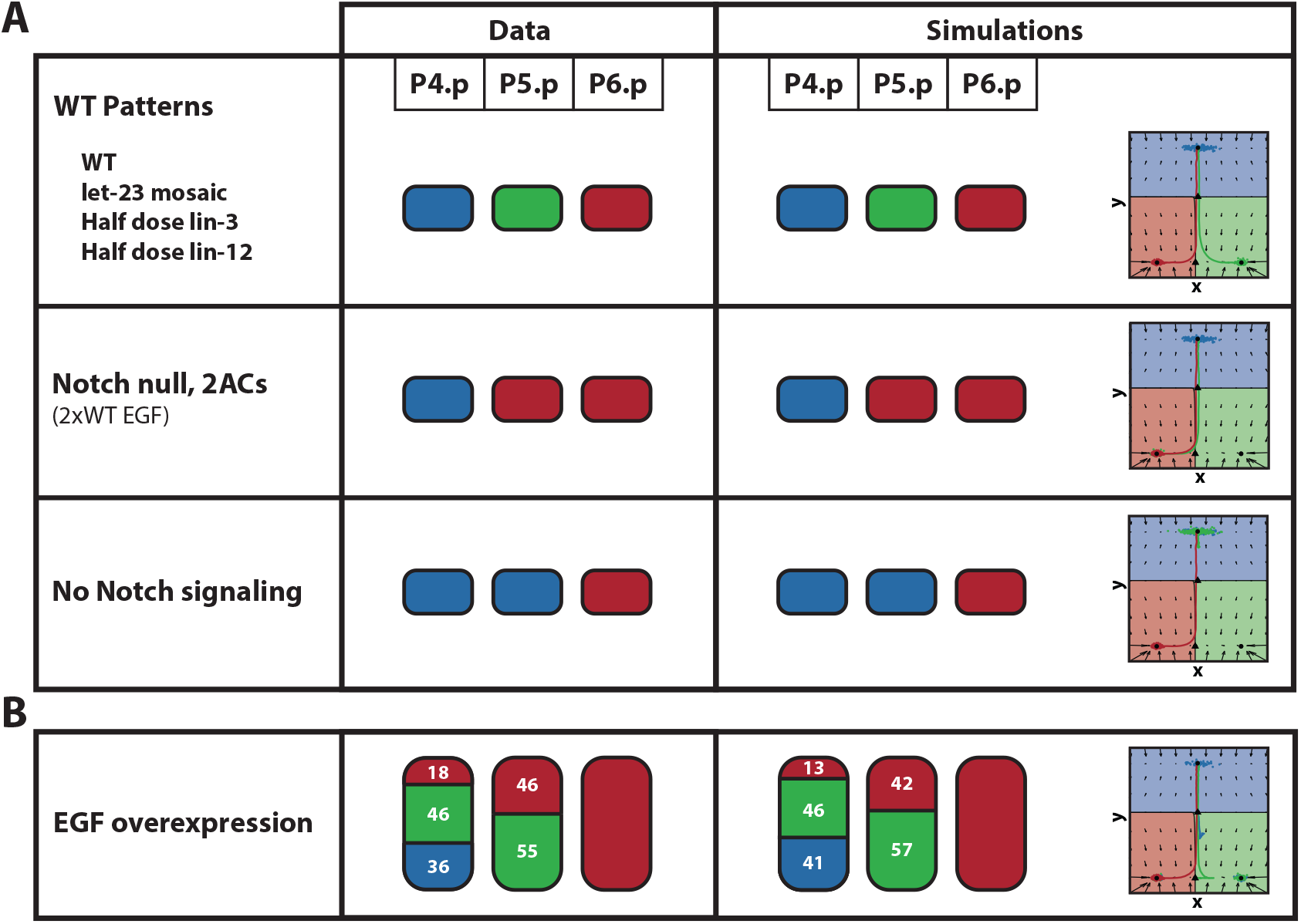
Fitting of the training data. (A) The patterns of the 6 fully penetrant phenotype perturbations considered in the training data set are correctly fitted by the binary flip with cusp model. (B) The patterns of the partially penetrant phenotype given by the EGF overexpression perturbation considered in the training data set is also correctly fitted by the model. On the left, the experimental patterns. On the right, the mean simulated patterns and trajectories on the landscape. In both the experimental and simulated patterns, primary, secondary and tertiary fates are represented by the red, green and blue colors, respectively, and proportions higher than 90% have been rounded. On the landscape, the mean simulated trajectories for the particle with best overall approximation for P4.p, P5.p and P6.p are colored in blue, green and red, respectively. **Figure 4–video 1.** Model dynamics of a simulated WT pattern. Trajectories of P4-6.p in the phase space (top) and their signalling profiles in the signal space (bottom). Each attractor and corresponding basin of attraction have been colored accordingly. **Figure 4–video 2.** Model dynamics of a simulated *let* − 23 mosaic mutant. Trajectories of P4-6.p in the phase space and their signalling profiles in the signal space. **Figure 4–video 3.** Model dynamics of a simulated half dose *lin* −3 mutant. Trajectories of P4-6.p in the phase space and their signalling profiles in the signal space. **Figure 4–video 4.** Model dynamics of a simulated half dose *lin* − 12 mutant. Trajectories of P4-6.p in the phase space and their signalling profiles in the signal space. **Figure 4–video 5.** Model dynamics of a simulated Notch null, 2ACs mutant. Top: Trajectories of P4-6.p in the phase space.Trajectories of P4− 6.p in the phase space and their signalling profiles in the signal space. **Figure 4–video 6.** Model dynamics of a simulated mutant with no Notch signaling and WT EGF. Trajectories of P4-6.p in the phase space and their signalling profiles in the signal space. **Figure 4–video 7.** Model dynamics of a simulated EGF overexpression mutant. Trajectories of P4-6.p in the phase space and their signalling profiles in the signal space.

**Figure 5.**
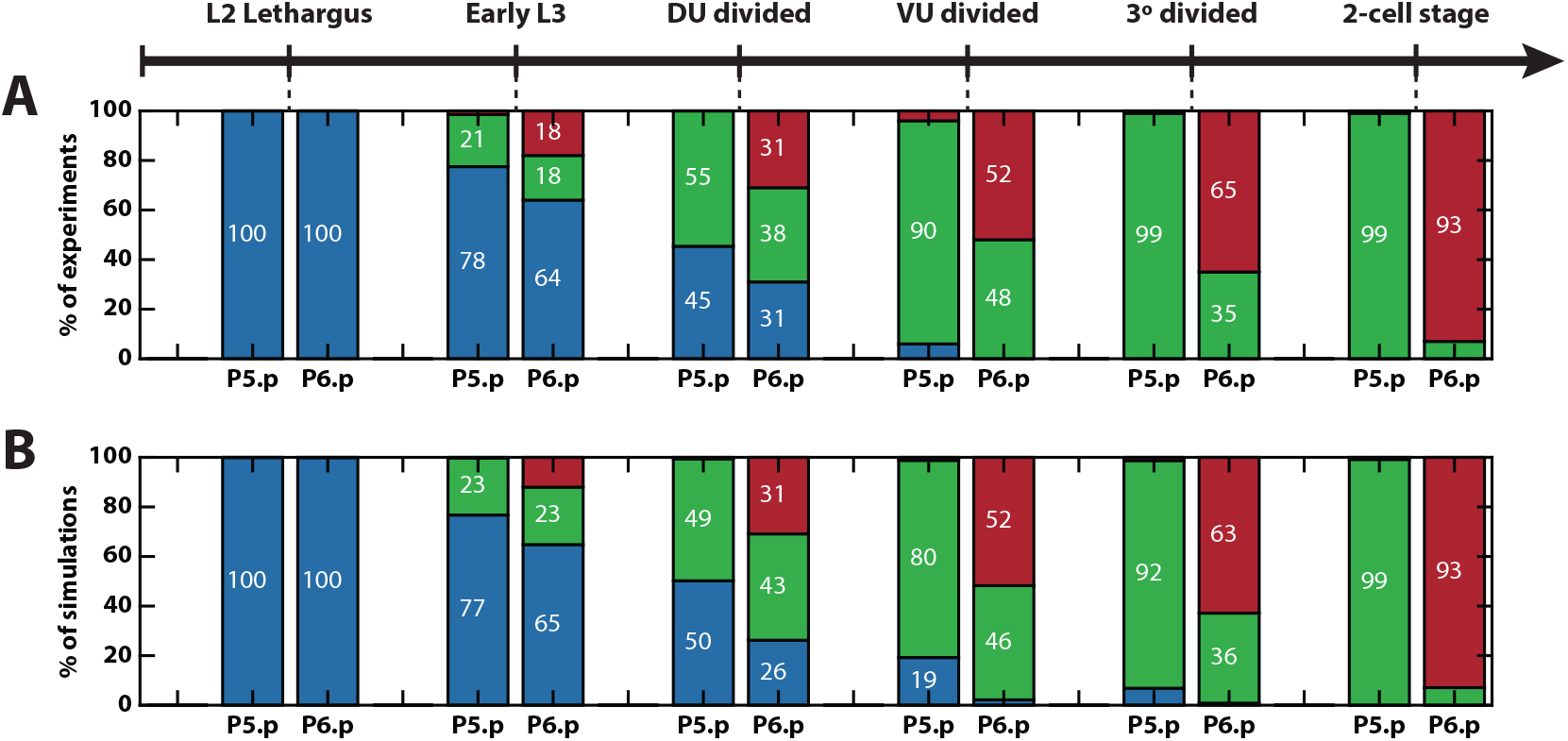
AC ablation fitting and validation. (A) Patterns observed when the AC is ablated at different developmental stages. (B) Mean simulated patterns for the different AC ablation conditions. Only proportions for P5.p and P6.p are available, which are presented in blue, green or red representing tertiary, secondary or primary fate, respectively. **Figure 5–video 1.** Model dynamics of a simulated mutant where the AC was ablated at the L2 lethargus stage. Trajectories of P4-6.p in the phase space (top) and their signalling profiles in the signal space (bottom). Each attractor and corresponding basin of attraction have been colored accordingly. **Figure 5–video 2.** Model dynamics of a simulated mutant where the AC was ablated at the early L3 stage. Trajectories of P4-6.p in the phase space and their signalling profiles in the signal space. **Figure 5–video 3.** Model dynamics of a simulated mutant where the AC was ablated at theDU divided stage. Trajectories of P4-6.p in the phase space and their signalling profiles in the signal space. **Figure 5–video 4.** Model dynamics of a simulated mutant where the AC was ablated at the VU divided stage. Trajectories of P4-6.p in the phase space and their signalling profiles in the signal space. **Figure 5–video 5.** Model dynamics of a simulated mutant where the AC was ablated at the 3° divided stage.Trajectories of P4− 6.p in the phase space and their signalling profiles in the signal space. **Figure 5–video 6.** Model dynamics of a simulated mutant where the AC was ablated at the 2-cell stage. Trajectories of P4-6.p in the phase space and their signalling profiles in the signal space.

***Figure 4***–***video 1*** shows how the simulated VPCs pattern in the WT case. The two-step decision logic of non-vulval vs vulval followed by primary vs secondary fates can be clearly seen. As P6.p starts receiving EGF signal, the cell leaves the attractor corresponding to 3° fate, i.e. the first decision is made, and moves into the regions of the fate map and landscape containing vulval fates. The increase in EGF signal and lack of Notch signal from its neighbors, positions its signaling profile in a region of the fate map where only primary fate is stable, and therefore, its trajectory on the landscape moves towards this attractor. A similar effect happens for P5.p but, due to the higher distance to the AC resulting in lower EGF signal, but higher Notch signal from the neighboring P6.p differentiating to primary fate, the cell differentiates towards 2° fate. Finally, the signals received by P4.p are not enough for it to escape the attractor corresponding to 3° fate, and therefore the cell differentiates into the non-vulval fate. ***Figure 4***–***video 2***-***video 7*** show the differentiation of VPCs in mutants 2-7 in ***Table 1***.

It is worth mentioning that, in our model, due to the lack of information about the exact timings of the stages at which AC ablations were performed, relative to each other, and also about the EGF and Notch signal dynamics, the increase in EGF and Notch signals are modelled by monotonically increasing functions, which are initially hypothesised to be sigmoidal functions of the simulation time. We find that the fitting of the data suggests a slight modification of the EGF dependence on time (see Appendix 1 Figure 17) which then allows for the model to fit the AC ablation data (***Figure 5***). This is probably due to the fast dynamics around the bifurcation point between the attractor corresponding to tertiary fate and the saddle. More details can be found in ***Appendix 1***.

Finally, we also observe that the results of fitting suggest that the geometry of the fate map is strongly constrained by the data, the details of which are shown in ***Appendix 2*** (see Appendix 2 Figure 3). Taken together, our results suggest that aspects from both the morphogen and the sequential model are important for the correct patterning, which combine into a two-step decision logic determined by the fitted fate map and controlled by the dynamics of the EGF and Notch signals received by the cells.

### Validating the model: the model reproduces epistasis between EGF and Notch

There is an extensive amount of experimental data available with different combinations of signaling perturbations. In particular, ***Barkoulas et al.*** (***2013***) provides quantitative signaling data for many perturbation lines. We compiled a set of mutants from the literature and tested whether our model was able to reproduce this data (***Table 1***). An important feature of the model developed in ***Corson and Siggia*** (***2012***, 2017) is that it can reproduce epistatic effects between the signals. Here we show that our model can also explain these effects.

As shown in ***Barkoulas et al.*** (***2013***), an EGF overexpression perturbation of 2.75-fold with respect to the WT (based on measured *lin-3* mRNA levels), named JU1107, increased P5.p induction towards primary fate, and P4.p towards secondary fate. Our model correctly reproduces these features as shown in ***Figure 6***A and ***Table 2***. This is achieved because an increase in the EGF signal of P5.p by that magnitude locates its signalling profile close to the cusp mid-line in the signal space, where primary and secondary fates are equally probable, and P4.p closer to the fold line in a region where secondary and tertiary fates are equally probable (***Figure 6***A).

**Figure 6.**
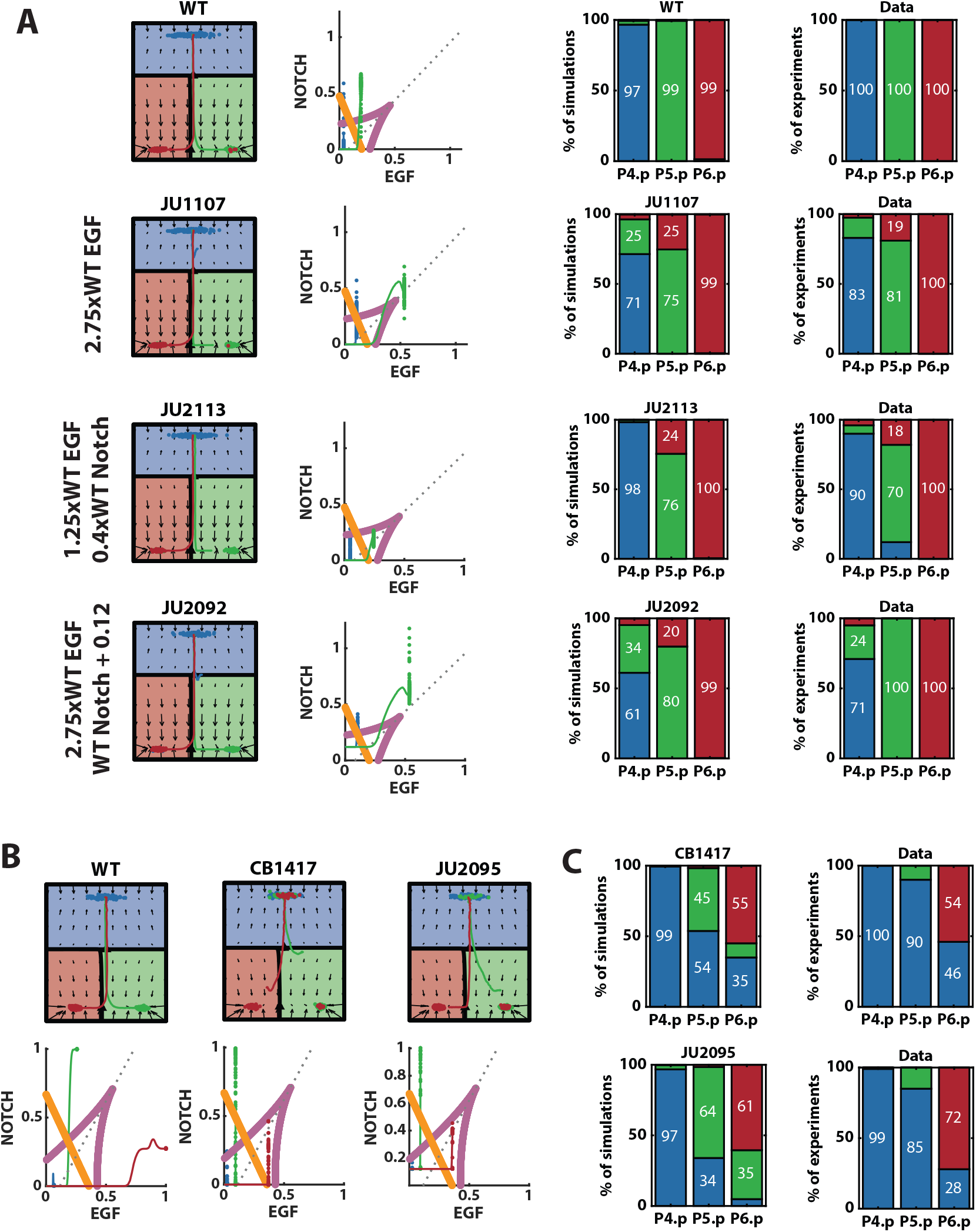
Model’s further validations. (A) Left: Representative mean trajectories on the landscape and signal space for simulated WT vulva and different signal perturbations included in the validation data set. Blue, green and red trajectories represent the evolution of the state of P4.p, P5.p and P6.p, respectively. Right: Mean simulated and experimental patterns for the different perturbations shown on the left. Blue, green and red represent tertiary, secondary and primary fates, respectively. (B) Representative mean trajectories on the landscape and signal space for simulated WT vulva and EGF hypomorph perturbations. Blue, green and red trajectories represent the evolution of the state of P4.p, P5.p and P6.p, respectively.(C) Mean simulated and experimental patterns for the EGF hypomorph perturbations. Blue, green and red represent tertiary, secondary and primary fates, respectively.

**Table 2.**
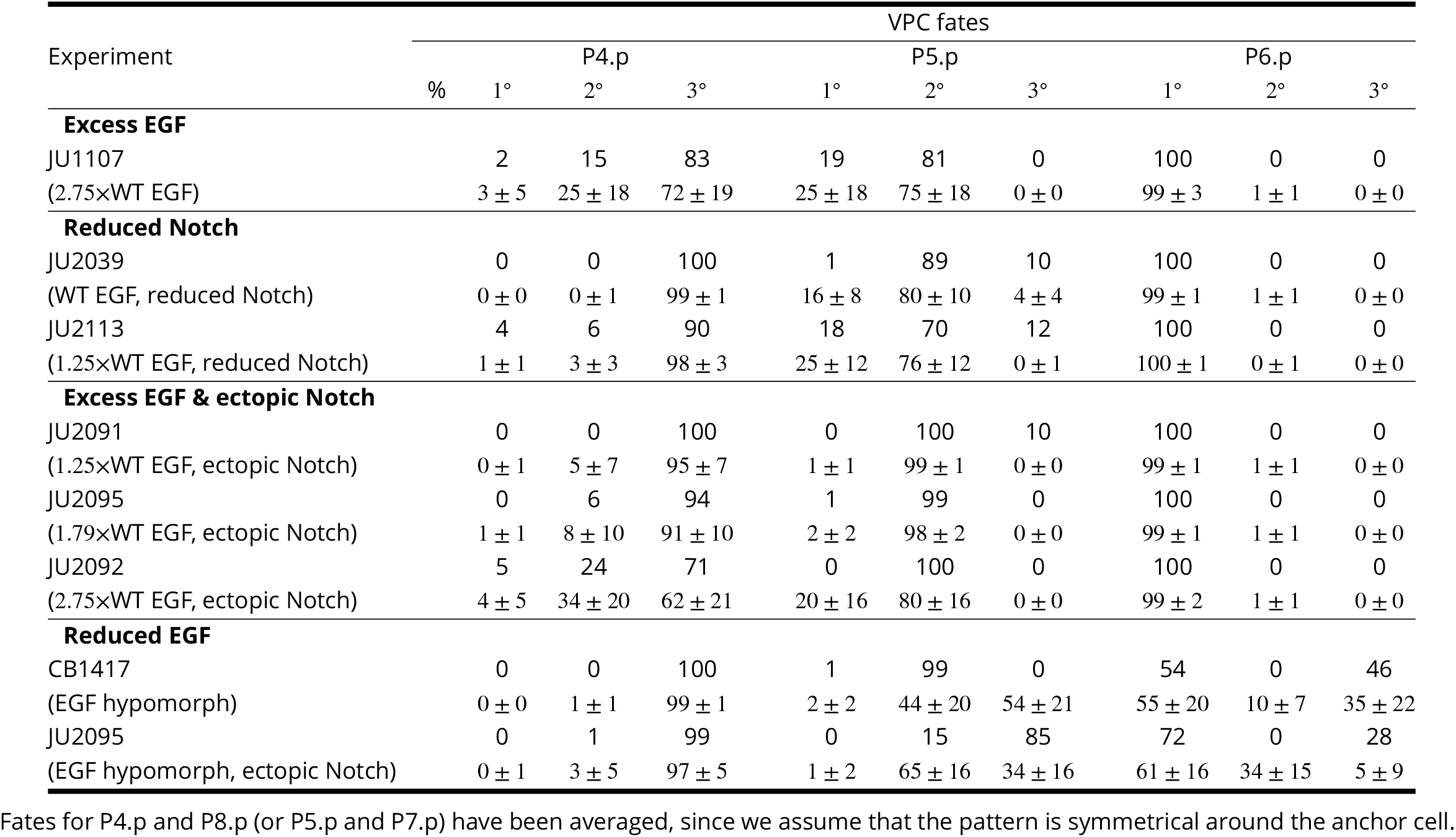
Comparison between experimental and simulated data for the validated experimental conditions.

Reduced Notch perturbations do not have a strong phenotype unless crossed with an EGF overexpression mutant (see ***Table 2***), in which case the probability of P5.p differentiating into secondary fate is slightly reduced. We checked whether our model was capable of reproducing these features. Since the level of reduction of Notch is not quantified in the experimental settings, we fitted the multiplicative magnitude of Notch reduction that reproduced the data. With a Notch reduction of 0.4×WT, our model is able to reproduce both JU2039, where EGF is WT and there is no strong phenotype, and JU2113, where EGF is slightly increased by 1.25-fold and secondary fate in P5.p is destabilized, increasing the probability of primary and tertiary fates. This fitted fold increase is, in fact, similar to the one considered in ***Corson and Siggia*** (***2017***). The model predicts that, under JU2039 signaling regime, P5.p would stay in a region of the signal space where secondary fate is still predominant. Although our model is not capable of reproducing a small number of tertiary fated P5.p cells under the JU2113 perturbation, it does reproduce the increase in primary fated P5.p, by approaching the cusp mid-line in the signal space, but not leaving the bistability region inside the cusp (***Table 2*** and ***Figure 6***A).

We also tested whether the model is able to reproduce mutants with ectopic Notch expression. Similar to the previous subset of data, these perturbations are silent unless crossed with EGF overexpression perturbations. As in ***Corson and Siggia*** (***2017***), we modeled these perturbations as an additive constant for the Notch signaling parameter (see ***Appendix 1***). We fitted this constant to the experimental outcome of the JU2092 mutant, with a resulting constant equal to 0.12. With this, we were able to fit the three crosses shown in ***Table 2***. Interestingly, in the strongest EGF overexpression perturbation, P4.p was positioned close to the right of the intersection between the cusp mid-line and the fold line, giving a mix of tertiary and secondary fates, as observed in the data (***Figure 6***A).

Finally, we simulated EGF hypomorph mutants. Interestingly, these perturbations show that a strong decrease of EGF signals gives a mix of primary and tertiary fated P6.p cells while not affecting the WT pattern of the rest of VPCs. Moreover, a cross with a mutant with mild ectopic Notch activity, such as the one considered above, promotes primary fated P6.p, showing again epistasis between the signals. To simulate these perturbations, we fitted a multiplicative downregulation of EGF signal to the egf hypomorph data, CB1417. With a reduction of 0.36×WT EGF, our model can reproduce the mix of tertiary and primary fated P6.p observed in the EGF hypomorph mutant (CB1417) (***Figure 6***C). This is achieved because a reduction in EGF positions the signalling profile of P6.p close to the fold line while still being to the left of the cusp-midline (***Figure 6***B). Simulating the cross of this perturbation with ectopic Notch (mutant JU2095) moves P6.p signaling profile further from the fold line, promoting vulval fates. However, it also positions it closer to the cusp mid-line and therefore, also promotes secondary fates in P6.p in our simulations (***Figure 6***B). The model is also not able to reproduce the high probability of P5.p staying in tertiary fate. We believe that the differences between simulations and experiments observed in P6.p could be fixed by adding the perturbations to the training data set. Reproducing the differences observed in P5.p would likely require a non-linear mapping of the fold into the signal space.

### Model predictions

Having reproduced a large set of data, we explored whether the model could predict interesting outcomes.

We tested whether crossing two silent perturbations in the training data could show a new epistatic event. Halving EGF ligand or Notch receptors does not affect the wild type phenotype (***Table 1*** (3-4)). In our model, this is because making either of these two perturbations leaves cells within the same stability regions of the fate map. However, our model suggests that, if both perturbations are crossed, we observe epistasis, where induction of P5.p to secondary fates is strongly reduced by almost half (***Figure 7***B). This is consistent with predictions shown in ***Corson and Siggia*** (***2012***). Biologically, this prediction suggests that EGF promotes secondary fates which, at first, could seem counterintuitive.

**Figure 7.**
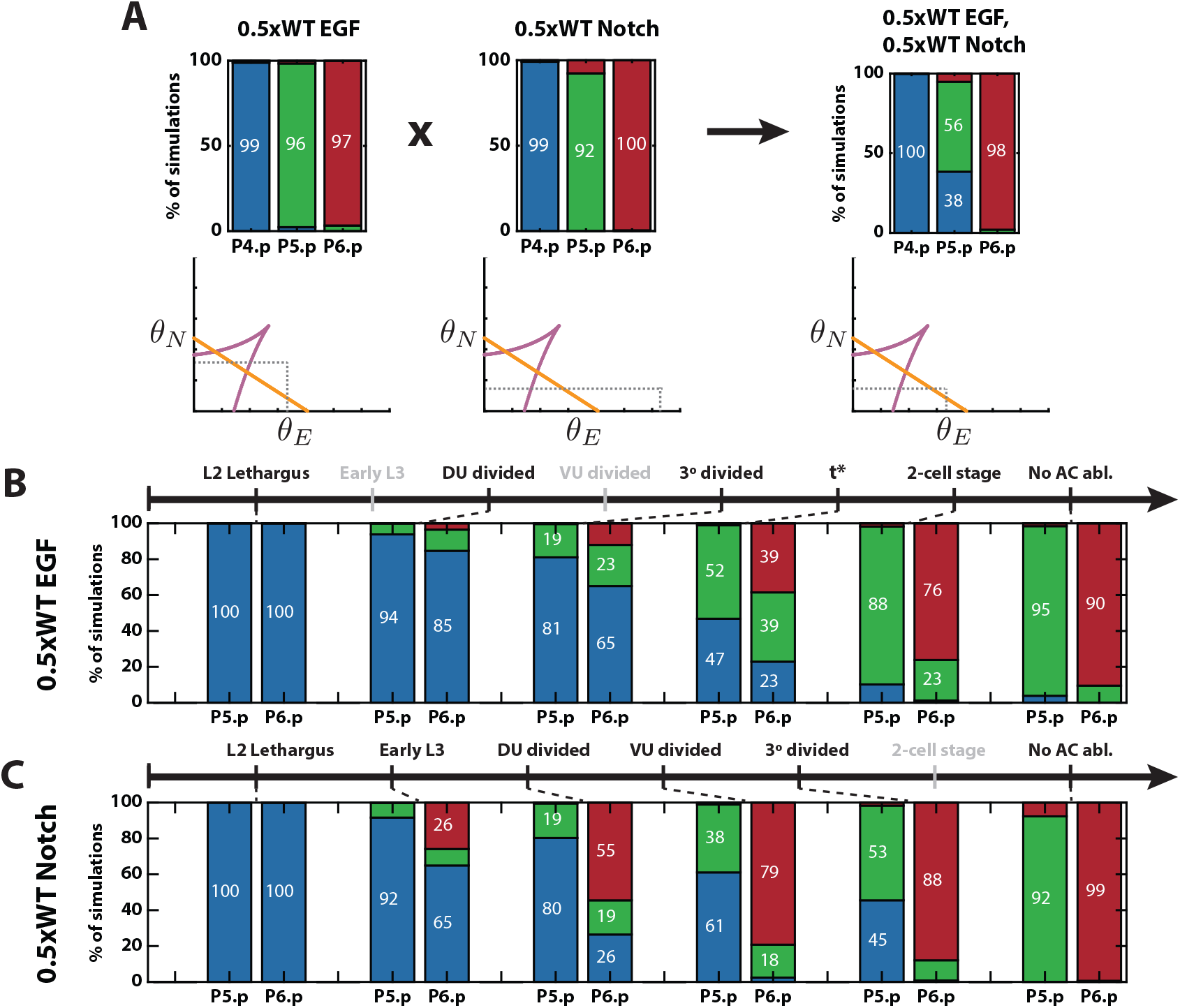
Model’s predictions. (A) Predicted epistasis under the cross of reduced EGF ligand and Notch receptors perturbations. Top: Simulated proportions under the corresponding signaling profiles. Bottom: Schematics of the simulated positions of P4.p in the signal space under the different signaling profiles. (B) Predicted pattern proportions for different AC ablation times under a reduced EGF ligand perturbation. *t*\* is a predicted time between 3° divided and 2-cell stages. The stages for which data is not shown are colored in gray. (C) Predicted pattern proportions for different AC ablation times under a reduced Notch receptors perturbation. The names of the stages for which data is not shown are colored in gray.

AC ablation experiments are very insightful, as they are the only experiments that provide information about the trajectory of the cells in the landscape. However, AC ablation data is only available under a WT signaling profile. As we mentioned earlier, a wild type pattern is achieved even under either half dose *lin-3* or *lin-12*. However, it is not known whether the pattern is dynamically formed in the same way as under WT signaling. We used our model to test this and observed an interesting effect.

Simulating AC ablation under a half dose of *lin-3*, the EGF ligand, showed that the process followed the same order as under WT signal, but proceeded more slowly (***Figure 7***C). Under this condition, our model predicts (1/3, 1/3, 1/3) probabilities for P6.p becoming one of the three fates if AC is ablated at a time between 3° divided stage and 2-cell stage. However, under a half dose of *lin-12*, a Notch receptor, our model predicted that there is very low probability of P6.p becoming a secondary fate under any ablation time, as observed in the WT and *lin-3* mutant. Instead, there is a direct transition from tertiary to primary fate. This is also consistent with the effect observed in ***Corson and Siggia*** (***2012***).

## Discussion

Here we have presented a method to build simple landscapes starting from qualitative observations, which can then be fitted to a large amount of quantitative data. With this method we have shown that a very simple three-way landscape is able to reproduce the complex vulval patterning process in *C. elegans*. Given the number of stable cell fates observed in the data, CT is a powerful theory to classify and build landscapes with the desired characteristics. By coupling it with an effcient parameter fitting method, such as the one presented based on ABC SMC, one can check which proposed models are consistent with the data and choose the simplest one.

An important difference between our model and the one developed in ***Corson and Siggia*** (***2012***, 2017) is that while in their model all states are equivalent, giving a three-fold symmetric potential, in our model, the cells go through two cell state transitions: a first decision between vulval and non-vulval fates, and then, for cells that adopt vulval fates, a decision between primary and secondary.

This may be a general landscape which can be used whenever a cell first decides whether to leave a precursor state or not, and then, once leaving that state, decides between a pair of options.

Here we have taken advantage of vulval development in *C. elegans* to illustrate this new mathematical framework, however, this is a very complex problem involving patterning of several cells that interact through both external and paracrine signals, and therefore, the state of each cell depends upon that of its neighbors. Moreover, we lacked detailed information about cell and signal dynamics in time, as data is only available on the final phenotypes. The only data available on dynamic perturbations were those in which the AC was ablated and even in this case, again the only available phenotype was the final one and the specific ablation times were not available. This resulted in relatively large number of unknowns and corresponding simplifying assumptions, such as the model for EGF increase in time, which complicated the model and the analysis. This approach, however, is ideal for modelling single cell data for which one is able to observe cell transitions along the protocol time as well as controlling signaling in time, as we show in ([article to be submitted]).

An advantage of using this approach, where the model is built from basic transitions, is that the system can be easily evolved. Here we have postulated that the mapping between signals and control parameters was linear, and we were able to reproduce a great amount of data. However, we also showed that the model struggled to reproduce some details observed in the EGF hypomorph perturbation. This could suggest that a more sophisticated mapping would be needed in this case, and with our approach, it would be possible to locally modify the mapping in the low EGF region without changing the remainder of the model.

With the fast development of experimental techniques to observe single cells, such as single cell RNA-seq together with live signaling and cell fate reporters, vast amounts of data are becoming available. Building gene regulatory networks that account for all the subtleties observed in the data proves to be challenging. Moreover, we can expect that the dynamics of any GRN will be largely described by the normal forms that Catastrophe Theory and Dynamical Systems Theory provide. Therefore, the framework proposed here, that centers on the essence of the process without the complicating molecular details, can alleviate these challenges and, in fact, could benefit from this new information. This approach to building landscape models has several advantages in comparison to GRN models. First, as mentioned before, since it focuses on the essence of the process rather than on the mechanistic details, one can build a landscape model from simple qualitative data about the state transitions. Secondly, while there are many network structures that can reproduce the similar observations, there are only a few distinct landscape topologies with a given number of attractors. We believe there are many exciting applications of this framework to more cell differentiation processes.

## Acknowledgments

We thank Eric D. Siggia and Francis Corson for extensive conversations and advice. We also thank Miguel Ángel Ortiz Salazar for his feedback on the manuscript. This research was partly completed while E.C. was a PhD student, supported by the University of Warwick. Some of this work was also performed at the KITP Santa Barbara and therefore this research was supported in part by the National Science Foundation under Grant No. NSF PHY-1748958, NIH Grant No. R25GM067110, and the Gordon and Betty Moore Foundation Grant No. 2919.01.

## Appendix 1

## Concepts from dynamical systems and catastrophe theory

Here we will provide some basic concepts in the fields of catastrophe and bifurcation theory for readers unfamiliar with them. Most of the concepts summarised in this section are taken from the book (***Poston and Stewart, 2014***), where we refer the reader if interested in more details.

Let *V* be a family of functions which, in our settings, will represent the familly of potential functions defining a landscape model:

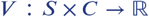

where *S* is a manifold in **ℝ**^*n*^ and *C* is another manifold in **ℝ**^*r*^. Let us call **ℝ**^*n*^ the **state space** and **ℝ**^*r*^ the **control or parameter space**. For a fixed value ***c*** in *C*, we will denote by *V_**c**_* : *S* → **ℝ** the potential function *V_**c**_* (***x***) = *V* (***x, c***) for ***x*** ∈ *S*.

For a fixed value ***c*** for the **control variables** or **parameters**, the function *V_**c**_* (***x***) will have certain critical points. Those will be given by the points ***x***\* such that *D_X_V*_*c*_ (***x***\*)= 0, i.e. the gradient of the potential at that particular point is zero. The critical point is **degenerate**if the rank of the Hessian matrix of *V_**c**_* evaluated at ***x***\* is not maximum, i.e. 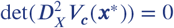. If the critical point is not degenerate, we say that it is **non-degenerate**. We are interested in knowing how the number of critical points and their stability change depending on the value ***c***, the parameters.

Let us define the **catastrophe manifold** 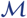 as the subset of **ℝ**^*n*^ × **ℝ**^*r*^ defined by:

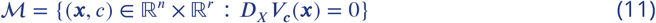

where *D_X_* is the gradient in the *X* variables. In other words, 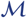 is the set of points (***x, c***) such that ***x*** is a critical point of the function *V_**c**_*, i.e. it contains all the critical points of the family of functions *V*.

The **catastrophe map** *χ* is the restriction to 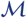 of the natural projection

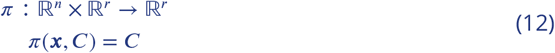

The **singularity set** 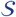 is the set of singular points in 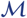 at which *χ* is singular, that is, where the rank of the derivative *D_χ_* is less than *r*. Actually, it is not hard to show that 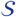 is the set of points 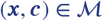 at which *V_c_* (***x***) has a degenerate critical point. The image 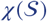 in *C* is called the **bifurcation set**. It follows that 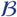 is the set on which the number and nature of the critical points change.

Summing up, the catastrophe manifold gives a description of the critical points for the family of functions *V_**c**_* (***x***), and thus it describes how they change as the parameters change. The bifurcation set describes the set of parameters for which important changes in the critical points take place.

In what follows, we will introduce these concepts in the context of two particular families of functions, the fold and the cusp, which we will use to build the binary flip with cusp landscape model.

## Fold catastrophe

The fold is the most basic catastrophe and it has the form:

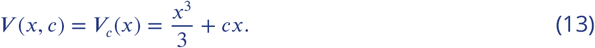

It is a cubic equation with graph always crossing the point *x* = 0. As we will see, depending on the value of the control parameter *c*, the potential function *V_c_* will have two, one or no critical points (Fig. Appendix 1 Figure 1A1–4).

**Appendix 1 Figure 1.**
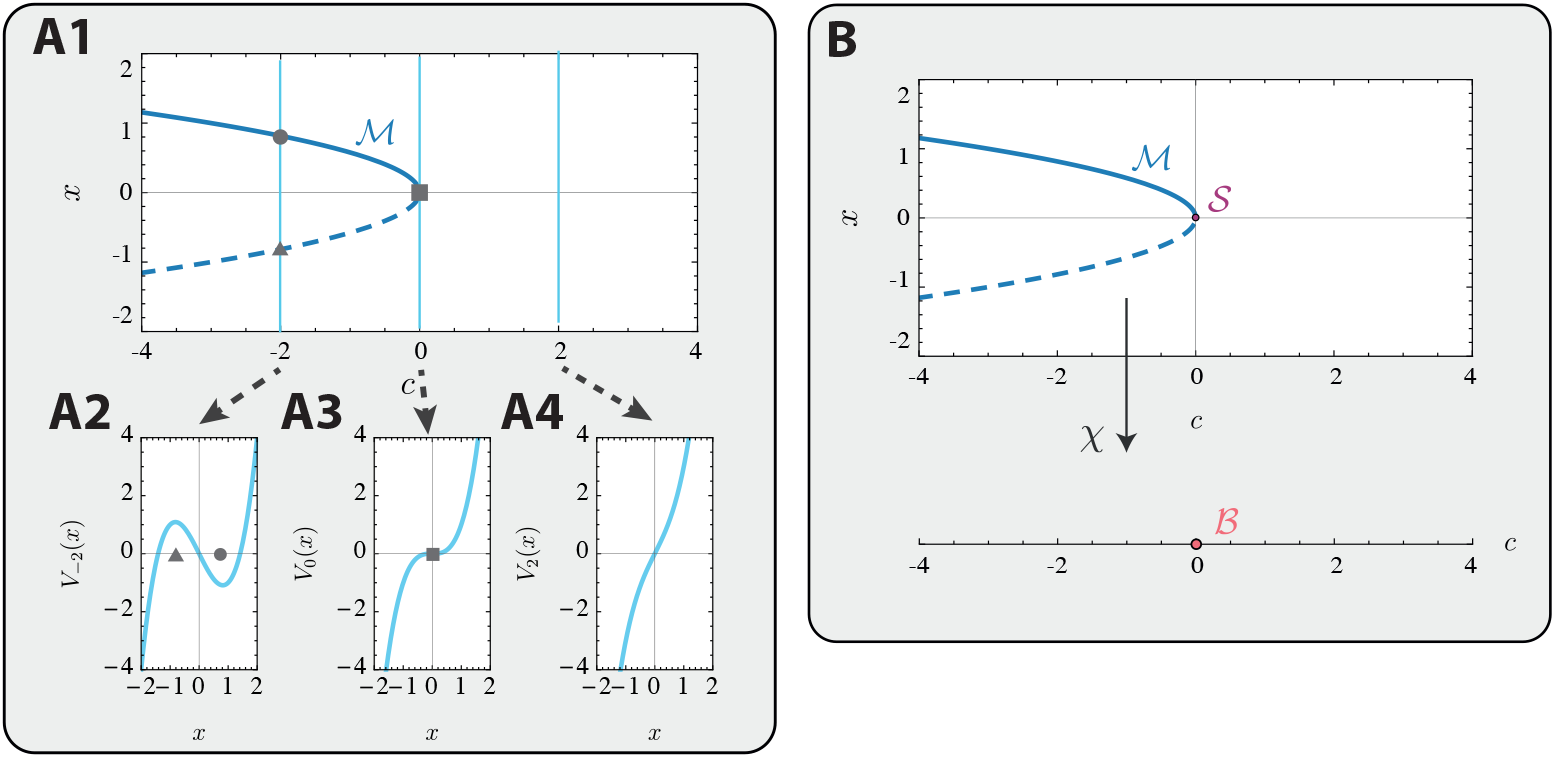
(A1) Bifurcation diagram or catastrophe manifold of the fold catastrophe. Catastrophe manifold 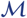. (A2-4) Plots of the fold potential for the corresponding values of the control parameter *c*. Continuous line in the bifurcation diagram and gray circle represent stable points, dashed line and gray triangle represent unstable points, gray square represents degenerate point (A2) In this case *c* = −2, for which the potential has two critical points, a minimum and a maximum. (A3) In this case *c* =0 and corresponds to a bifurcation point. The potential function only contains a degenerate critical point at *x* = 0. (A4) In this case *c* =2 and the potential function does not contain any critical point. (B) Catastrophe manifold 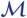, catastrophe map *χ*, singularity set 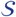 and bifurcation set 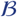 of the fold catastrophe given in Eq. 15. Again, continuous line represents stable points, dashed line represents unstable points.

The catastrophe manifold 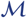 is given by the critical points of the potential function for each value of *c*:

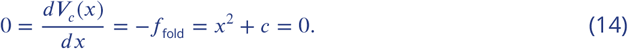

Therefore we can use the *x*-coordinate as a chart for 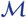 and write it as

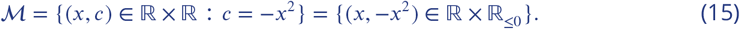

This means that for each value of c, the potential function has two critical points (*c* < 0), one critical point (*c* = 0) or no critical points (*c* > 0) (Fig. 1A1).

If we look at the stability of such critical points, 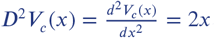, which evaluated at the points (*x*, − *x*^2^) of the catastrophe manifold, is equal to 2*x*, which sign depends on the sign of *x*. If *x* < 0, the critical point is unstable. If *x* > 0, the critical point is stable. And if *x* = 0 the critical point is degenerate (Fig. 1A1-4). In fact, the singularity set is the set of points 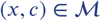 such that

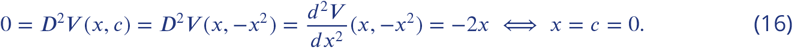

Hence, the singularity set 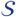 of the fold catastrophe is the point (0, 0) and the bifurcation set 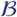 is the point *c* = 0 in the control space (Fig. 1B).

The catastrophe manifold helps us to identify regions in the control space for which the potential function will have the same critical points. For example, for the fold, if *c* < 0, the potential will contain two critical points (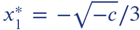 and 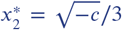); if *c* = 0, the potential has just one degenerate critical point (*x*\* = 0); and if *c* > 0 the potential function *V_c_* (*x*) contains no critical points (see Figure 1).

## Cusp catastrophe

This catastrophe has the form

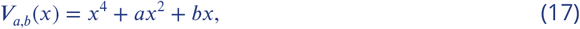

which is a polynomial of order 4 in one variable. In this case, the control space is 2-dimensional on *a, b*, and depending on their values, this potential function will contain one, two or three critical points (see Appendix 1 Figure 2A1–4).

**Appendix 1 Figure 2.**
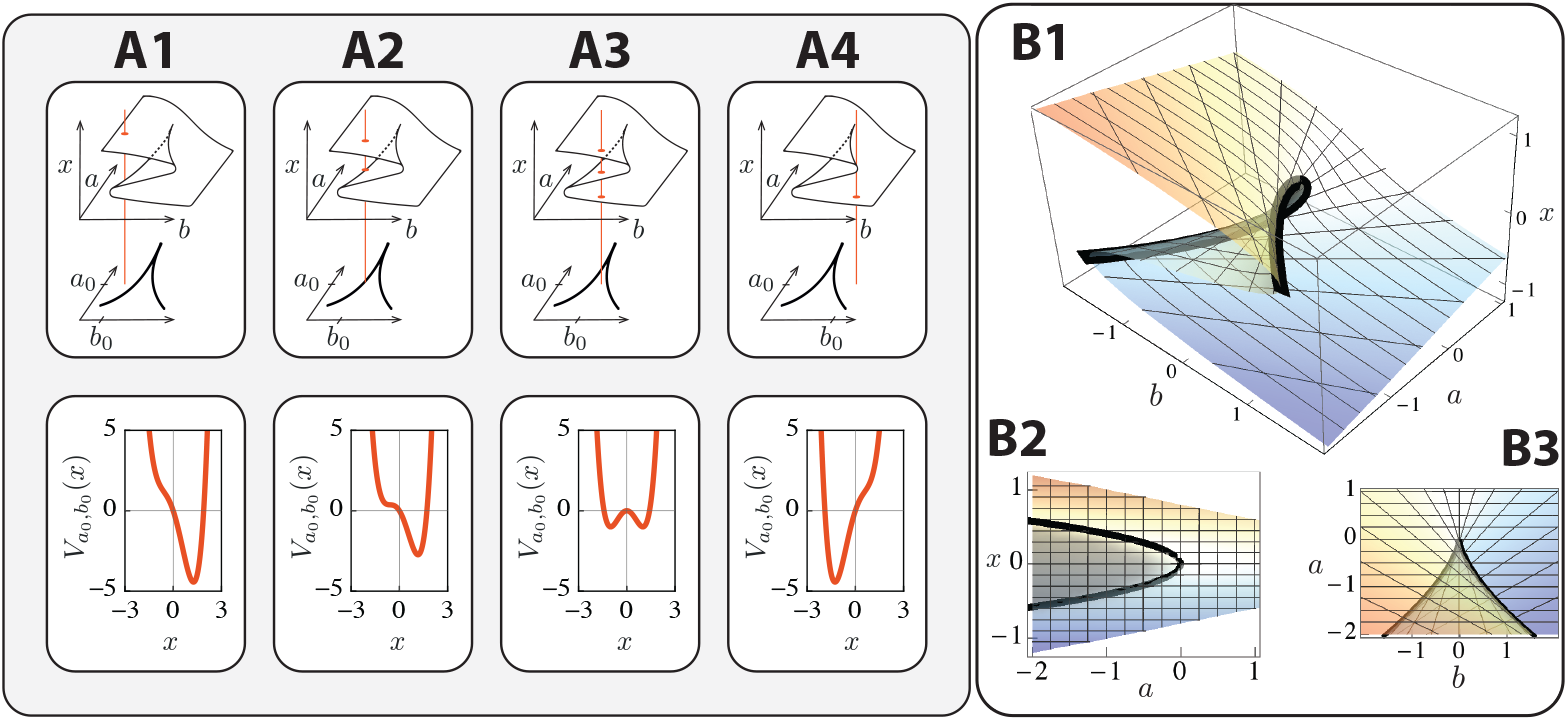
(A1-4) Four different potential functions from the cusp family for control parameters in different regions of the control space. Each row of plots shows, on the top figure, the catastrophe manifold intersected with the line (in orange) 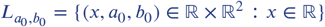 for a fixed value *a*_0_ < 0 and different values of *b*_0_, displaying the critical points (orange ellipses); and, on the bottom figure, the potential function corresponding to such values of *a* and *b* with the predicted critical points. (A1) The control parameters are such that 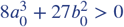 and *b*_0_ < 0, therefore the potential has only one critical point with *x* > 0. (A2) In this case 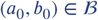 and *b*_0_ > 0, so the potential has two critical points, one of them is degenerate. (A3) The control parameters are such that 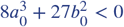 so the potential function has three critical points. (A4) Similar to (A1), The control parameters are such that 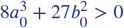 but now *b*_0_ > 0, therefore the potential has one, now negative, critical point. (B1) Catastrophe manifold 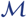 of the cusp catastrophe. Singularity set represented as a thick black line. (B2) Different view of the catastrophe manifold. Projection onto the (*x, a*) space. (B3) Projection of the singularity set onto the control space. The thick black line represents the bifurcation set of the cusp in the control space.

The catastrophe manifold 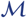 is given by the points in **ℝ** × **ℝ**^2^ such that

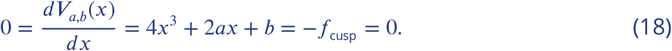

Hence, we can use (*x, a*) as a chart on 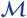 and write *b* in terms of *x* and *a* as *b* = −4*x*^3^ − 2*ax*. This allows us to write the catastrophe manifold as the set of points

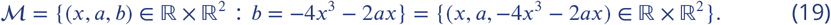

Appendix 1 Figure 2B1 contains a plot of this two dimensional manifold in **ℝ**^3^.

The singularity set 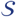 of the cusp catastrophe is given by the points in the catastrophe manifold such that

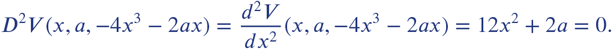

Therefore, *a* = −6*x*^2^; and since *b* = −4*x*^3^ − 2*ax* in 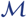, then

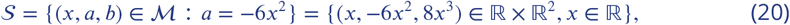

which is a curve in **ℝ**^3^ (see Appendix 1 Figure 2B1).

Finally, the bifurcation set 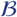 is given by the image of this set by the catastrophe map, which is the projection of this curve on the control space:

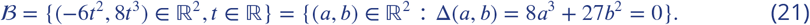

As we see in Appendix 1 Figure 2B3, the bifurcation set defines three regions in the control space. For values of the control parameters (*a*_0_*, b*_0_) in the region such that 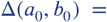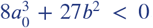, the potential function contains three critical points (two stable and one unstable). If (*a*_0_*, b*_0_) is in the bifurcation set, the potential function contains two critical points (one stable and one degenerate). And if the control parameters are in the region such that Δ(*a*_0_*, b*_0_) > 0, then the potential function contains just one stable critical point (see Appendix 1 Figure 2). Since the critical points are given by the zeros of a third-degree polynomial, the critical points in each case can be easily computed with Cardano’s formula.

## Stability of the binary flip with cusp landscape model

It follows from CT that these two bifurcations are universal for gradient-like systems in the following sense. Suppose a gradient-like system 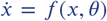 with state *x* = (*x*_1_, … , *x_n_*) and parameters *θ* = (*θ*_1_, … , *θ_d_*) has a bifurcation at *x* = *x*\*, *θ* = *θ*\* that is generic for 1-parameter (resp. 2-parameter) systems, then for (*x, 0*) near (*x*\*, *θ*\*), *f* is induced from the normal form *g* = *f*_fold_ (resp. *g* = *f*_cusp_) for the saddle-node (resp. cusp) in that there is a change of coordinates for which

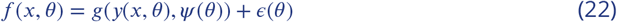

where *ψ* (*θ*) are the relevant parameters of the normal form [refTrotman]. By constructing our landscape from these two catastrophes is that it inherits this universality property and that as we will see now, the position and stability of the critical points and their dependence on the parameters is transparent.

The critical points of the binary flip with cusp model, defined in Eq. 2 in the main text, are given by the zeros of 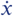 and 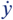.

First, let us focus on the zeros of the equation regarding the change in time of *y*, since it does not involve *x*. This equation involves the flow determined by the fold, where we have introduced an additional parameter ***M***, which allows for control of the position of the critical points:

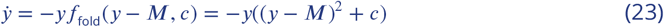

The possible zeros of 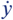 are 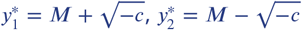 and 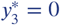. Since we assume that the state variables are real, 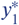 or 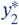 will only exist if *c* ≤ 0.

The stability of the points 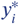 will depend on the sign of 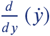 evaluated at the corresponding equilibrium point. Fixing the parameter ***M*** > 0, Appendix 1 Figure 3 shows the corresponding bifurcation diagram. Note that the value of the parameter ***M*** does not really change the stability of the system, qualitatively. It only affects the interval in which *c* can take values, and the coordinates of the equilibrium point. We have decided to take positive values of ***M***, and *H*(*y*) to be the Heaviside function described in the main text. If ***M*** was negative, one would just need to take a different step function which value is 1 for *y* ∈ [0, ∞), and this would just flip the state space over the *y*-axis.

**Appendix 1 Figure 3.**
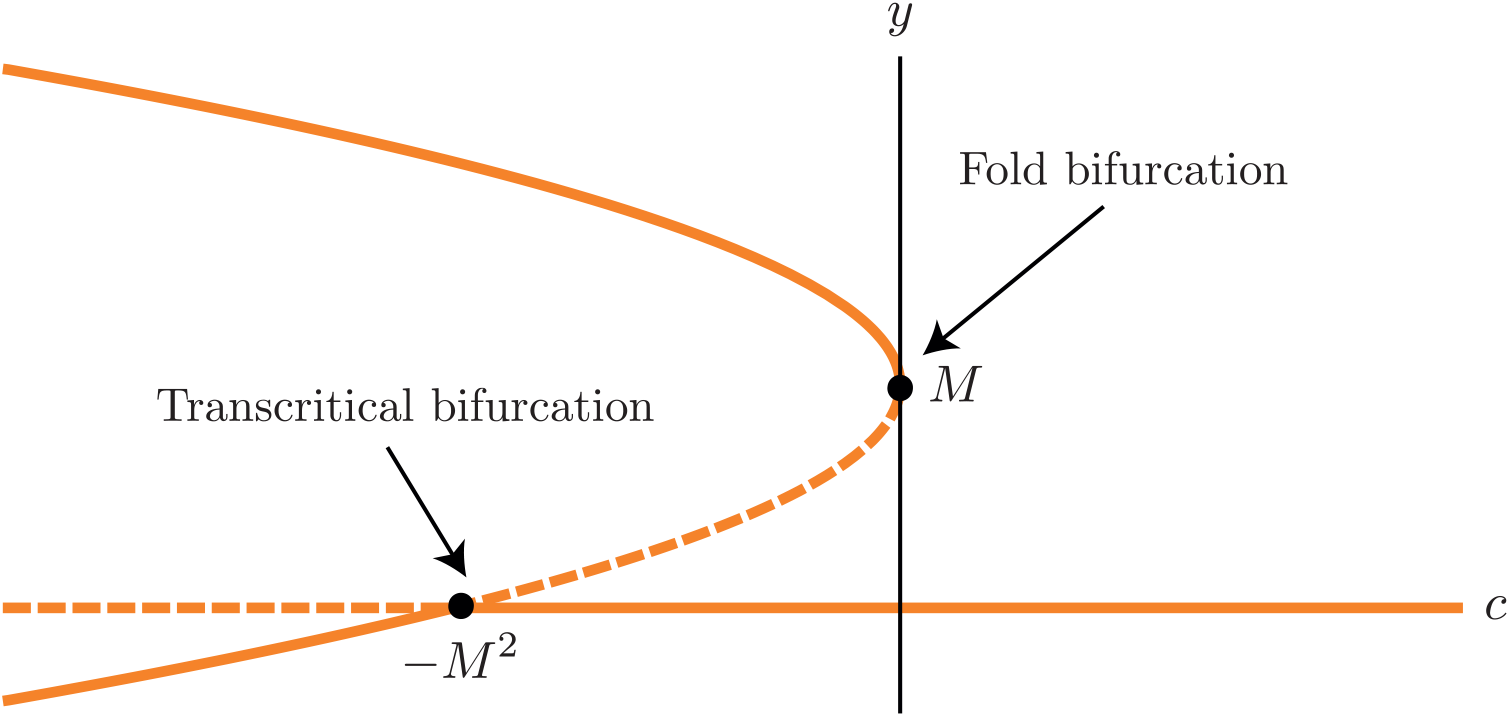
Bifurcation diagram of Equation (23). With ***M*** > 0 fixed, the orange lines show the equilibria of the system for different values of *c*. Continuous lines represent stable equilibria and dashed lines represent unstable equilibria.

We are only interested in the fold bifurcation that happens when *c* = 0, and not in the transcritical bifurcation at *c* = −***M***^2^. Therefore we will assume that

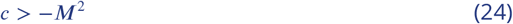

from this point onwards. The parameter *M* does not affect the stability or the bifurcation of the system, it only controls the position of the critical points on the *y* axes and constrains the value of the parameter *c*. This is why we will not consider it as a control parameter in what follows. Summing up, depending on the value of *c*, Eq. (23) has one (*c* > 0), two (*c* = 0) or three (*c* < 0) equilibria.

Now let us consider the complete system described in Eq 2. of the main text, and study the *x*-coordinates. In order to find the equilibrium points of the whole system we need to find the zeros of both equations. For 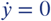, as we saw before, the equation will have one, two or three zeros depending on the value of *c*. Let us now study the zeros of 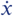 in the case when *c* < 0, as the two other cases follow a similar logic.

When *c* < 0, as we saw earlier, 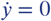 has three possible roots: 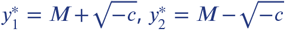 and 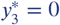. Let us show their corresponding *x*-coordinates:

- If 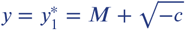, since 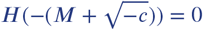 and 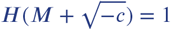,

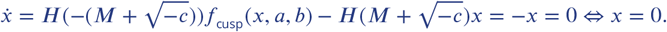 Therefore there is an equilibrium point at 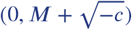.
- Also, if 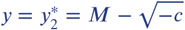, since 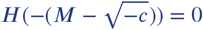and 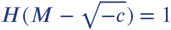,

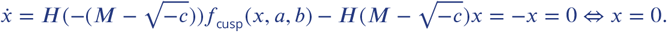 Hence, there is another equilibrium point at 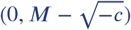.
- And if 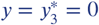, since *H*(0) = 1,

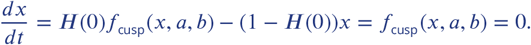 Therefore the values for the *x*-coordinates, in this case, correspond to the equilibria of a cusp catastrophe with parameters *a* and *b*. Depending on the value of the discriminant Δ = 8*a*^3^ + 27*b*^2^, the equation will have one (Δ > 0), two (Δ = 0) or three (Δ < 0) real roots. This means that, depending on the value of Δ, there will be one, two or three equilibria on the *x*-axis.

The stability of these points can be checked by looking at the eigenvalues of the jacobian matrix:

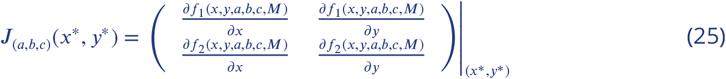

where (*x*\**, y*\*) is an equilibrium of the system in Eq. 2 in the main text, with parameters *a, b, c, M*. Since 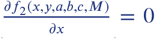, for all values of *x, y, a, b, c, M*, the Jacobian matrix is upper triangular and the eigenvalues are given by the elements on the diagonal.

In fact, the point 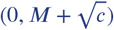 is an attractor (which will represent tertiary fate), 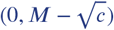 is a saddle point and the stability of the critical points on the *x*-axis depends on the value of the discriminant. Appendix 1 Figure 4 shows the possible configurations of the equilibrium points in the state space.

**Appendix 1 Figure 4.**
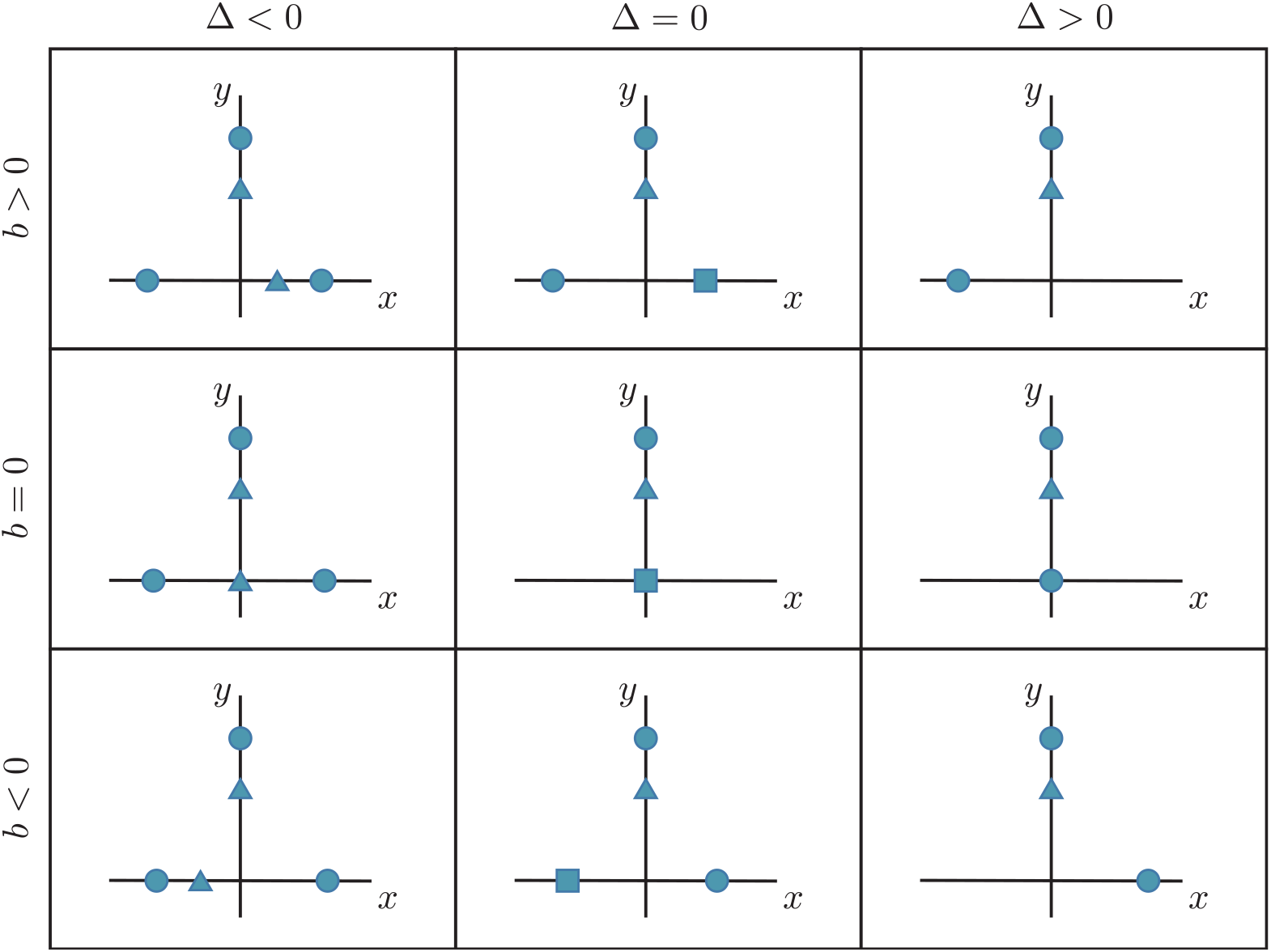
Sketch of the equilibria of the system for −*M*^2^ < *c* < 0 depending on the values of *b* and Δ. Stable equilibria are represented by circles. Saddles are represented by triangles. Degenerate steady states are represented by squares.

When *c* = 0, it follows that there is only one (degenerate) critical point at (0, ***M***), and the critical points on the *x*-axis depend on the discriminant Δ and *b*, exactly as shown before. And finally, if *c* > 0, the tertiary fate has bifurcated away and the equilibria at the *x*-axis are the only ones that remain, which again depend on Δ and *b*.

Summing up, the system can have from 5 to 1 equilibrium points in the state space depending on the values of the parameters *a, b, c*. There is a fold bifurcation point on the *y*-axis controlled by the parameter *c*, and a cusp bifurcation on the *x*-axis controlled by the parameters *a* and *b*.

## Bifurcation set of the binary flip with cusp landscape model

We well now study the bifurcation set of the system by taking advantage of catastrophe theory. The catastrophe manifold of the system, 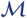, is the set of points (*x, y, a, b, c*) ∈ **ℝ**^5^ such that (*x, y*) is an equilibrium point for the system in Eq. 2 for parameter values *a, b* and *c*. In other words,

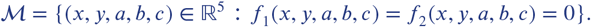

Here we will show that 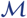 is a manifold that can be written as the union of two sets, one that corresponds to the fold bifurcation and another one that corresponds to the cusp bifurcation. Indeed, 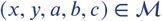 if and only if

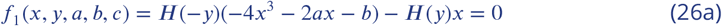

and

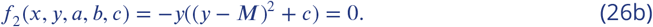

From Eq. (26b) we have that *y* = 0 or 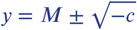. Now, if *y* = 0, from Eq. (26a):

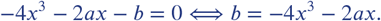

And if 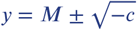, from Eq. (26a), *x* must be 0.

We can then express 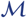 as the disjoint union of two three-dimensional manifolds, 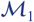 and 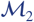, where:

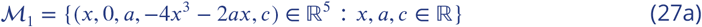

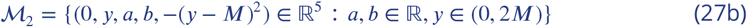

Therefore 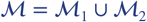 is a three-dimensional manifold in **ℝ**^5^.

In order to visualise 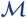 we can plot the intersection with the three-dimensional space

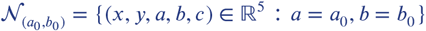

to see how the equilibrium points change when *c* changes (see Appendix 1 Figure 5).

**Appendix 1 Figure 5.**
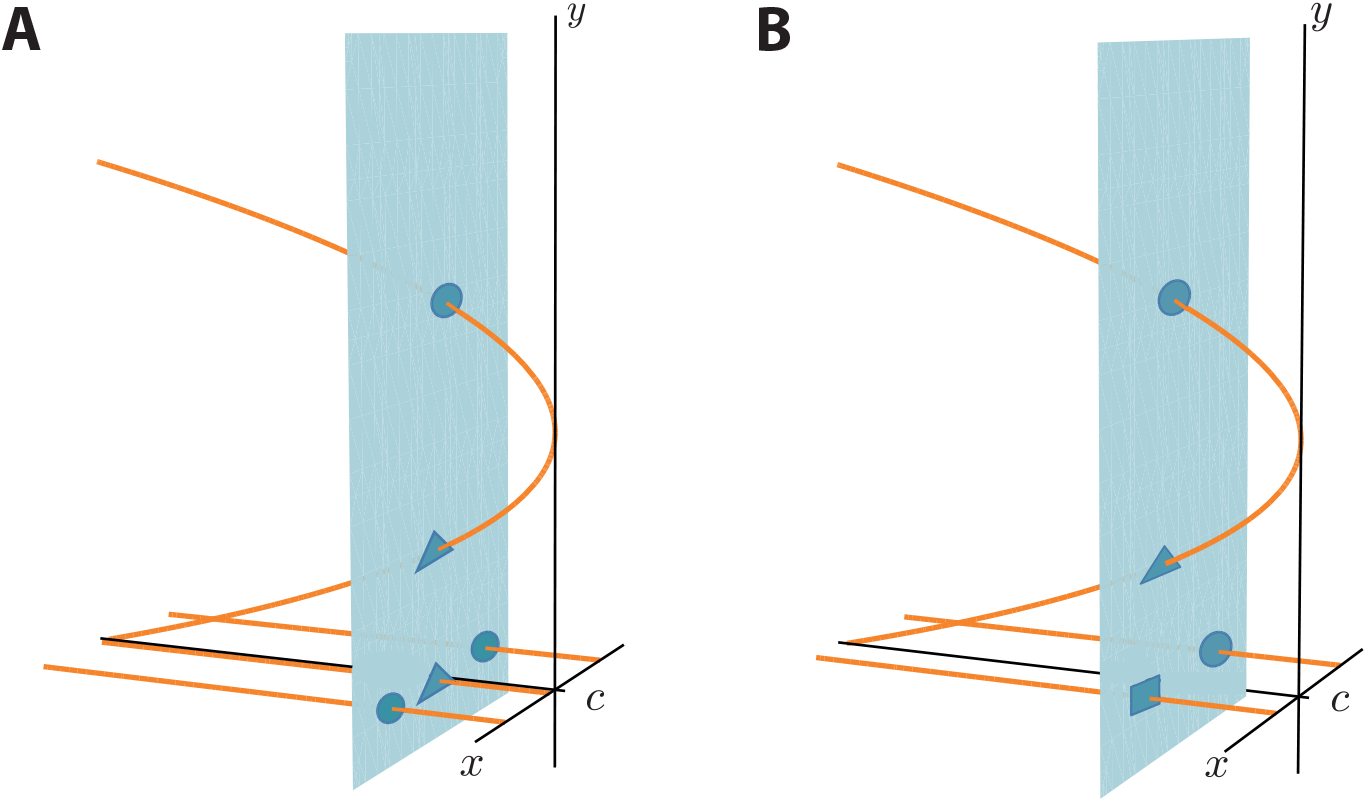
Visualisation of the catastrophe manifold of the system in Eq. 2 of the main text. In orange, 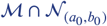 for different values of *a*_0_*, b*_0_. The plane *π* in cyan is obtained by fixing *c* = *c*_0_ < 0, in other words, *π* = {(*x, y, c*) ∈ **ℝ**^3^ : *c* = *c*_0_}. The intersection of *π* and 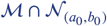 gives the equilibria of the system for parameters *a* = *a*_0_*,b* = *b*_0_*,c* = *c*_0_. If sliding the plane *π* along the *c* axis, the equilibria on the *y* axis will move along the parabola. As before, circles represent stable equilibria, triangles represent saddles and squares represent degenerate points.

The **bifurcation set** is the set of values of the parameters at which bifurcations happen, i.e. generally the parameter values at which an attractor and a saddle collide and disappear. As mentioned in the main text, this set lets us characterise regions in the parameter space of common stability. By understanding the bifurcation set of the system, we can predict which equilibrium points will be present in the state space when knowing the values of the control parameters.

We first find an expression for the **singularity set** of the system, 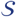, defined in earlier, which is given by:

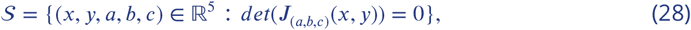

which is the subset of degenerate points of 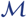.

Let us denote 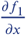 by *D_x_f*_1_ and 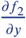 by *D_y_f*_2_. Since *det*(*J*_(*a,b,c*)_(*x, y*)) = *D_x_f*_1_*D_y_f*_2_, as explained earlier,

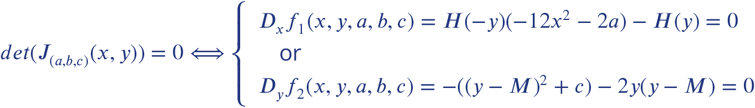

Therefore we can write 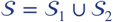 where

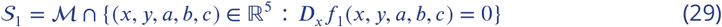

and

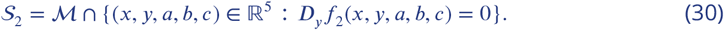

Let us understand the geometry of these sets. The set 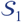 can be written as

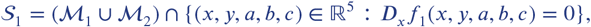

where 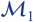 and 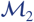 are defined in Eqs. 27a and 27b.

Applying the distributive property of the intersection over the union,

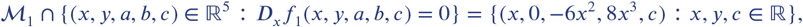

and

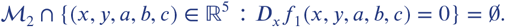

Therefore,

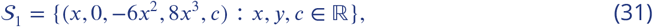

which corresponds to a cusp for every value of *c* (See Eq. 20).

On the other hand, the set 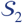 can be written as

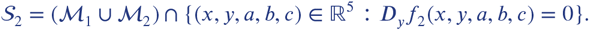

Again, applying the distributive property of the intersection over the union,

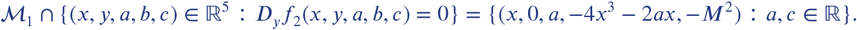

But, since we are only taking into account values of *c* > −*M*^2^, we don’t consider this set. Moreover,

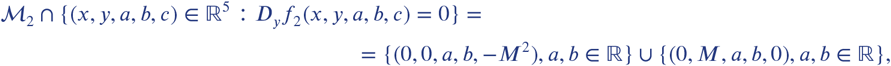

where, again, we do not consider the first subset since *c* > −*M*^2^. Therefore

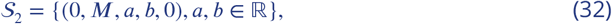

which is a plane on the parameter space.

From Eqs. (31) and (32) we can write the singularity set, 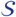, of the system as:

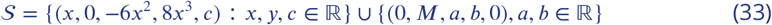

Finally, projecting 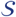 into the control space we obtain an expression for the bifurcation set, 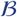:

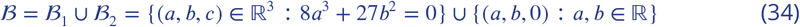

which is a two-dimensional set in the three-dimensional control space (see Appendix 1 Figure 6).

**Appendix 1 Figure 6.**
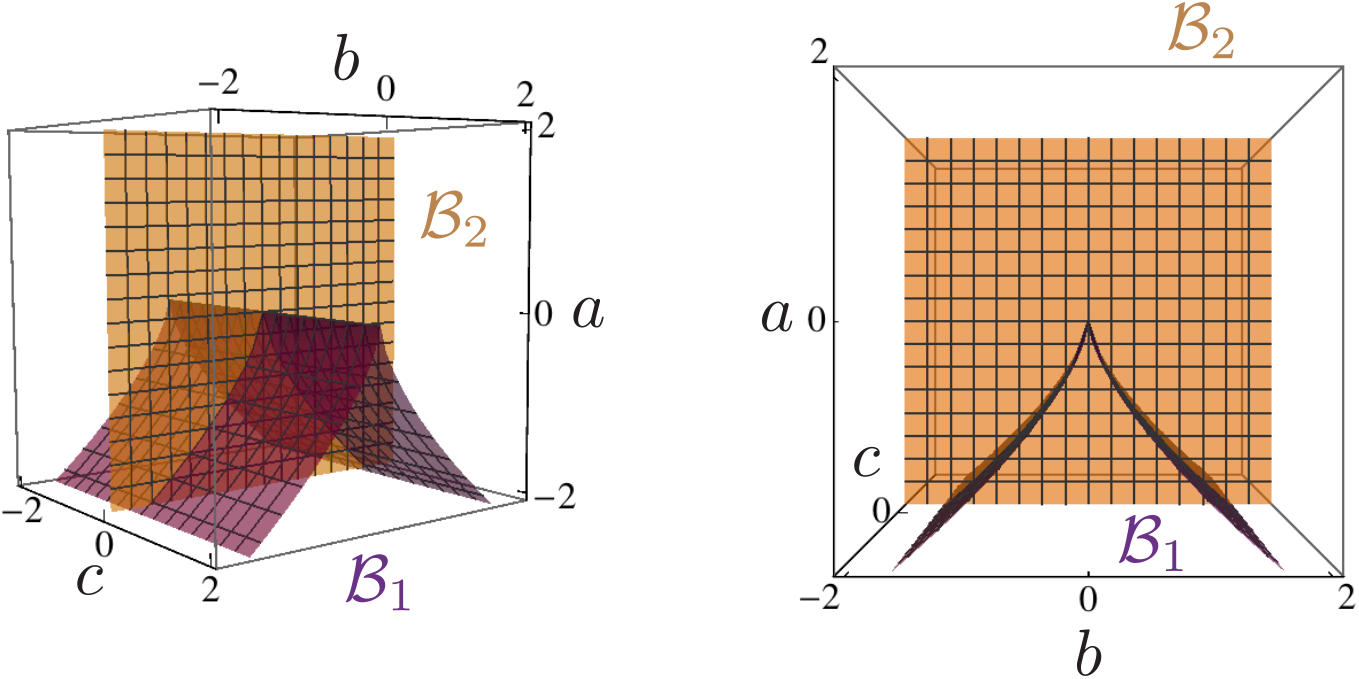
Plots showing different views of the bifurcation set given in Eq. (34). The set 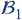 corresponds to the purple cuspoidal cylinder, while the set 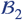 corresponds to the orange plane. For every value of *c* the system has a cusp bifurcation set with *a, b* as parameters.Also, for any value of *a, b*, the points 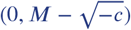 and 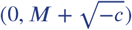 bifurcate when *c* = 0, hence the plane.

Appendix 1 Figure 7 gives a description of the different landscapes in the state space depending on the values of the control parameters (*a, b, c*). We can see that crossing the orange plane in Appendix 1 Figure 7C-E would bifurcate the points 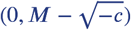 and 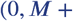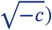 and crossing the purple surface in Appendix 1 Figure 7A-C and Appendix 1 Figure 7E-G would bifurcate the points on the *x*-axis.

**Appendix 1 Figure 7.**
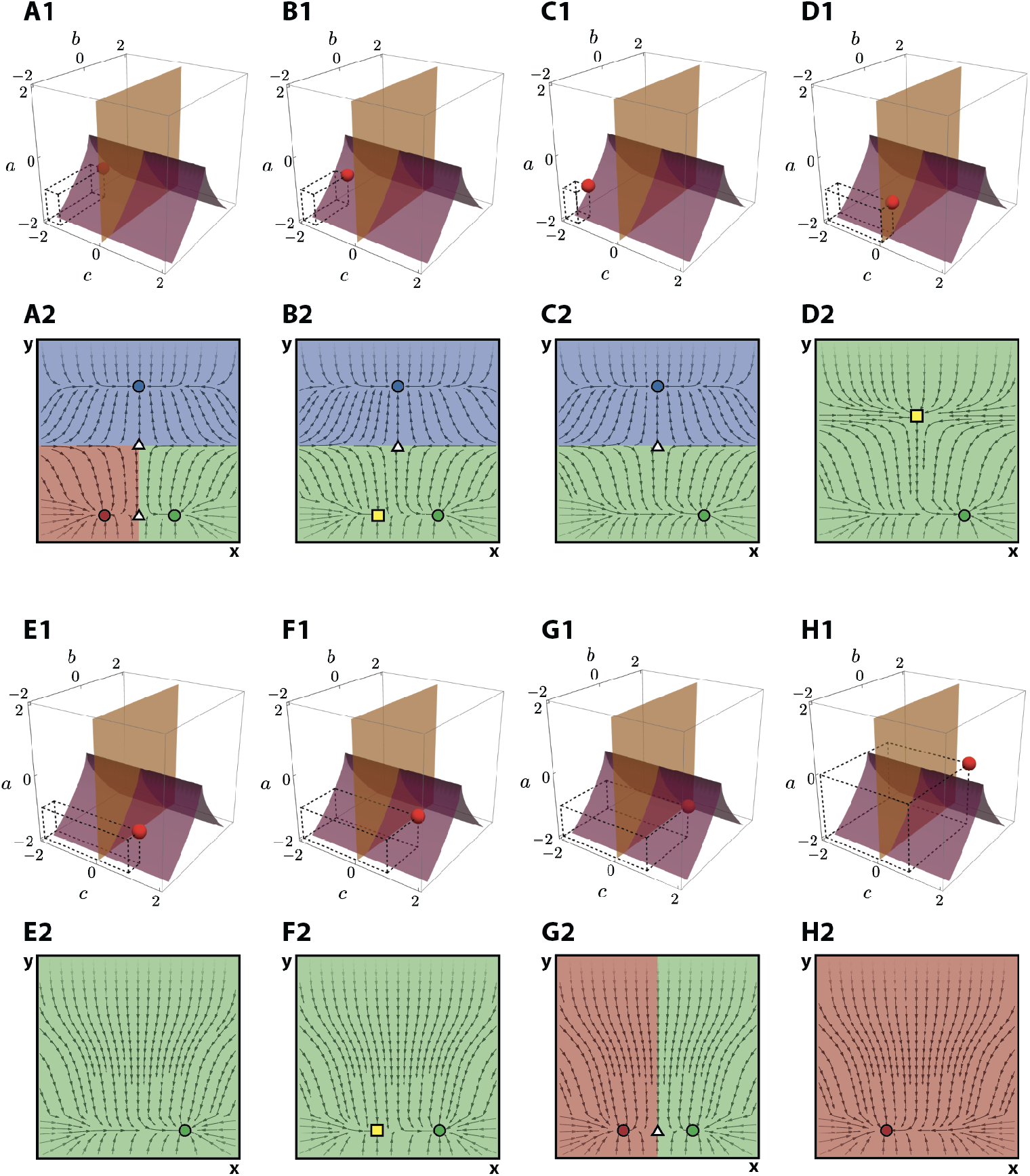
Description of the flows defined by values the control parameters (*a, b, c*) positioned in the different regions of the control space defined by the bifurcation set. (A) *a* = −1, *b* = 0, *c* = −1.5 (B) *a* = −1, 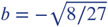, *c* = −1.5 (C) *a* = −1, *b* = −1.5, *c* = −1.5 (D) *a* = −1, *b* = −1.5, *c* =0 (E) *a* = −1, *b* = −1.5, *c* = 1 (F) *a* = −1, 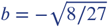, *c* =1 (G) *a* = −1, *b* = 0, *c* = 1 (H) *a* = 0, *b* = 1, *c* = 1

We have now obtained a map between the parameter space and the state space. Given a parameter value we know the exact topology of the state space. This gives us a description of how the topology of the space changes as we vary the parameters and therefore helps us to define the dynamical system that will describe the state of each VPC.

## Dependence upon the morphogens

In the previous section we have defined a dynamical system with a two-dimensional state space and a three-dimensional control space, and we have classified the possible landscapes that can appear in the state space. The goal is to take advantage of this dynamical system to define a system of differential equations such that its solution is a description of the state of each VPC during the process of differentiation.

As described in the main text, this process is controlled by two signalling pathways: induction from the anchor cell by EGF signal and lateral signalling through Notch. In consequence, the dynamics of the real system are controlled by these two parameters: level of EGF that the cell is receiving (this parameter will be denoted by *θ*_E_) and level of Notch that the cell is receiving (parameter that will be denoted by *θ*_N_). This means that the topology of the state space in the mathematical model should be related to the parameters *θ*_E_ and *θ*_N_. In other words, the parameters *a, b, c* should be written as functions of *θ*_E_ and *θ*_N_.

*θ*_E_ and *θ*_N_ can be regarded as a coordinate basis that generates a two-dimensional space. Let us call it the **signal space**. The coordinates of a point in that space will represent the values of the EGF and Notch signal that a cell receives.

Our goal is to find a correspondence between the signal space and the state space. That is to say that we aim to find which topology in the state space will correspond to which point in the signal space. And we can achieve this by finding a transformation from the (*θ*_E_, *θ*_N_) coordinate system to the (*a, b, c*) (Appendix 1 Figure 8), which in turns determines the flow in the state space.

**Appendix 1 Figure 8.**
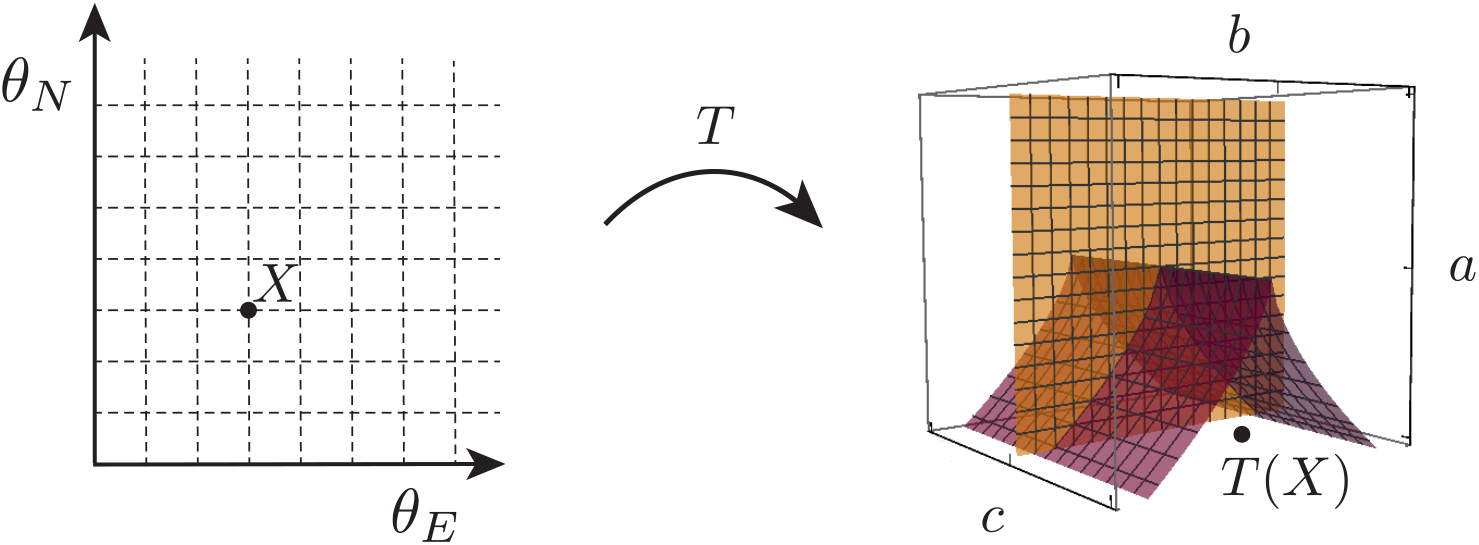
Change of coordinates from the signal space to the control space. A transformation *T* will map a point *X* in the signal space (on the left) to a point *T* (*X*) in the control space (on the right).

For simplicity, we will focus on affine transformations that map the signal space into the control space. This affine transformation, *T*, will be an embedding from the affine space **ℝ**^2^ to the affine space **ℝ**^3^, that maps the signal space into a plane in **ℝ**^3^. These transformations will be characterised by the intersection of the planes that they define and the bifurcation set 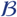, since that will determine the possible dynamics of the system.

These affine transformations will have the form:

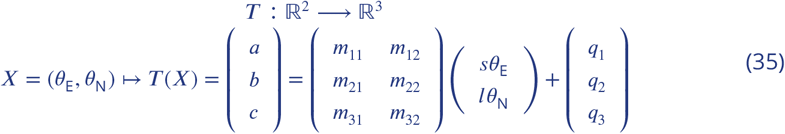

where *m_ij_* and *q_i_* define the transformation, and *s* > 0 and *l* > 0 are scaling parameters. This transformation maps a point in the signal space to a point in the plane *π_T_* = {(*a, b, c*) ∈ **ℝ**^3^ : *Aa* + *Bb* + *Cc* = *D*} in the control space where *A* = *m*_31_*m*_22_ − *m*_21_*m*_32_, *B* = *m*_11_*m*_32_ − *m*_31_*m*_12_, *C* = *m*_12_*m*_21_ − *m*_11_*m*_22_ and *D* = −*Aq*_1_ − *Bq*_2_ − *Cq*_3_.

This plane intersects the bifurcation set 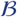 in different subsets depending on the values of *m_ij_*, and the origin of the (*θ*_E_, *θ*_N_) coordinate space in that plane will be determined by the parameters *q_i_* (see Appendix 1 Figure 9).

**Appendix 1 Figure 9.**
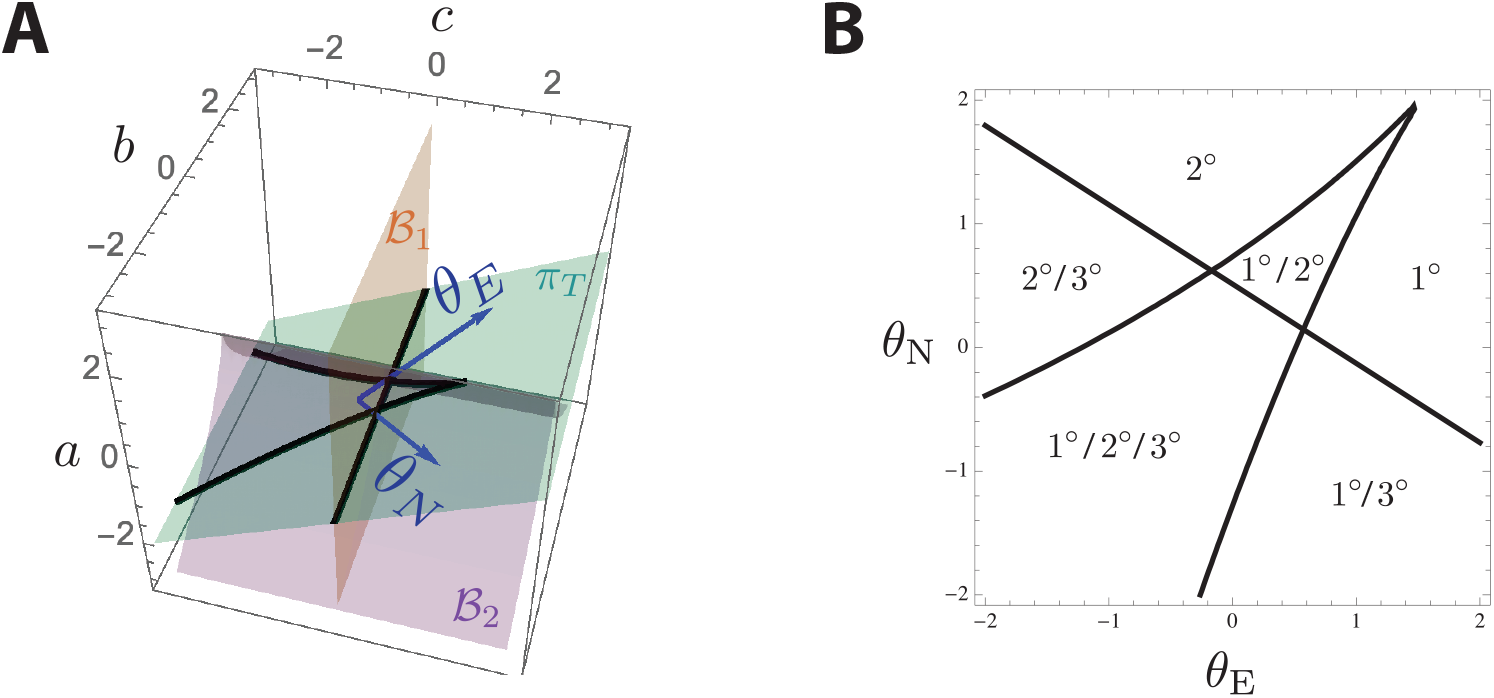
An example of linear transformation *T* from the signal space to the control space. (*A*) The figure shows, in the control space, the plane *π*_*T*_ in light blue, the bifurcation set 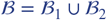 in orange and purple, the intersection 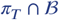 in thick black lines and the transformation of the (*θ*_E_, *θ*_N_) axis in dark blue. For simplicity, we have named this new coordinate system in the control space as the original in the signal space. (*B*) Signal space. The thick black lines correspond to the set 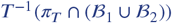. These lines define regions in the signal space that will generate different configurations of attractors in the state space. In each region, the fates which attractors will be present for a value of signals in the corresponding region are written. For *θ*_E_ =0= *θ*_N_, the system will contain the three attractors (one for each fate). For a *medium* value for *θ*_E_ and only the attractors corresponding to 1° and 2° fates will be present. For high values for *θ*_E_ and low values of only the attractor corresponding to 1° fate will be present. And, for low values for *θ*_E_ and high values of, only the attractor corresponding to 2° fate will be present.

We will assume that

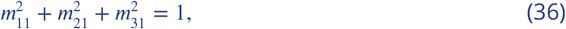

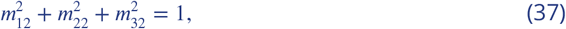

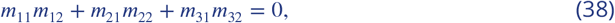

i.e., *T* is conformal (it preserves angles).

We also assume that the origin of the (*θ*_E_, *θ*_N_) coordinate system is mapped to a point in which there is tristability, this is

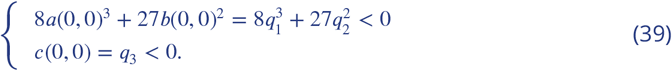

This is because of the definition of commitment of stem cells. Once a stem cell is committed, signals can be removed and the stem cell will not alter its fate. In the process of differentiation of the VPCs of a WT worm, they become specified into the three different fates (See ***Table 1***) (***Sternberg, 2005***). If we imagine the signal history of a VPC as a path in the signal space, this means that after having received different signals during their development, if the signals are switched off and the path returns to the point *θ*_E_ = *θ*_N_ = 0, the three attractors must be present for each VPC to be able to specify to their corresponding WT fates (P4.p specifies into 3°, P5.p specifies into 2° and P6.p specifies into 1°).

We could constrain the transformation so that high *θ*_E_ pushes towards monostability of 1° fate (i.e. *m*_21_ > 0) and high *θ*_N_ pushes towards monostability of 2° fate (i.e. *m*_22_ < 0).

However, we decide not to constraint the system so much, and see what the data fitting decides as the best strategy.

Finally, considering possible intersections between the plane *π_T_* and the bifurcation set, we decide to only accept transformations such that *A* ≠ 0 ≠ *C*. This is not an important constraint, since the sets *A* =0 or *C* = 0 in **ℝ**^3^ have zero measure. Moreover, by taking *A* = *ε* or *C* = *ε* with *ε* very small, one could approximate these special cases. Making this restriction allows us to easily classify the types of transformation that can be allowed. With this in mind we assume that *A* ≠ 0 ≠ *C* and, consequently, the transformations can be of two types: affine transformations such that *AC* < 0 (let us call them Type I) or affine transformations such that *AC* > 0 (let us call them Type II).

## Type I transformations

Type I transformations are such that *π_T_* intersects the plane *b* =0 on a line *a* = (*D* − *Cc*)/*A*, and the gradient −*C*/*A* is positive (see Appendix 1 Figure 10).

**Appendix 1 Figure 10.**
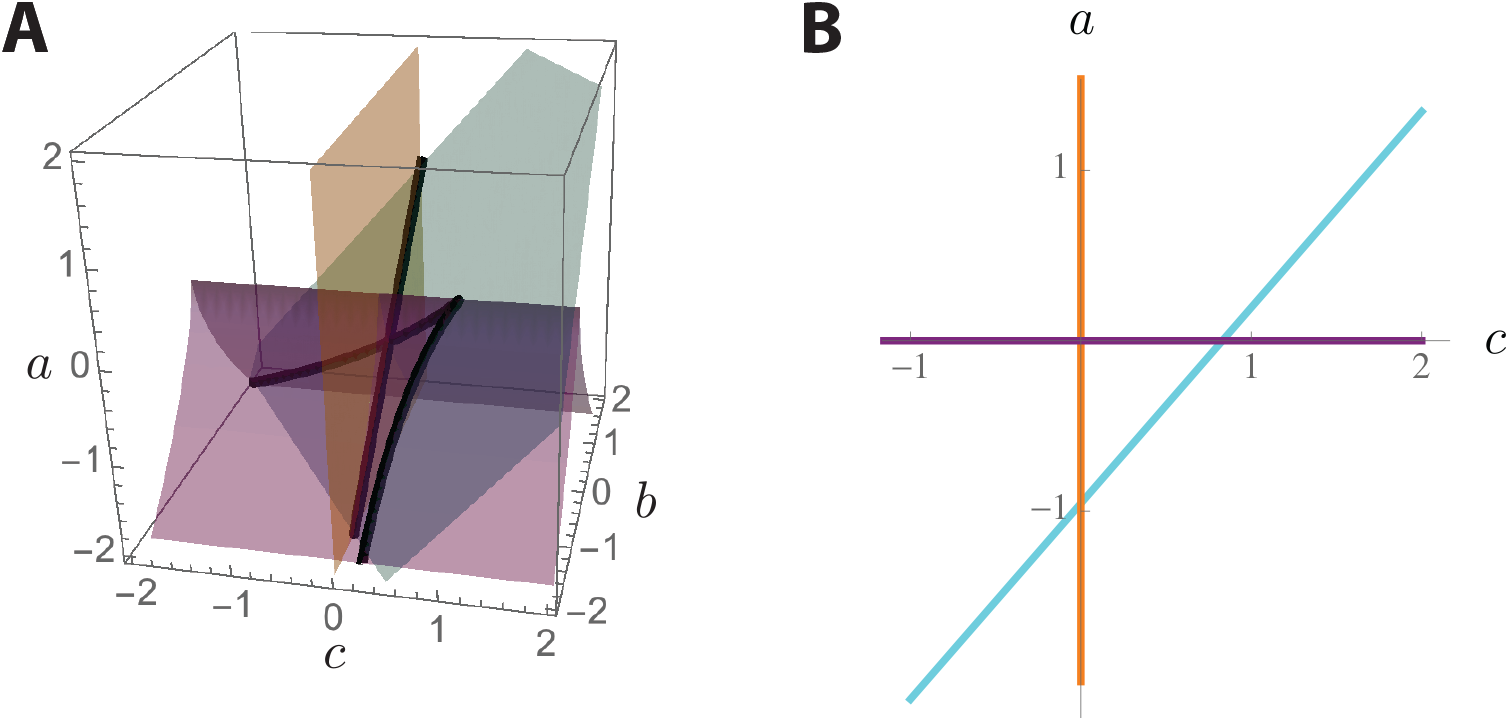
Example of the intersection 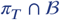 for a *T* such that *AC* < 0. (*A*) Intersection of the plane *π_T_* = {(*a, b, c*) ∈ **ℝ**^3^ : *a* − *b* − *c* = −1} and the bifurcation set 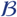. The set 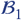 is plotted in orange, the set 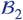 is plotted in purple and the plane *π_T_* is plotted in cyan. The intersection is marked by thick black lines. (*B*) In orange, the intersection 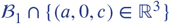. In purple, the intersection 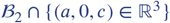. In cyan, the intersection *π_T_ cap*{(*a,* 0*,c*) ∈ **ℝ**^3^}. As we can see, the line has positive gradient.

Since a tristable region needs to be present in the intersection 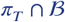, this forces

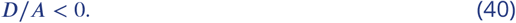

On the other hand, because of the experimental results for the Notch null with 2×EGF mutant (See experiment (5) in ***Table 1***), and *lin* − 12 gain-of-function experiments in ***Sternberg and Horvitz*** (***1989***) (which can be considered as *θ*_N_ high), we constrain the system so that *θ*_E_ high or *θ*_N_ high drive the system out of the tristable region. Therefore,

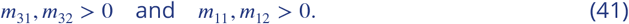

Examples of Type I transformations that satisfy constraints in Eqs. 40 and 41 are given in Figures 11.

**Appendix 1 Figure 11.**
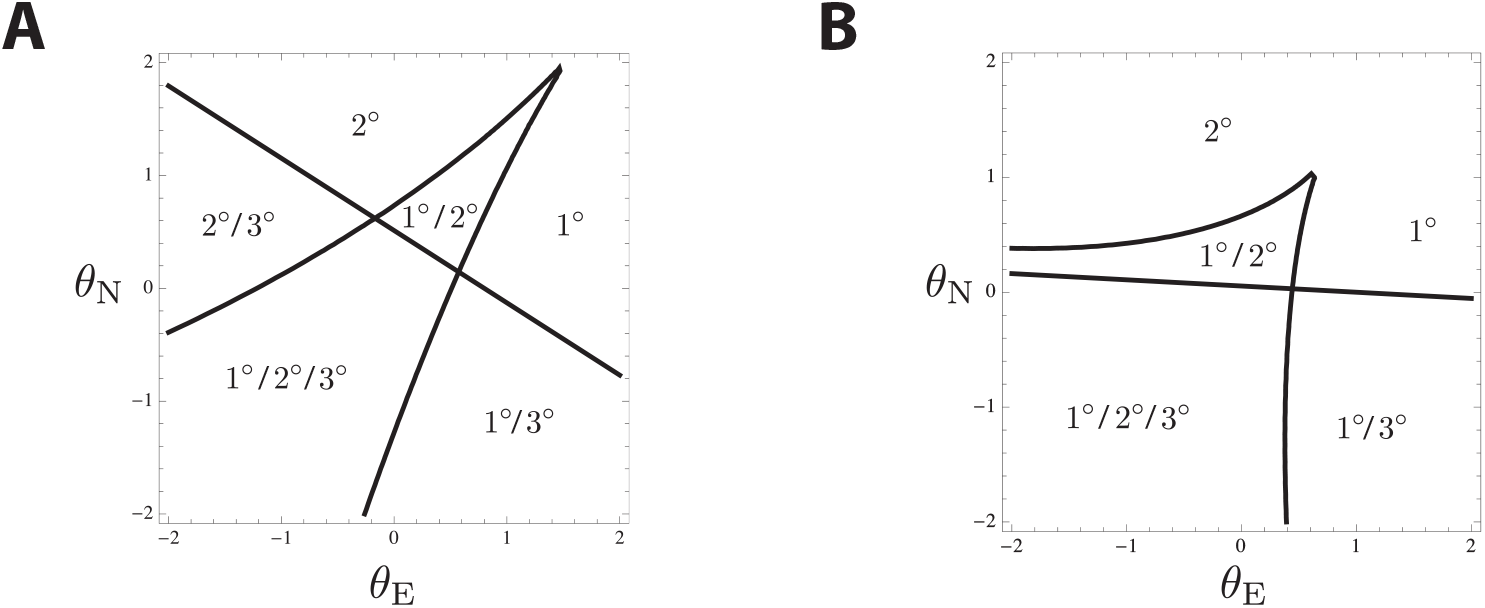
Two examples of bifurcation sets in the signal space given by the intersections 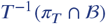 for two different *T*s of Type I satisfiying constraints in Eqs. 40 and 41. As in Appendix 1 Figure 9, different regions are labelled with the corresponding attractors.

Eqs. 36, 37, 38 and 41 again allow us to rewrite the parameters *m*_31_, *m*_22_ and *m*_32_ of the transformation as functions of the other parameters, lowering the levels of freedom. In particular:

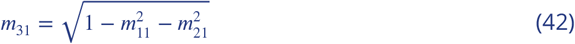

which let us write *m*_22_ and *m*_32_ as the solutions of the system

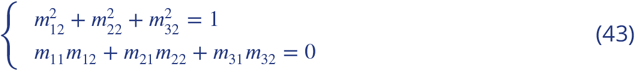

## Type II transformations

On the other hand, Type II transformations are such that *π_T_* intersects the plane *b* =0 in the control space on a line *a* = (*D* − *Cc*)/*A*, and the gradient −*C*/*A* is negative (see Appendix 1 Figure 12).

**Appendix 1 Figure 12.**
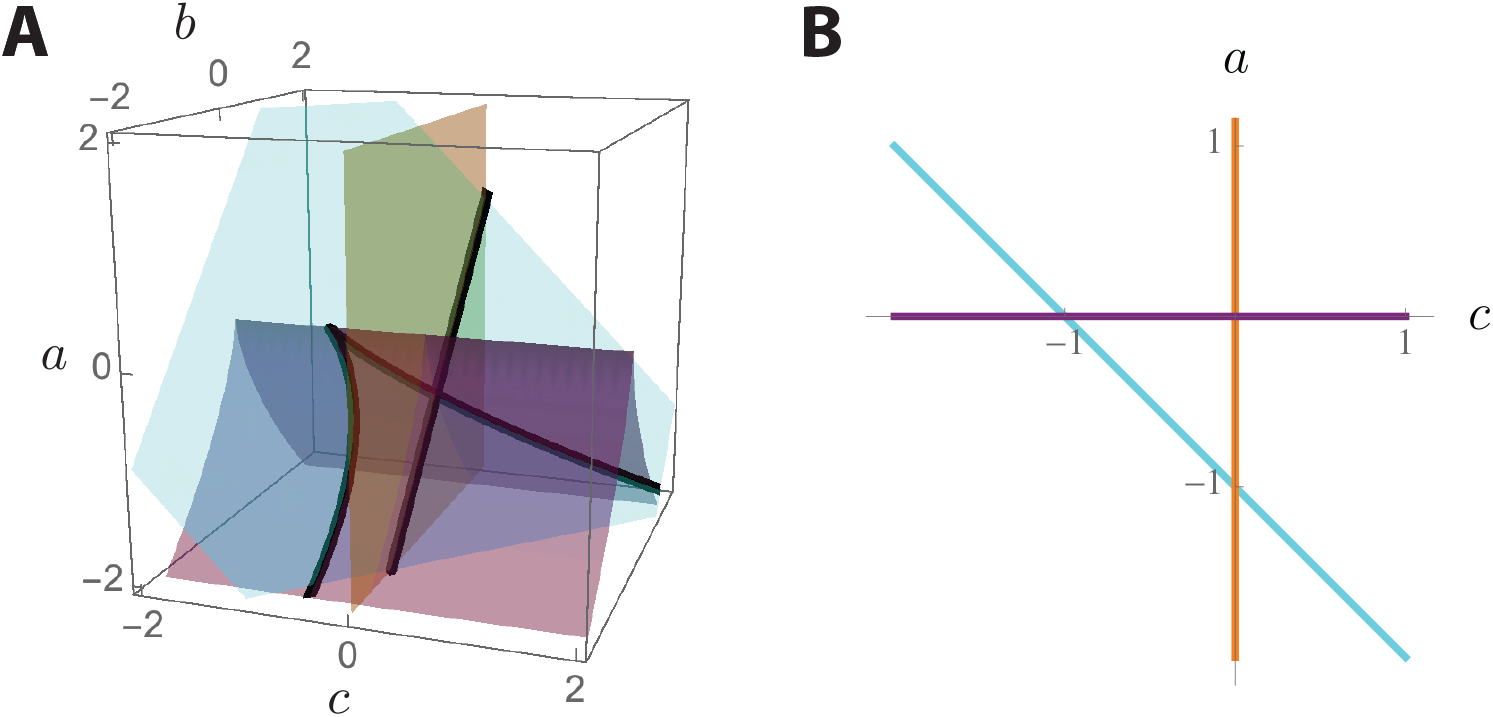
Example of the intersection 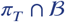 for a *T* such that *AC* > 0. (*A*) Intersection of the plane *π_T_* = {(*a, b, c*) ∈ **ℝ**^3^ : *a* + *b* + *c* = −1} and the bifurcation set 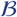. The set 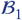 is plotted in orange, the set 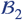 is plotted in purple and the plane *π_T_* is plotted in cyan. The intersection is marked by thick black lines. (*B*) In orange, the intersection 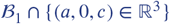. In purple, the intersection 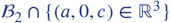. In cyan, the intersection *π_T_* ∩ {(*a,* 0*,c*) ∈ **ℝ**^3^}. As we can see, the line has negative gradient.

Since a tristable region needs to be present in the intersection 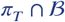, this forces

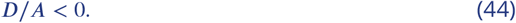

On the other hand, because of the experimental results for the Notch null with 2×EGF mutant (See experiment (5) in ***Table 1***), and *lin* − 12 gain-of-function experiments in ***Sternberg and Horvitz*** (***1989***) (which can be considered as high), we constrain the system so that *θ*_E_ high or high drive the system out of the tristable region. Therefore,

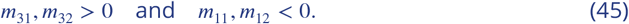

Examples of Type I transformations that satisfy the constraints in Eqs. 44 and 45 are given in Appendix 1 Figure 13.

**Appendix 1 Figure 13.**
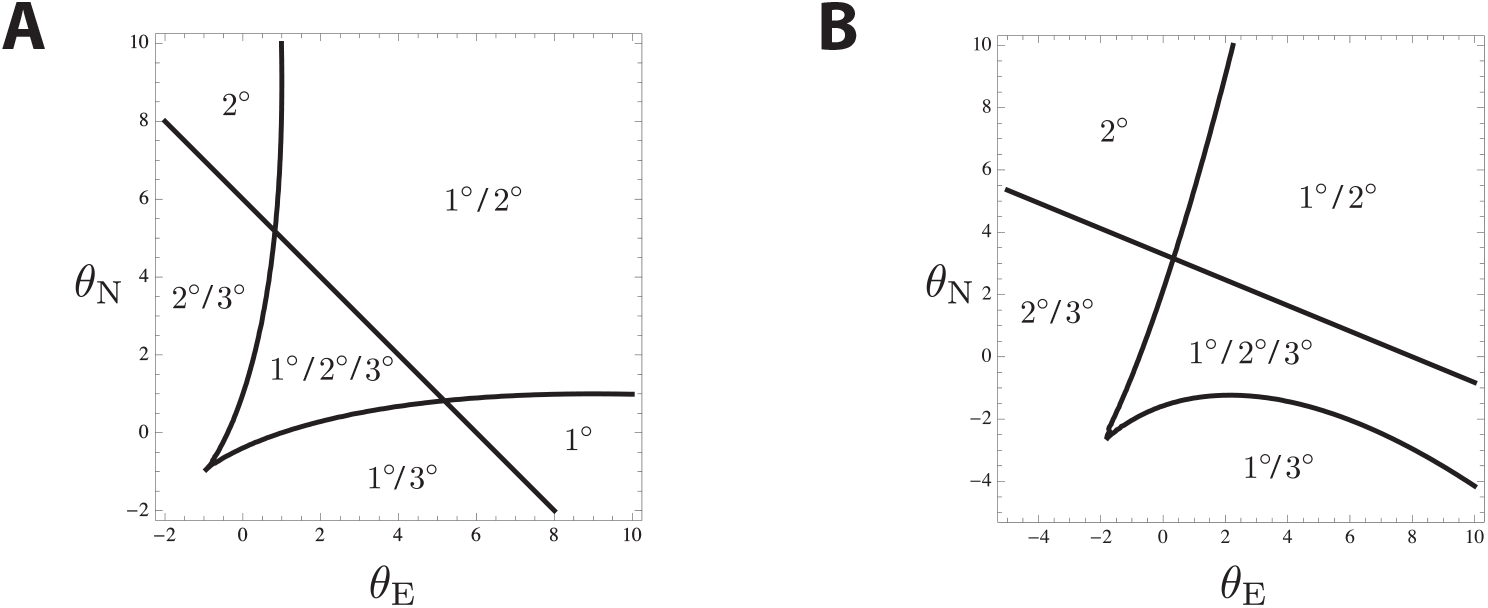
Two examples of bifurcation sets in the signal space given by the intersections 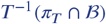 for two different *T*s of Type I satisfiying constraints in Eqs. 44 and 45. As in Appendix 1 Figure 9, different regions are labelled with the corresponding attractors.

Eqs. 36, 37, 38 and 45 allow us to rewrite three parameters of the transformation as functions of the other parameters, lowering the levels of freedom. In particular:

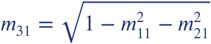

which let us write *m*_22_ and *m*_32_ as the solutions of the system

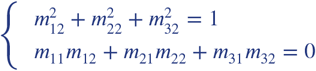

Here we focus on Type I transformations, as they fit the data better. Type II transformations failed to reproduce some EGF overexpression data.

Considering the system defined in Eq. 2 in the main text, and the linear transformations with the corresponding constraints, we can now present the proposed model that will describe the process of differentiation of the VPCs.

## Model implementation and numerical simulations

So far we have developed a model that describes the process of differentiation of a single VPC in time, given the signals EGF and Notch. However, the biological system is formed by 6 VPCs (P3-8.p), that interact with each other. However, taking into account that P3.p normally fuses with the hypodermis and that the pattern is usually symmetric around the AC, we will model the development of 3 VPCs: P4.p, P5.p and P6.p, where P6.p is the closest to the AC. This means that the mathematical model will be comprised of three trajectories, each corresponding to a VPC, moving on their corresponding landscapes which shapes will depend on the signals that each cell receives. Our proposed model is therefore described by the following set of equations, with (*x*_1_*, y*_1_) being the coordinates representing the state of differentiation of P4.p, (*x*_2_*, y*_2_) the coordinates representing the state of P5.p and (*x*_3_*, y*_3_) the coordinates representing the state of P6.p:

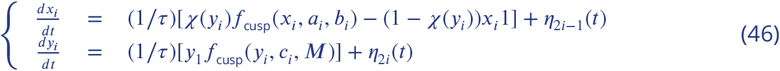

where, 1/*τ* is a time constant, *f*_cusp_ (*x, a, b*) = −4*x*^3^ − 2*ax* − *b*, *f*_fold_ (*y, c, M*) = (*y* − *M*)^2^ + *c* are the functions explained above and, in order to work with smooth functions, we changed *H*(*y*) to *χ*(*y*), a sigmoidal function of the form:

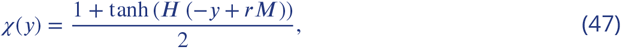

where we choose *H* = 25 and 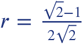. A white noise *η_i_* is also added to the system, with variance given by a parameter 2*σ_dif_*, to account for intrinsic variability:

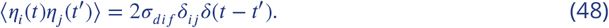

The values of the control parameters for each cell are given by:

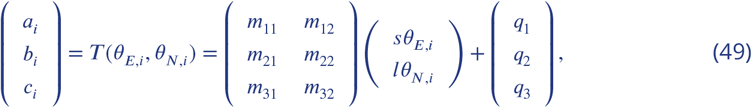

where, as explained above, *T* is either of Type I, satifying the constraints in Appendix 1 Table 1, where the definitions of *A, C* and *D* and the constraints were explained before.

**Appendix 1 Table 1.**
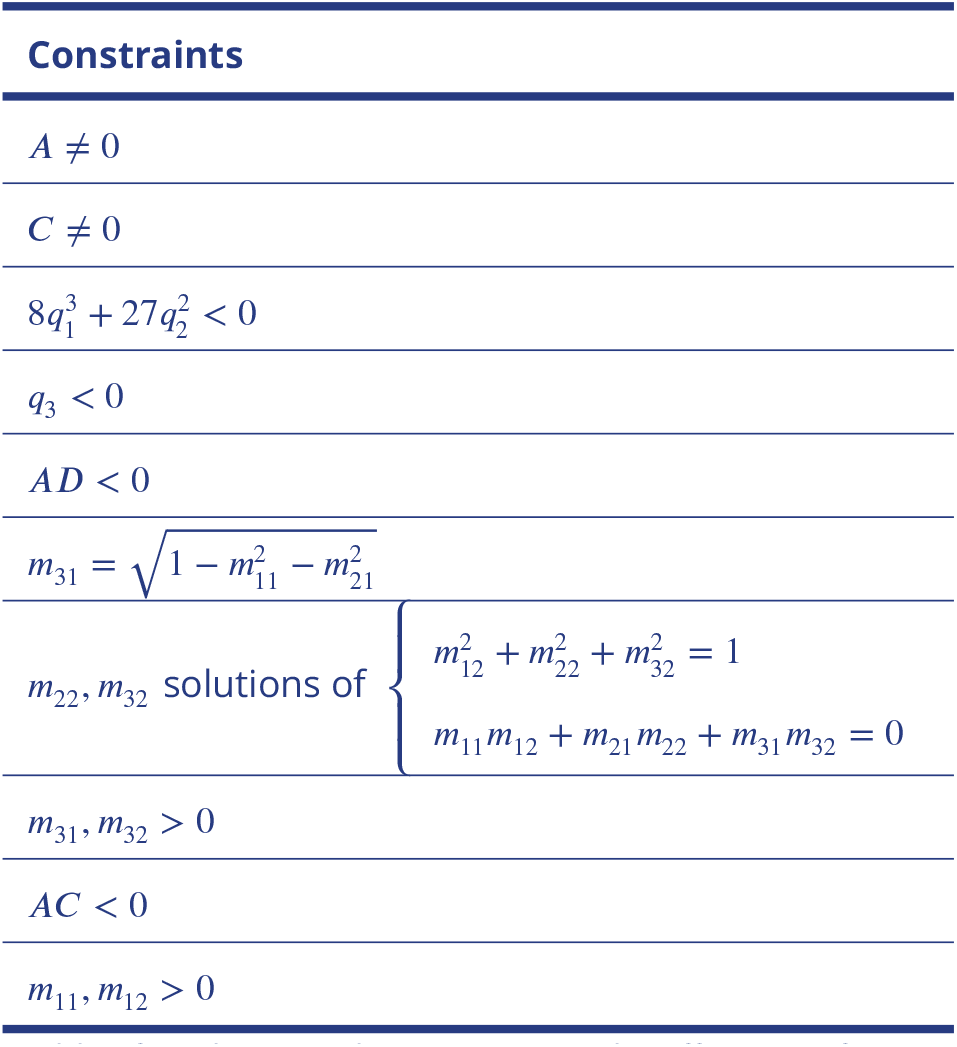
Table of mathematical constraints on the affine transformation *T* of Type I from the signal space to the control space.

Following the approach by ***Corson and Siggia*** (***2012***, 2017), we assume that the difference of EGF signal between consecutive cells is regulated by a scaling parameter *γ*, which derivation follows from the diffusion of EGF morphogen (*γ* < 1). However, we also model the increase of EGF signal in time as a monotonous increasing function *σ*(*t*):

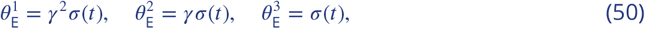

where the parameter *s* is a scaling factor and the function *σ*(*t*) is given by:

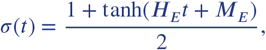

where *H_E_* and *M_E_* are parameters to be fitted.

We also define the parameter to be proportional to the sum of the autocrine Notch signal of the cell (signal produced by the cell itself) and the paracrine Notch signal received by the cell (produced by the neighboring cells). The autocrine signal is multiplied by a parameter *α* which parametrises the relative importance of autocrine and paracrine signalling. We also take into account that 1° fated cells downregulate Notch receptor lin-12, and therefore downregulate the Notch signal received, as suggested in ***Shaye and Greenwald*** (***2002***). This downregulation is related to the cell’s production of Notch signalling, scaled by a parameter *l_d_* which defines the strength of such downregulation. Therefore, we define the levels as

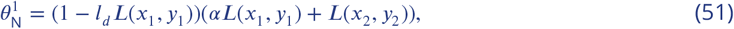

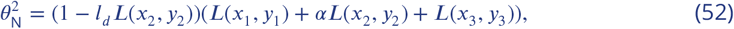

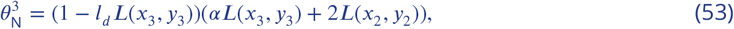

where the parameter *l* is a scaling factor. The function *L* represents the Notch signal emitted by a cell, which depends on the current state of the cell, and it is chosen as:

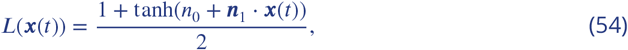

where *n*_0_ and ||***n***_1_|| are constants. *L* should be increasing as the state of the cell approaches the attractor corresponding to 1° fate, therefore we fit the *y*-coordinate of ***n***, 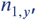, and fix its *x*-coordinate to be 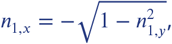, so that *L* increases as a cell moves toward the basin of attraction of 1° fate.

Taking into accounts the constraints of the linear transformation discussed above, the development of the VPCs is then defined by a 6-dimensional system of stochastic differential equations depending on the twenty parameters in Appendix 1 Table 2.

**Appendix 1 Table 2.** Table of model parameters. Table containing the landscape parameters that map the signaling values into the shape of the landscape and 15 signaling parameters that define the signaling profile that each cell is exposed to in time.

|  | Parameter | Description |
| --- | --- | --- |
| Landscape | $1/\tau$ | Time constant |
| | $M$ | Position of the saddle point between 3° fate and 1°-2° fates |
| | $m_{11}$ | Linear transformation between signal space and control space |
| | $m_{12}$ | Linear transformation between signal space and control space |
| | $m_{21}$ | Linear transformation between signal space and control space |
| | $q_1$ | Position origin signal space in control space, linear transformation |
| | $q_2$ | Position origin signal space in control space, linear transformation |
| | $q_3$ | Position origin signal space in control space, linear transformation |
| | $s$ | EGF signal scaling constant |
| | $l$ | Notch signal scaling constant |
| Signals | $\gamma$ | EGF diffusion |
| | $H_E$ | Gradual EGF increase in time |
| | $M_E$ | Gradual EGF increase in time |
| | $l_d$ | Stregnth of Notch inhibition in 1° fated cells |
| | $\alpha$ | Autocrine Notch signaling strength |
| | $n_0$ | Notch signal sigmoidal function |
| | $n_1$ | Notch signal sigmoidal function |
| | $n_{1,y}$ | Notch signal sigmoidal function |
| | $\lambda_E$ | Exponential decay of EGF |
| | $\lambda_N$ | Exponential decay of Notch signal |
| | $\sigma_{dif}$ | White noise variance |

The model and following steps are implemented in Matlab, and the code is provided in the following github repository: https://github.com/ecamacho90/VulvalDevelopment

## Initial condition

In order to perform a simulation, first, it is necessary to choose an initial condition that will represent the state of the VPCs at the start of the experiment. We assume that the VPCs are initially equivalent and that their initial condition should not depend on the signals they will receive. Since in the absence of signals they take the 3° fate, we assume that their initial condition should lie in the basin of attraction of the attractor corresponding to the 3° fate.

It is also reasonable to incorporate some variability in the initial state, so we decide to choose as initial condition of the full system, the stationary distribution around the equilibrium of the following system of SDEs

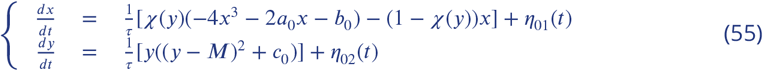

where *a*_0_ = *q*_1_, *b*_0_ = *q*_2_, *c*_0_ = *q*_3_ are obtained from equation (49) by setting *θ*_E_ =0 and *θ*_N_ =0 and *1*_01_ and *1*_02_ represent independent white noises with variance 2*σ_dif_*.

As initial condition of the system in Eq. 55 we choose a normal distribution centered at (*x*(0)*, y*(0)) = (0*,M*) with zero covariance matrix, to make sure that the initial condition stays in the basin of attraction of the tertiary fate.

We decide to approximate such stationary distribution numerically. In order to do that, we considered two approaches. As a first attempt, we approximated such distribution by running *N* simulations (*N* = 100, 1000) for a long enough period of time by taking advantage of the Euler-Maruyama method. The stationary distribution was approximated by the distribution of the end points of those *N* simulations. As a second approach, we approximated the solution of the SDE based on a similar method to derive the Linear Noise Approximation (LNA) ***Fearnhead et al.*** (***2014***); ***Wallace*** (***2010***), and computed the solution for a long enough period of time (*t* ∈ (0, 10)). In order to get the same accuracy with the two methods, the first approach took twice as much time as the second approach, so we decided to use the second one.

## Simulation procedure

Eq. 46 is solved by using Euler-Maruyama (***Higham, 2001***; ***Kloeden and Platen, 1992***; ***Wilkinson, 2011***). We draw *N* initial conditions from the initial distribution computed as we explained before and simulate *N* random walks with the Euler Maruyama method. The system in Eq. 46 is simulated from *t* = 0 to *t* = 1, with time step *dt* = 0.005, with signals active, to account for the competence period in which the VPCs respond to signals. Then, the system is continued from *t* = 1 to *t* = 3, with *dt* = 0.005, with signals off to account for the post-competence period. We consider this post-competence period because, in order for cells to be specified, they should keep their fates even if they do not receive any signals. During this post-competence period we consider an exponential decay of the signals (we found anomalous solutions if we instantaneously set *θ*_E_ = *θ*_N_ = 0, and it is also more biologically reasonable). Therefore, during the post-competence period, we replace *s* and *l* by:

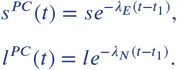

The parameters *λ_E_* and *λ_N_* are fitted but constrained so that, at *t* = 3, *s^PC^* (3), *l^PC^* (3) ≈ 0. At time *t* = 3, for each one of the *N* simulations, the fates of the three VPCs are scored as will be explained in the following section. The simulated outcome is given by the proportion of times that each cell took each fate. Therefore, the simulated outcome can be sumarised in a matrix

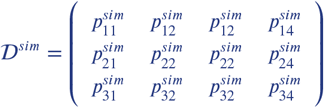

where for *j* = 1, 2, 3 the value 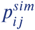 is the proportion of times, in the *N* simulations, that P(*i*+3).p took fate *j*°. For *j* = 4 the value 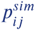 is the proportion of times, in the 150 simulations, that P(*i* + 3).p could not be assigned a fate (See next subsection for more details). We checked the sensitivity of 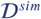 with respect to the number of simulations and we found that *N* = 150 gave best results.

## Fate assignment

Given one simulation of the system, the goal is to compute the fate of each cell. As explained above, we score the fates at time *t* = 3. At that time, the signals are switched off for the three VPCs, so they lie on the same fixed flow defined by the system of equations:

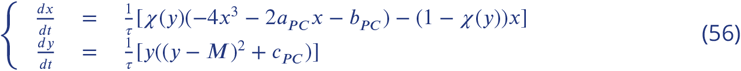

where *χ*(*y*) is defined in Eq. 47, *a_PC_* = *q*_1_, *b_PC_* = *q*_2_, *c_PC_* = *q*_3_, such that it contains the three attractors (one for each fate), as explained above.

Depending on the value of *b_PC_* = *q*_2_, the basins of attraction will look like the ones represented in Appendix 1 Figure 14.

**Appendix 1 Figure 14.**
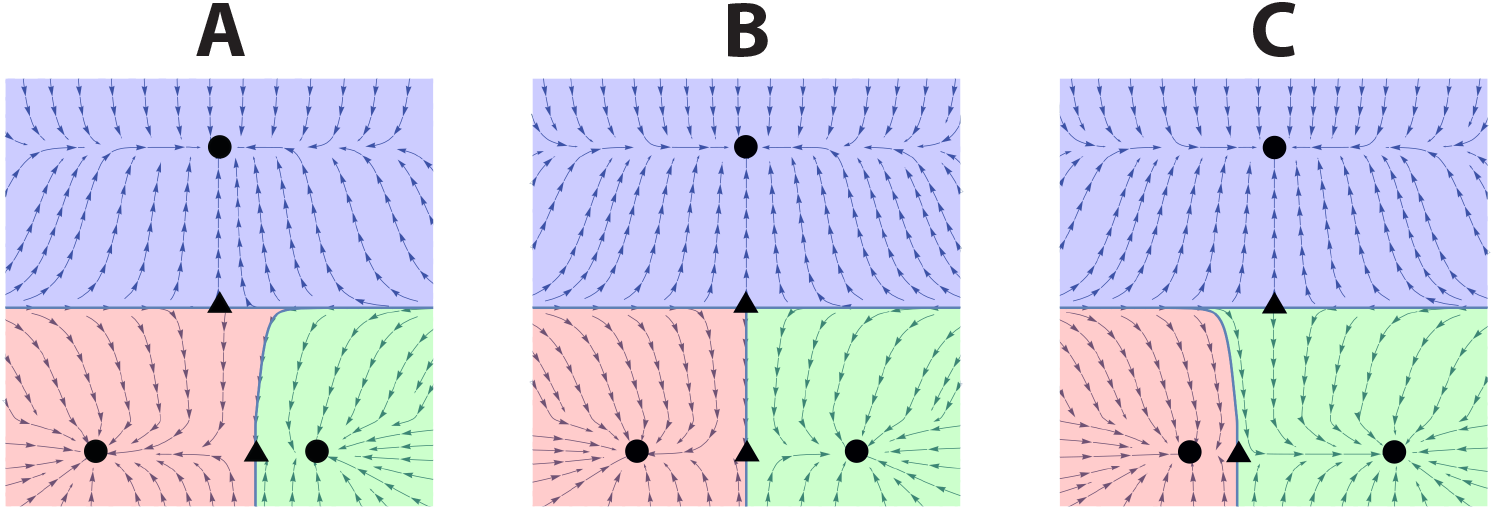
Possible landscapes for post-competence period for different values of *q*_2_, when the control parameters take values *a_PC_* = *q*_1_, *b_PC_* = *q*_2_, *c_PC_* = *q*_3_. Arrows represent the flow defined by the system in Eq. 56. Basins of attraction are coloured blue, green and red from attractors corresponding to the 3°, 2° and 1° fates, respectively.

In order to assign a fate to a VPC in one simulation, we find the basin of attraction in which the trajectory lies at time *t* = 3, and allocate the corresponding fate.

Let us call (*x_i_, y_i_*) coordinates of a cell at time *t* = 3 for a particular simulation. In order to find the basin of attraction in which it lies we do the following:

1. Suppose 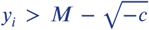. As we see in Appendix 1 Figure 15, the basin of attraction corresponding to the 3° fate is the region of the state space defined by 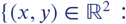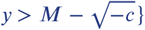, where 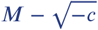 is the *y*-coordinate of the yellow saddle in Appendix 1 Figure 15. Therefore, if 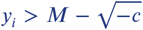, then we assign the 3° fate to this VPC in such a simulation.
2. If 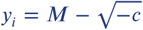, then the point lies on the boundary between two basins of attraction. We consider that we cannot assign a fate to the VPC in this case.
3. If 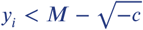 then the point lies in the basin of attraction of either 1° or 2° fate.

a. If *q*_2_ = 0, the saddle between the red and green regions in Appendix 1 Figure 14 has coordinates (0, 0) and it is half way between the attractors corresponding to fates 1° and 2°. In this case, the basin of attraction corresponding to 1° fate is defined by the set of points with *x* < 0, 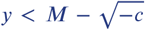. Moreover, the basin of attraction corresponding to the 2° fate is defined by the set of points given by *x* > 0, 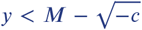. Consequently, we assign fate 1° (resp. 2°) to such a VPC if *x_i_* < 0 (resp. *x_i_* > 0).
b. If *q*_2_ ≠ 0, we observe that the stable manifold of the saddle between the red and green regions, which serves as boundary between the two basins of attraction, bends towards the closer of the two attractors (see Appendix 1 Figure 14). Consider the case *q*_2_ < 0 (similarly if *q*_2_ > 0). In this case, the landscape would look like the leftmost landscape in Appendix 1 Figure 14. Let 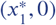 be the coordinates of the attractor corresponding to the 1° fate, 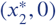 be the coordinates of the attractor corresponding to the 2° fate, 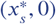 be the coordinates of the saddle determining the boundary between the basins of attraction of 1° and 2° fates, and 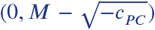 be the coordinates of the saddle determining the basin of attraction of the 3° fate. We assign the fate as follows:

i. If 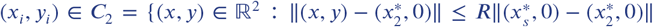, i.e. (*x_i_, y_i_*) is in the dark green circle arround the attractor corresponding to 2° fate in Appendix 1 Figure 16), we assign the fate 2°.
ii. If 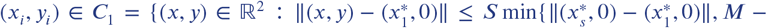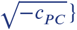 i.e. (*x_i_, y_i_*) is in the dark red circle around the attractor corresponding to 1° fate in Appendix 1 Figure 16), we assign fate 1°.
iii. Otherwise, we solve the system of ordinary differential equations in Eq. 56 until the trajectory crosses either *C*_1_ or *C*_2_, assigning the corresponding fate.

**Appendix 1 Figure 15.**
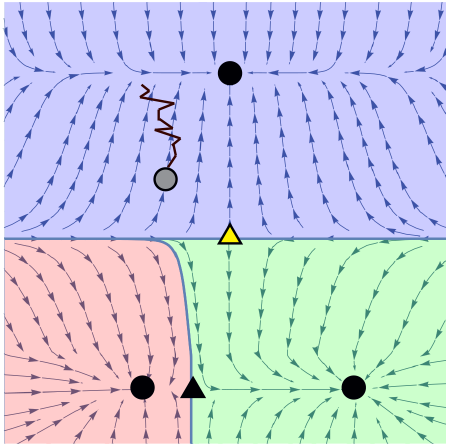
Example of an assignment of the fate 3 to a VPC in a simulation. Arrows represent the flow defined by the system in Eq. 56. Basins of attraction are coloured blue, green and red from attractors corresponding to the 3°, 2° and 1° fates, respectively. The end point (gray circle) of a simulated trajectory of a VPC during the post-competence period (black path) lays in the basin of attraction of 3° fate (blue region).

**Appendix 1 Figure 16.**
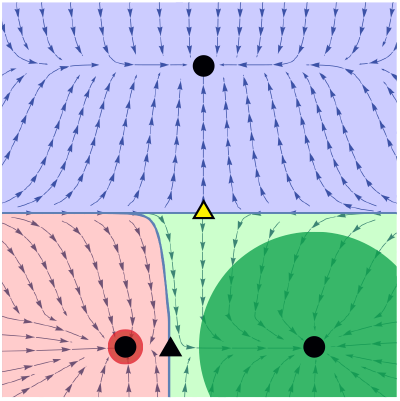
Assignment of 1° or 2° fates when *q*_2_ < 0. Arrows represent the flow defined by the system in Eq. 56. Basins of attraction are colored blue, green and red from attractors corresponding to 3°, 2° and 1° fates, respectively. Dark green region corresponds to the set 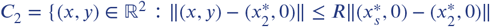 Red green region corresponds to the set 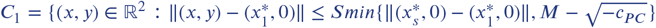

Where we take *R* = 0.9 and *S* = 10−^3^.

## Simulation of mutants

If we call

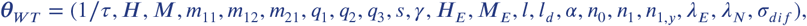

the parameter vector that represents WT conditions in the model, we can simulate all the mutants in ***Table 1*** by changing the corresponding parameters the following way.

## *let-23* mosaic

In this mutant worm, there are no EGF receptors in P5.p. Therefore we simulate this mutant with the same parameter vector Θ_*WT*_ as the WT but we change the parameter that controls the EGF signal for P5.p. In particular, we set *s* =0 only in P5.p for this mutant.

## Half dose of *lin-3*

In this case, the EGF ligand is reduced by half with respect to the WT case in all the VPCs. We can translate this into our model by taking the parameter *s*/2 instead of *s* in the WT parameter vector.

## Half dose of *lin-12*

In this case, the Notch receptor is reduced by half with respect to the WT case in all the VPCs. To do this we reduce by half the WT parameter *l*.

## Notch null, 2× WT EGF

In this mutant, the VPCs lack the Notch receptor, and the EGF that they receive is twice as much the EGF signal that they receive in the WT worm. Taking this into account, we simulate this mutant by doubling *s* and setting *l* to zero in the WT parameter vector.

## No notch signalling, WT EGF

As in the previous case, the VPCs do not receive any Notch signal because they lack the Notch receptor. However, in this case, EGF signal is the same as in the WT case. We can translate this into our model by setting *l* to zero in the WT parameter vector.

## Excess EGF

These mutants are modelled as a multiplicative increase in EGF signal. For the mutant JU1100, the VPCs receive a higher EGF signal but the fold change is unknown. We add a parameter *s_eE_* > 0 to the model, that will represent the fold change of EGF mutant in this mutant with respect to the WT case. Therefore we multiply *s* by *s_eE_* in the WT parameter vector. On the other hand, in the case of JU2113 it is known that the EGF level is increased by 1.25-fold, so for this mutant we multiply *s* by 1.25.

## Reduced Notch

These mutants are modelled as a multiplicative decrease in Notch signal. In particular, we consider a reduction of 0.4× *l*, as explained in the main text.

## Excess EGF & ectopic Notch

Excess EGF is modelled as explained above, while ectopic Notch is modelled as an additive increase in the signal, where now the *θ*_N_ described in Eqs. 51,52,53 become 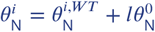, where 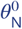 is constant and equal to 0.12.

## Reduced EGF

Reduced EGF is modelled as a multiplicative decrease in EGF signal. For the mutants considered, the VPCs receive a low EGF signal but the fold change is unknown. We consider a parameter 1 > *s_rE_* > 0 to the model, that will represent the fold change of EGF mutant in this mutant with respect to the WT case. Therefore we multiply *s* by *s_rE_* in the WT parameter vector. In our case, *s_rE_* = 0.36.

## AC ablation mutants

These mutants correspond to experiments in which the AC is ablated (i.e. EGF signal is removed) at different stages of the development of the worm. If we simulate the competence time of the WT case from *t* =0 to *t* = 1, we simulate the competence time of these mutants by taking the parameter vectors:

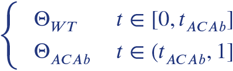

where Θ_*ACab*_ is obtained by setting 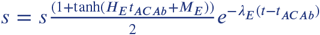 in the WT parameter vector, meaning that EGF is removed so it decays exponentially from the point to which it reached at *t_ACAb_*, and *t_ACAb_* represents the developmental stage. Since the numerical timing of each developmental stage is not available, we first followed the approach taken by ***Corson and Siggia*** (***2012***, 2017) and assumed that the AC ablation times were uniformly distributed in [0.2, 0.8], as shown in Appendix 1 Table 3. However, after fitting the data corresponding to the stages L2 Lethargus and DU divided, the data suggested a different correspondence, as shown in Table Appendix 1 Table 3. This change is equivalent to a change in the EGF monotone function, *σ*(Ψ(*t*)), that we assumed to be sigmoidal (see Appendix 1 Figure 17) and is, in fact, is consistent with data in ***Milloz et al.*** (***2008***), where the expression of an EGF pathway transcriptional reporter *egl-17::CFP* is measured at some developmental stages. It could also be achieved by locally changing the flow around the saddle. However, we find the change in the EGF function is not very significant, and due to the lack of data for EGF dynamics, we accept this change.

**Appendix 1 Table 3.** Table of correspondance between developmental stages and modelling time.

| Developmental stage | Initial $t_{ACAb}$ | $\Psi(t_{ACAb})$ from fitting |
| --- | --- | --- |
| L2 lethargus | 0.2 | 0.2 |
| Early L3 | 0.32 | 0.34 |
| DU divided | 0.44 | 0.40 |
| VU divided | 0.56 | 0.45 |
| 3° divided | 0.68 | 0.49 |
| 2-cell stage | 0.8 | 0.78 |

**Appendix 1 Figure 17.**
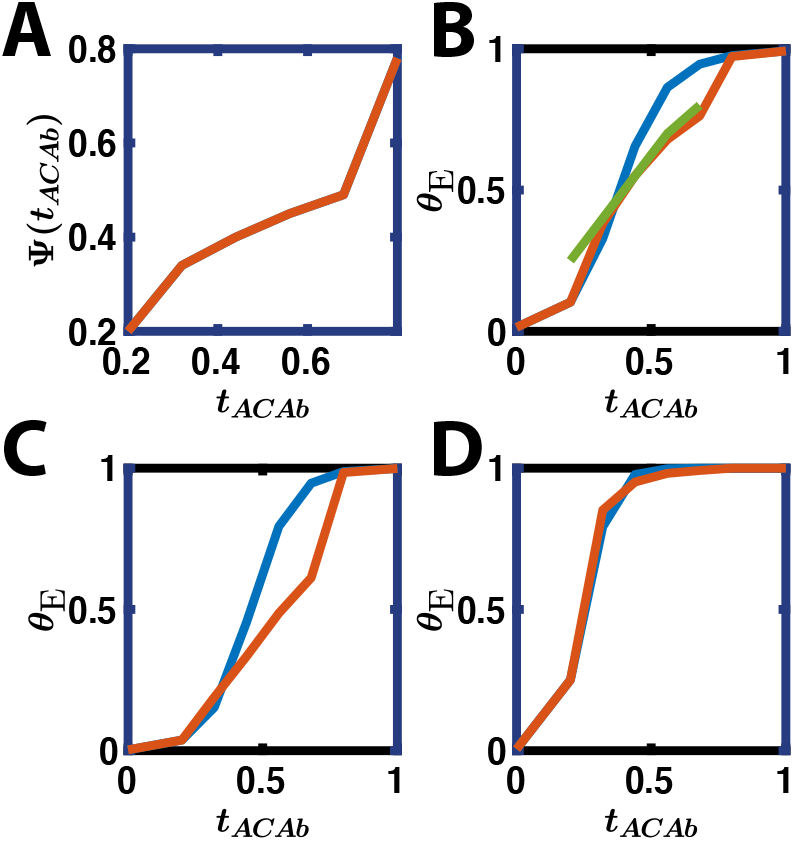
(A) Change in time Ψ obtained from fitting. (B) In blue, the mean over all the particles of the original EGF monotone sigmoidal function, *σ*(*t*). In orange, the mean over all the particles of the modified EGF monotone function, *σ*(Ψ(*t*)). In green, scaled *egl-17* expression measured in ***Milloz et al.*** (***2008***). (C-D) In blue, the original EGF monotone sigmoidal function, *σ*(*t*), in orange, the modified EGF monotone function, *σ*(Ψ(*t*)), for different parameter values

## Appendix 2

## Parameter estimation

We take advantage of sequential Monte Carlo ABC (ABC SMC) algorithm in ***Toni et al.*** (***2009***). This version of the ABC algorithm borrows ideas from importance sampling and sequential Monte Carlo, as the name suggests. The ABC SMC was first proposed by ***Sisson et al.*** (***2007***), and later corrected by ***Beaumont et al.*** (***2009***); ***Sisson et al.*** (***2009***); ***Toni et al.*** (***2009***). Various ABC SMC algorithms are proposed in the literature (see for example ***Beaumont et al.*** (***2009***); ***Moral et al.*** (***2012***); ***Sisson et al.*** (***2007***); ***Toni et al.*** (***2009***)), but here we focus on the fairly general ABC SMC algorithm by ***Toni et al.*** (***2009***). The ABC SMC sampler methodology approximates a sequence of probability distributions {*π*_*t*_}_0≤*t*≤*T*_ that satisfies the condition that *d*(**X**(θ)),**X**_0_)≤ε_*t*_

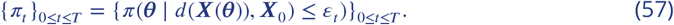

In order to produce these probability distributions, the algorithm starts by sampling parameter values ***θ***\* from a prior distribution *π*(***θ***). Then, the algorithm accepts *N* parameter values that satisfy *d*(**X**(θ*)),**X**_0_)≤ε_*1*_. It assigns a set of equal weights, 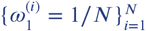, to each accepted parameter value 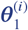, *i* = 1, … *,N*, in this context called particles.

Then the algorithm proceeds in a sequential manner. In each step *t*, a set of *N* particles 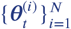 is generated. This is done by first sampling a particle ***θ***\*\* from the discrete distribution with support the finite set of particles generated in the previous step, 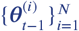, and probabilities given by the weights 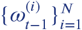. Then, the sample is perturbed using the Markov kernel function *K*_*t*_(***θ***\**, **θ***\*\*). These steps are repeated until N particles ***θ***\* that satisfy the conditions *π*(***θ***\*) *>* 0 and *d*(***X***(***θ***\*)*, **X***_0_)≤ε_*t*_ are found.

**Algorithm 1:**
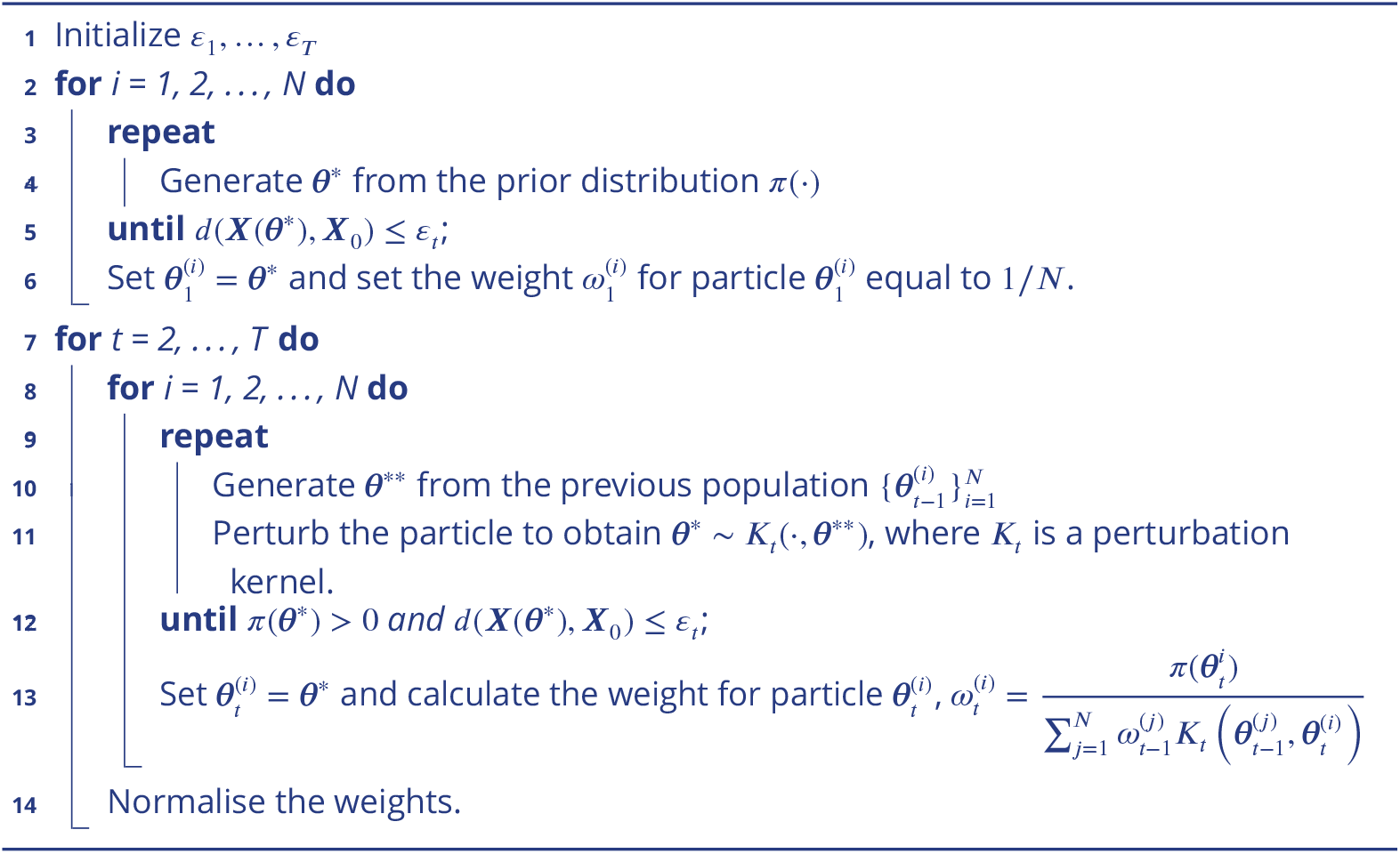
ABC SMC algorithm

For this implementation, we first need to study the dependence on the parameters of our model. We will study the range of the parameters, consider some constraints that the model and certain mutants in ***Table 1*** impose on the parameters, and provide the corresponding priors.

Secondly, we need to define a distance function *d* that will measure the similarity between the experimental data set and the corresponding simulated data set, i.e. the goodness of fit.

For the implementation of the ABC SMC method, we also need to define a decreasing sequence of thresholds {ε_*t*_}_*t*≥0_ that will determine the maximum distance between the experimental data and the simulated data in each step *t* of the algorithm. These, in turn, define the intermediate distribution 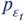 from which the algorithm samples at step *t*.

Also, we need to choose the number *N* of particles sampled from the distribution 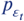 at each step *t*.

Finally, we need to provide the perturbation kernels 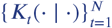 that will set the limits of exploration of the parameter space in each step.

We will discuss these points further in the following subsections. These steps are implemented in Matlab, and the code is provided in the following github repository: https://github.com/ecamacho90/VulvalDevelopment

## Constraints on the parameters

In Appendix 1 Table 1, we summarised the constraints that the transformation from the signal space to the control space imposes on the parameters. In the following subsections we will outline three other constraints imposed on the parameters by the model and the data in ***Table 1***.

## Constraint imposed by the sigmoidal function *χ*(*y*)

As mentioned above, we approximate the indicator function *H*(*y*) by a sigmoidal function that has the form

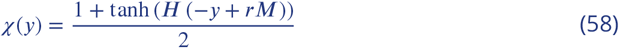

where we choose 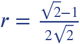.

Focusing on the deterministic system at first, the trajectory corresponding to the state of a VPC is moving on a flow characterised by the equilibria of the system:

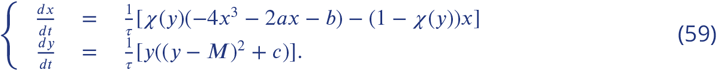

This is similar to Eq. 2 in the main text, but we have substituted the Heavyside function by the sigmoidal function. The control parameters *a, b, c* are functions of the signals as we explained before.

As studied above, the critical points of the system in Eq. 59 will have *y*-coordinates equal to 0, 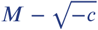 and 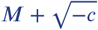 (these two latter ones only when *c* < 0).

The constraint that we introduce here is firstly due to the fact that *χ*(0) needs to be approximately equal to 1, so that the values of the *x*-coordinates of the critical points come only from the cusp equation. Secondly, 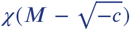 and 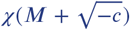 need to be approximately equal to zero so that there is only one possible value of the *x*-coordinate for these critical points and it must be equal to zero.

Consequently, we need to impose:

1. *χ*(0) ≈ 1 This condition can be translated into the condition 1 ≥ *χ*(0) ≥ 1− *ε* where *ε* is small. Substituting in Eq. 58 we get:

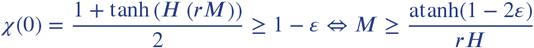 This constrains the smallest value that the parameter *M* > 0 can take. This minimum value of *M* will depend on how accurate we would like the function *χ*(*y*) to be or, in other words, how small *ε* is, as well as on the steepness *H* of the sigmoidal function. We choose *ε* = 10^−12^/2, and leave *H* > 0 as a parameter to be estimated. Therefore

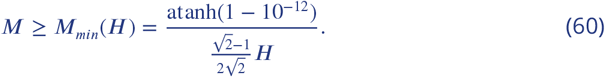 See Appendix 2 Figure 1 for a graphical explanation.

**Appendix 2 Figure 1.**
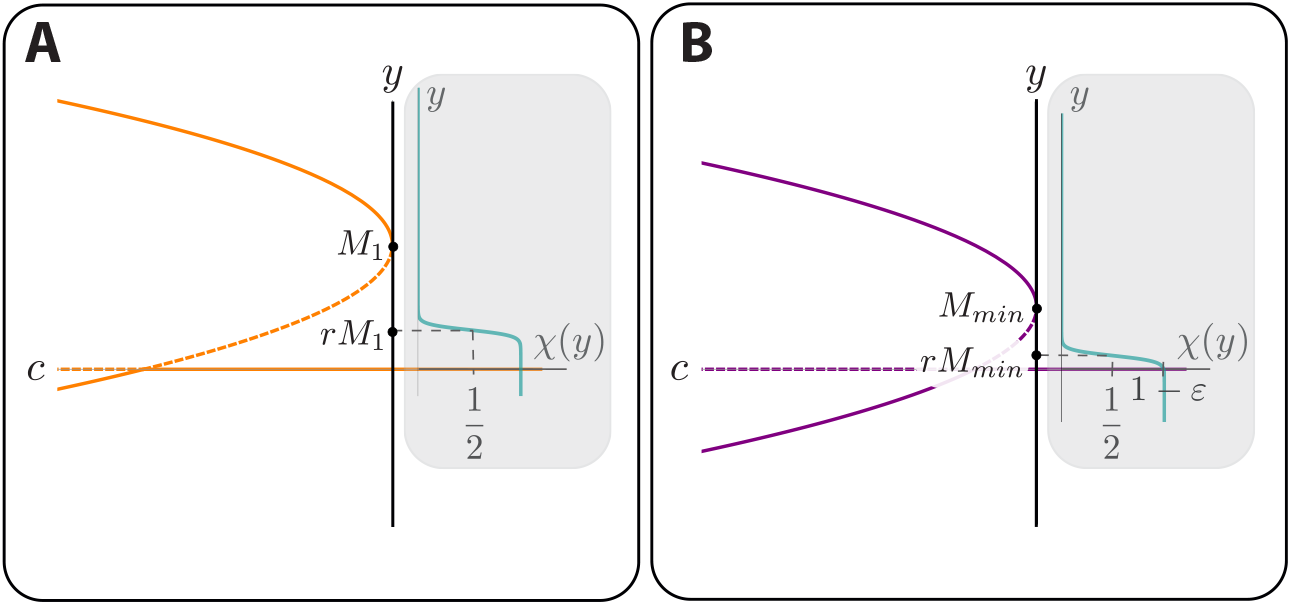
Bifurcation diagram of the *y*-coordinate with respect to the control parameter *c*, together with a plot of *χ*(*y*), for *M* = *M*_1_ (on the left plot (*A*)) and *M* = *M*_*min*_ (on the right plot (*B*)). Fixed a value of *H*, the value of *M* “*shifts*” the graph of *χ*(*y*) along the *y*-axis, by changing the value of *rM* where *χ*(*rM*) = 1/2. The minimum value *M*_*min*_ that *M* can take is such that *χ*(0) = 1 − *ε*, otherwise *χ*(0) would be “too far” from 1.

2. *χ*(*y*)≈0 when 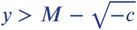 for all *M* and *c* allowed. This constraint is necessary so that the value of *χ*(*y*) evaluated at the critical points 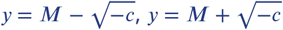 is approximately zero for all values of *M* and *c* allowed. We know that the sigmoidal function is monotonically decreasing, therefore 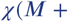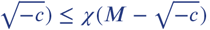. This means that we only need to impose 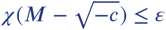, where *ε* takes the same value as before (*ε* = 10^−12^/2). To achieve that, we restrict the value of *c* to be bigger than a value *c*_*min*_, where *c*_*min*_ is such that 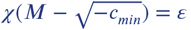. If *c* < *c*_*min*_ then 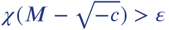, because *χ*(*y*) is monotonically decreasing. Therefore we should avoid values *c* < *c*_*min*_ (see Appendix 2 Figure 2). We should therefore impose:

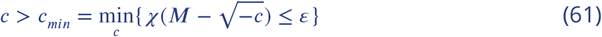

Now,

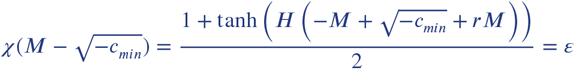

implies that

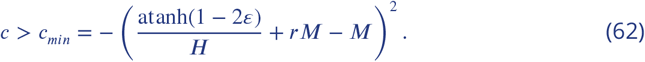 But in our model, the control parameter *c* is written as a function of the signals. In particular:

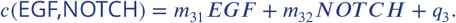 Considering the forms of the *θ*_E_ and functions (see Eq. 50, 51, 52, 53, 54) and the fact that *m*_31_*, m*_32_ *>* 0 (see Eq. 1) we know that *c*(EGF,NOTCH) ≥ *q*_3_. This leads us to the final condition:

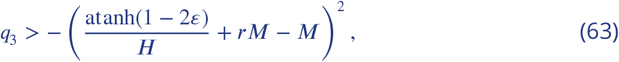

where 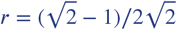 and *ε* = 10^−12^/2.

**Appendix 2 Figure 2.**
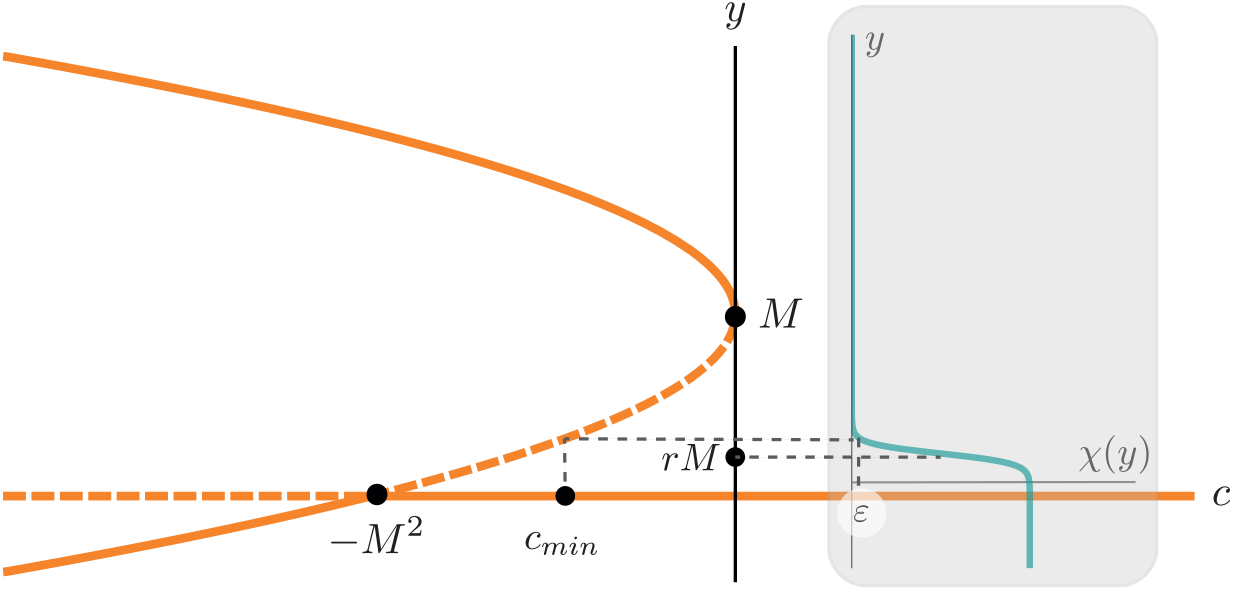
Bifurcation diagram of the *y* coordinate together with a plot of *χ*(*y*). *χ*(*y*) = 1/2 when *y* = *rM*. The value *c*_*min*_ is such that 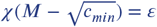.

## Constraint imposed by mutant (5) Notch null, 2xWT EGF

For this mutant, there is no Notch signal in any of the VPCs since they lack the Notch receptor. This means that, in our model, the *θ*_N_ coordinate is zero for all the cells at all times, i.e. *l* = 0.

In ***Table 1*** we can see that P5.p and P6.p adopt fate 1°. However, in our model, P5.p and P6.p could only adopt fate 1° if their trajectory escapes the basin of attraction of the attractor representing fate 3° when the signals are not present. This will only happen if the attractor corresponding to 3° fate bifurcates away for the values of the control parameters in Eq. 3, where *θ*_N_ =0 and *θ*_E_ is multiplied by two with respect to WT.

Since we know that *θ*_E_ increases in time for both cells, and that their maximum value is equal to *θ*_*E*2_ = 2*sγ* for P5.p and *θ*_*E*3_ = 2*s* for P6.p, this gives us the equations

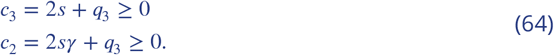

However, since *γ* ∈ (0, 1) and *s* > 0, we only need to impose 2*sγ* + *q*_3_ ≥ 0.

## Constraint imposed by mutant (6) Notch null, WT EGF

As before, there is no Notch signal in any of the cells and EGF is constant and equal to the WT signal. This means that, in our model, the *θ*_N_ coordinate is zero for all the cells at all times, i.e. *θ*_N_ = 0. As before, P6.p assumes primary fate (see ***Table 1***). With a similar reasoning as the one given before, this can only happen if the control parameters corresponding to P6.p satisfy the condition *c*_3_ = *s* + *q*_3_ ≥ 0.

## Ranges and priors

The range of values that each parameter can take is given on Appendix 2 Table 1. These ranges are imposed by the model definition and constraints in Appendix 1 Table 1.

**Appendix 2 Table 1.**
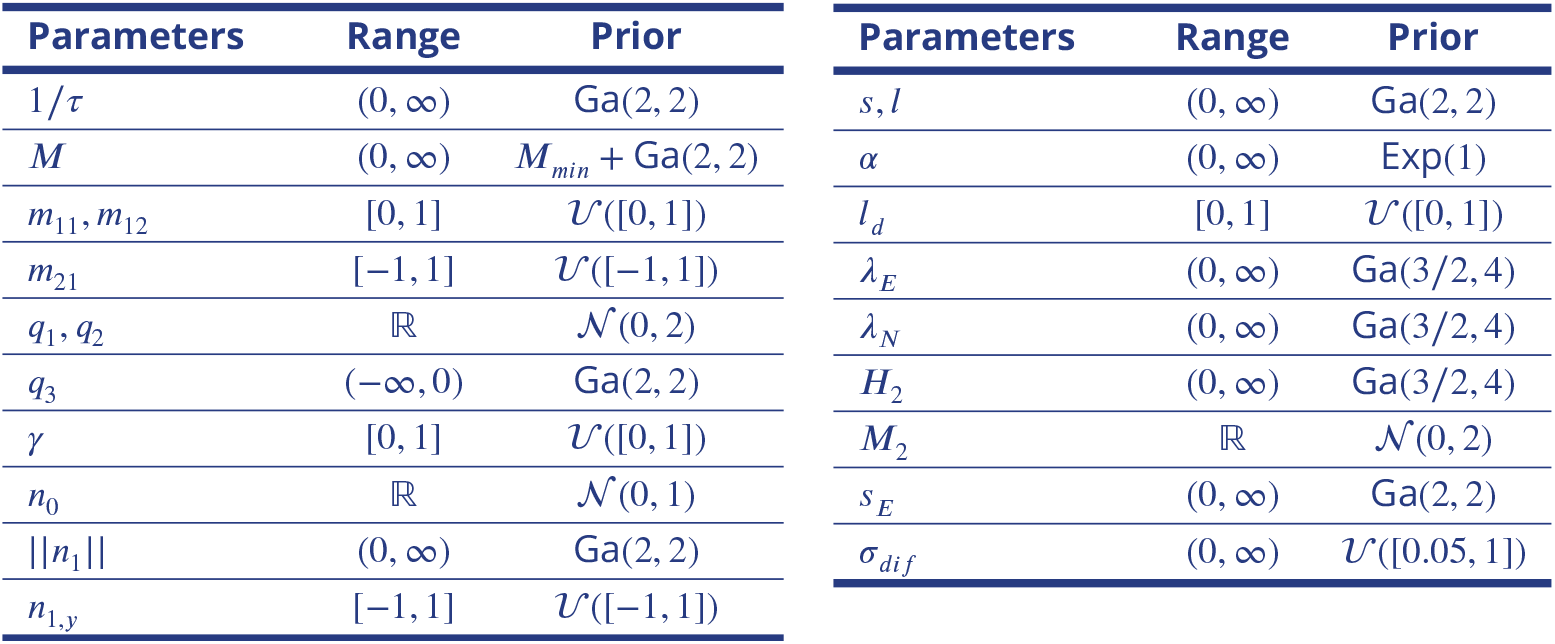
Tables of ranges and priors of the parameters.

In order to define the priors, we need to take into account their ranges and the constraints imposed on them by the data and by the model. The priors are chosen so that they are fairly non informative but still reflecting our knowledge about their ranges.

For bounded parameters *m*_11_*, m*_12_*, m*_21_*,γ*, *n*_1*,y*_ and *l*_*d*_ we assign uniform priors on their range.

Positive non-bounded parameters 1/*τ, H, H*_1_*, H*_2_, ∥*n*_1_∥ *, s, l* and *s*_*E*_ are assigned Gamma priors Ga(*κ, β*) where *κ* and *β* are the shape and scale of the distribution, respectively. Note that the probability density function of *U* ~ Ga(*κ, β*) is given by

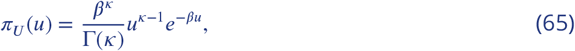

where Γ is the gamma function. We choose *κ* = 2, *β* = 2, and in this case Γ(*κ*) = Γ(2) = (2 − 1)! = 1, or *κ* = 3/2, *β* =4 for parameters where we would like to favor smaller values. For the positive and non-bounded parameter *M*, since it needs to be bigger than *M*_*min*_(*H*) (see Eq. 60), we impose *M* ~ *M*_*min*_(*H*)+ Ga(2, 2).

On the other hand, *M*_1_*, M*_2_*, q*_1_*, n*_0_ and *q*_2_ are assigned Normal priors 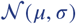 where *μ* is the mean of the distribution and *σ* is the standard deviation. Note that the probability density function of 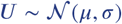 is given by

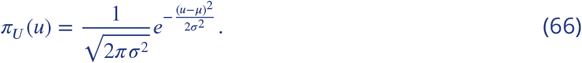

We choose *μ* =0 and *σ* =1 or 2.

For the parameter *α* we would like to favour smaller values, so we choose an exponential prior Exp(*λ*), where the probability density function of *U* ~ Exp(*λ*) is

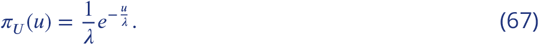

We choose *λ* = 1.

The parameter *q*_3_ is a negative non-bounded parameter, so we impose *q*_3_ ~ −Ga(2, 2).

And finally, since we would like to keep the noise level relatively low, we give *σ*_*dif*_ a flat prior on [0.05, 1], i.e. a uniform distribution 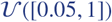.

Appendix 2 Table 1 contains a list of the priors for each parameter.

## Distance

In order to approximate the posterior distribution of the parameters, the ABC SMC algorithm computes a sequence of intermediate distributions that are obtained by comparing the simulated data and the experimental data. This comparison is made by means of a distance function *d* that measures the level of similarity between two data sets.

As explained in the main text, ***Table 1*** provides 171 data points or probabilities which correspond to the proportion of times that VPC *c* (*c* = 1, 2, 3 for P4.p, P5.p and P6.p respectively), adopted fate *f* (*f* = 1, 2, 3 for 1°, 2° and 3° respectively) in experiment *e* (*e* = 1, 2, … , 21 for the corresponding experiment in ***Table 1***). Each data point will be represented by *p*_*e,f,c*_.

Given a parameter vector ***θ***, we can simulate each data point using our model to obtain the corresponding 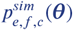.

If 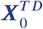 is the subset of data corresponding to experiments in the training data set (see ***Table 1***), we define the distance between 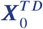 and the corresponding simulation ***X***^*TD*^ (***θ***) as

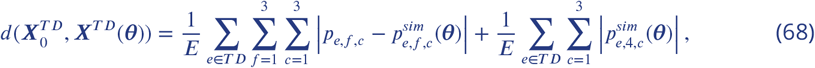

where *E* is the number of experiments in the training data set, *p*_*e,f,c*_ and 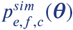 are the experimental and simulated probabilities of cell *c* becoming fate *f* in experimental condition *e*, respectively, and 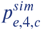 is the proportion of times that our model could not assign a fate to cell *c* when simulating experiment *e*.

## Sequence of thresholds and number of particles

The ABC SMC algorithm approximates the posterior distribution of the parameters by sequentially sampling from the distributions in Eq. 57. For this, one needs to choose a decreasing sequence of tolerances *ε*_1_ > *ε*_2_ > … *ε*_*T*_.

The most common approach is to choose *ε*_*t*+1_ to be the *α* quantile of the population of particles obtained at time *t* (***Beaumont et al., 2009***; ***Liepe et al., 2014***), taking into account a trade off between acceptance rate of the particles and the succesful convergence of the method (***Silk et al., 2012***). Here we use *α* = 0.3 quantile, so the method determines the value of the threshold *ε*_*t*_ at the beginning of the step *t* of the algorithm by sorting the distances obtained at the previous step and setting *ε*_*t*_ such that the 0.3% of the simulated data at step *t* −1 are below it. In our case, this sequence is given by *ε*_1_ =5 > … > 0.43 = *ε*_14_. We decided to stop at this threshold value because the resemblance between the simulated data and the training data was good and the variation across particles suggested that reducing the value further would result in overfitting.

An advantage of ABC SMC is that it can be easily parallelised since the *N* particles corresponding to a step *t* are sampled independently. With that in mind, taking into account the number of parameters to be fitted and the computational time of the simulations, we choose to sample *N* = 2 × 104 particles at each step *t* of the algorithm, and parallelise the computations, streaming computation into 100 jobs which compute 500 subsets of 40 particles each, significantly reducing the computation time of the algorithm.

## Perturbation kernel

The choice of perturbation kernel is also very important as it can speed up the convergence of the method (***Filippi et al., 2013***) if it samples from the regions of interest of parameter space.

For simplicity, let us denote by 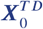 the set of training data. We denote the *i*-th particle obtained at step *t* −1 as ***θ***^(*i,t*−1)^, its corresponding weight as *w*^(*i,t*−1)^ and the data simulated with ***θ***^(*i,t*−1)^, for the same experiments as in 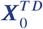, as ***X***(***θ***^(*i,t*−1))^. It holds that

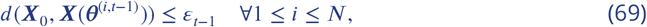

where *N* is the number of sampled particles at each step.

The particles at step *t* are computed by randomly choosing a particle ***θ***^(∗*,t*−1)^ from the previous population 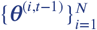 with weights 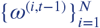, perturbing it with a chosen kernel *K*_*t*_(·|***θ***^(∗*,t*−1)^) and checking that the candidate parameter vector ***θ***\*\* obtained from that perturbation satisfies *d*(***X***_0_*, **X***(***θ***\*\*)) *< ε*_*t*_.

Many types of perturbation kernels *K*_*t*_(·|·) can be used. The most simple choice would be the component-wise perturbation kernel, in which the parameter vector ***θ*** = (*θ*_1_, … *, θ*_*m*_) (*m* being the number of parameters to be estimated) is perturbed component-wise. In other words, each parameter *θ*_*j*_ is perturbed according to a Gaussian distribution 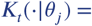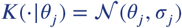 or Uniform distribution 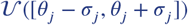. Note that, in this case, there is a kernel distribution for each component of the parameter vector and it is independent of the other components. Moreover, the kernel distributions are the same for all *t*. As a result, these kernels are not very efficient since they do not take into account the fact that some parameters can have a certain correlation.

Following the study in ***Filippi et al.*** (***2013***) we decide to implement the multivariate normal kernel with optimal covariance matrix (OLCM). This perturbation kernel considers a multivariate normal distribution around each particle (assessing the correlation between the parameters) and, moreover, the covariance matrix will differ from particle to particle, taking into account the structure of the *good* particles sampled. Let us define the following set

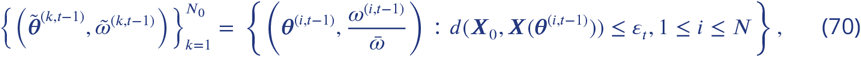

where 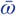 is a normalising constant such that 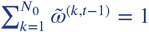. The set in Eq. (70) is the set of particles of the population obtained at time *t* −1 for which the simulated data is closer to the experimental data than the current threshold *ε*_*t*_, i.e. the set of *good* particles at time *t* − 1. The weights 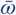 are obtained by normalising this subset of *N*_0_ particles.

In this approach, the perturbation kernel follows a normal distribution, 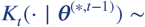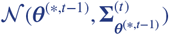, which is centered at the particle that will be perturbed ***θ***^(∗*,t*−1)^ with covariance matrix also dependent on the particle value. ***Filippi et al.*** (***2013***) propose the optimal covariance matrix 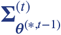 to be equal to the covariance of the set of particles from step *t* −1 whose distance is smaller than the current threshold *ε*_*t*_ (i.e. particles in the set described in Eq. 70) plus a bias term related to the discrepancy between the mean of the particles in such population and the particle of interest ***θ***^(∗*,t*−1)^:

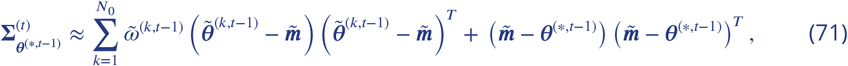

where 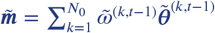 is the mean of the population of particles described in Eq. 70. We therefore implement these multivariate normal perturbation kernels 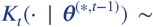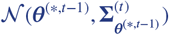 with optimal covariance matrix given in Eq. 71.

## Exploring the single-cell fate map

We also studied how the data constrained the single-cell fate map in the signal space, given by the map between the control space and the biological signals.

As we described before, the non-vulval to vulval transition is determined by the line *r*_*c*_ = {(*θ*_E_*, θ*_N_) : *θ*_N_ = tan(*ω*_*c*_)*θ*_E_ − *q*_3_/*m*_32_*l*} where *ω*_*c*_ = −*m*_31_*s*/*m*_32_*l*, that is, when the control parameter *c* is equal to zero; while the primary to secondary fate transition is determined by the cusp, which mid-line is determined by the line *r*_*b*_ = {(*θ*_E_*, θ*_N_): *θ*_N_ = tan(*ω*_*b*_)*θ*_E_ − *q*_2_/*m*_22_*l*} where *ω*_*b*_ = −*m*_21_*s*/*m*_22_*l*, that is, when the control parameter *b* is equal to zero (see Eq. 3) (see Appendix 2 Figure 3A). It is important to note that *θ*_E_ and *θ*_N_ have arbitrary units. And therefore, understanding the positions of *r*_*c*_ and *r*_*b*_ can give us information about how signal perturbations can affect the differentiation of a single cell and, in turn, vulval patterning.

Interestingly, even though we did not impose any constraints on the steepness of the lines *r*_*c*_ and *r_b_*, the results of fitting suggest that they are strongly constrained by the data (Appendix 2 Figure 3A). We find that the angle that forms the line *r*_*c*_ with the *θ*_E_ axis, *ω*_*c*_, is always between *π*/2 and 3*π*/4, converging to a mean steepness of −2.16± 0.5 after the 14 steps of ABC SMC (Appendix 2 Figure 3).

Similarly, we observe that the steepness of the *r_b_* line is also strongly constrained by the data, converging to a positive value of 1.66 ± 0.65 after the 14 steps of ABC SMC (Appendix 2 Figure 3A). Moreover, we observe that the two lines *r*_*c*_ and *r*_*b*_ normally intersect in an angle of close to *π*/2.

The position of the cusp point within the *r*_*b*_ line, however, is more variable, as shown in Appendix 2 Figure 3B. We wondered whether this variability is due to the degree of freedom given by the choice of the scaling parameters *s* and *l*. However, we see that there is no correlation between these two parameters and the position of the cusp point (Appendix 2 Figure 3B).

**Appendix 2 Figure 3.**
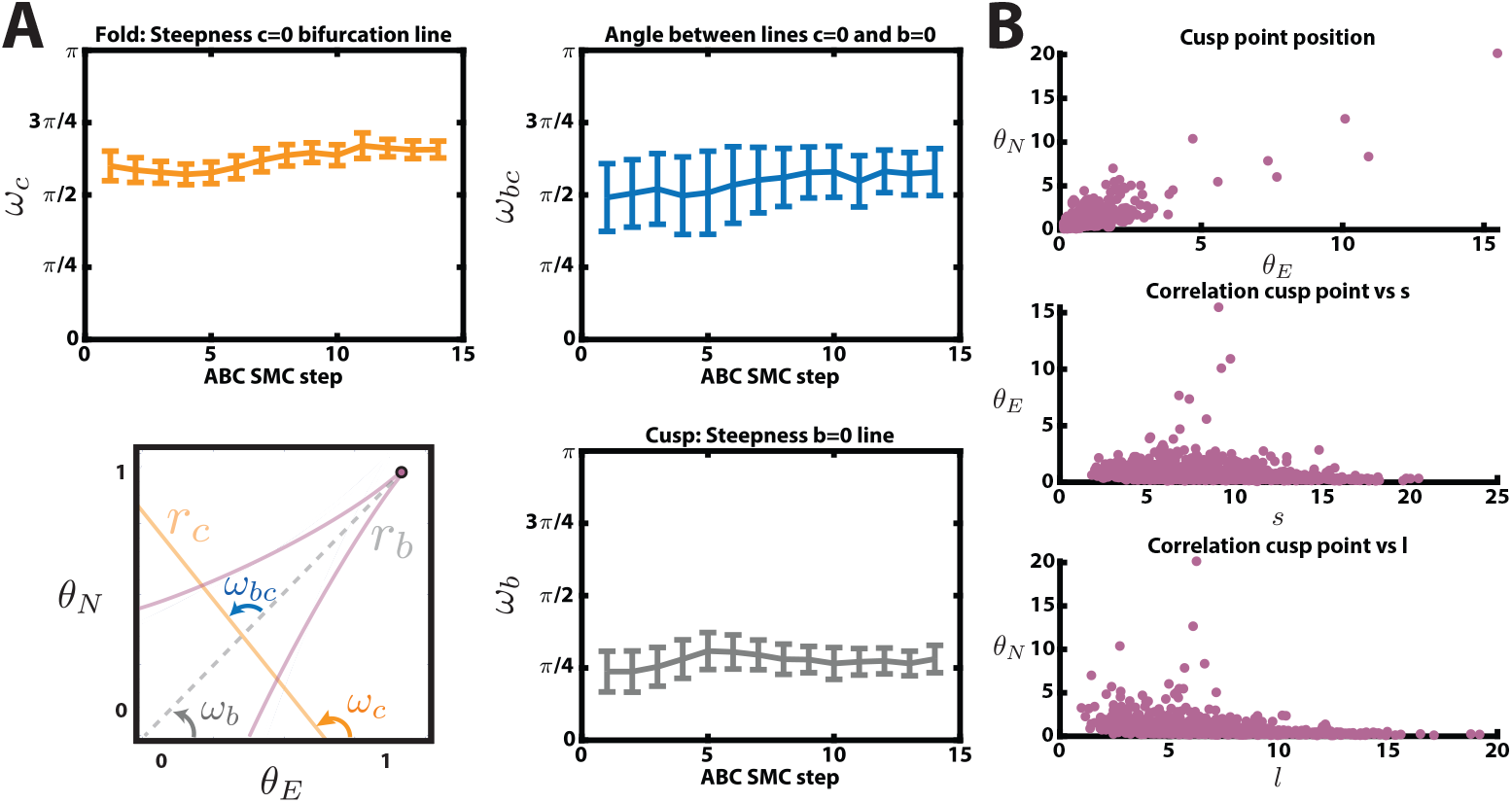
Data-fitted single-cell fate map. (A) Evolution of the steepness of the fold line *r*_*c*_ (orange), the steepness of the cusp mid-line *r*_*b*_ (gray) and the angle between them (blue) over the steps of the ABC algorithm. (B) Distribution of the positions of the cusp point for each particle obtained in the last step of the algorithm. Top: Coordinates of such cusp points in the (*θ*_E_*, θ*_N_) space. Middle: Correlation between EGF scaling parameter *s* and position of the cusp point on the EGF-axis. Bottom: Correlation between Notch scaling parameter *l* and position of the cusp point on the NOTCH-axis.

---

**Figure 4–Figure supplement 1.**
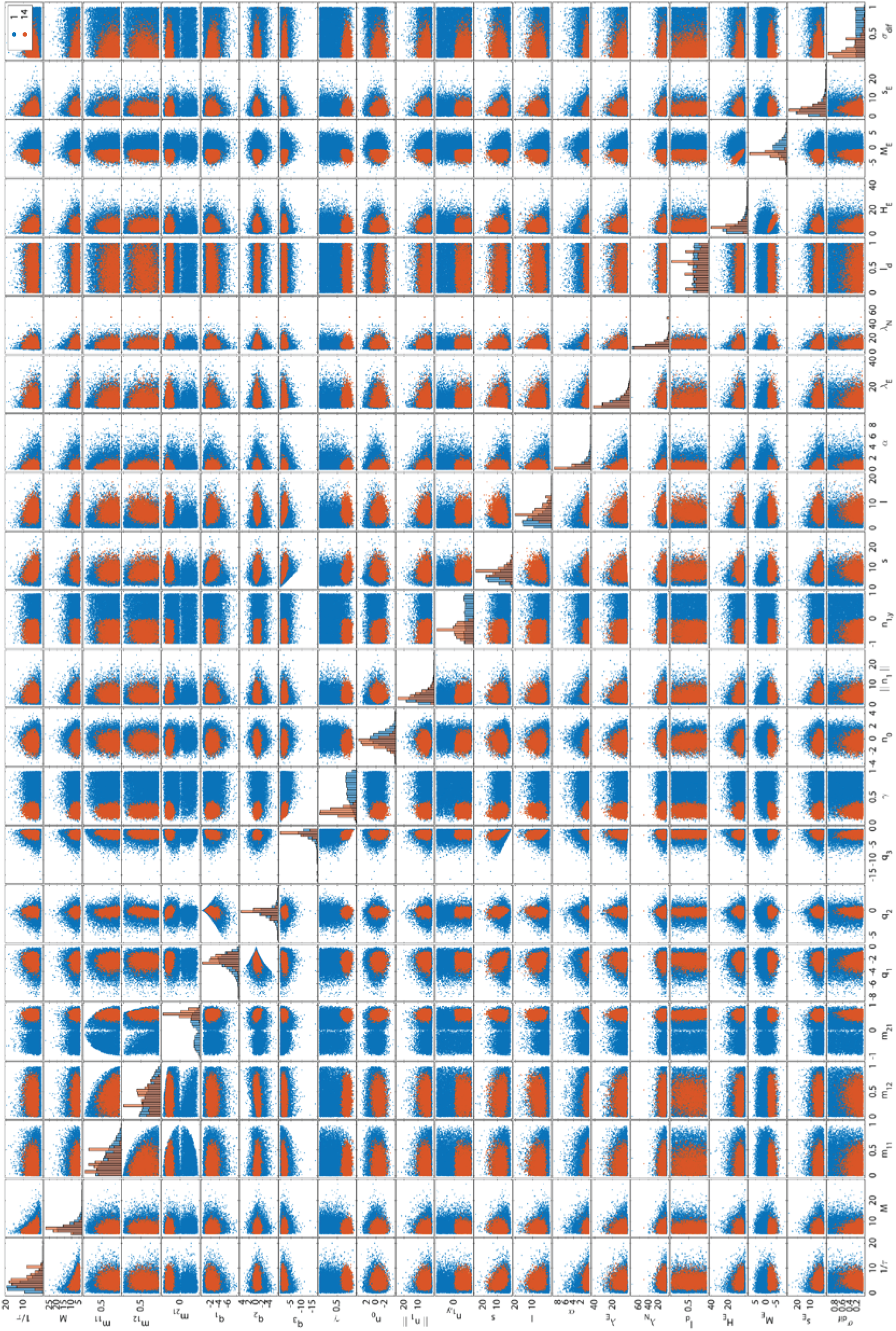
Histograms and two dimensional scatter plots of the *N* = 2 × 10^4^ particles sampled from the approximated posterior distribution given the training data in **Table 1** at the first (blue) and last (orange) steps of the algorithm.

